# PLMView: collaborative protein language model representations for fast and scalable specialized protein function inference

**DOI:** 10.64898/2026.07.31.742035

**Authors:** Vinh-Son Pho, Alessandro Natale Bianchi, Matteo Scarsini, Chris Bowler, Alessandra Carbone

## Abstract

The functional classification of protein sequences remains a major bottleneck in biology. Although protein language model (PLM)-based approaches have substantially improved broad protein function prediction, most protein sequences still lack precise annotation at the level of specialized functions—the fine-grained molecular roles that define specificity within protein families. We present PLMView, an unsupervised framework for fine-grained protein function classification directly from sequence. PLMView reframes protein function inference as a relational problem: instead of embedding sequences in isolation, it positions them within a collaborative functional space defined by comparisons with PLM embeddings of anchor sequences, thereby capturing subtle sequence–function relationships. Without requiring labeled data, family-specific training, or PLM fine-tuning, PLMView accurately distinguishes specialized functions among homologous proteins and highlights residues likely to determine functional specificity. The method achieves high precision while remaining computationally efficient, classifying approximately 10,000 sequences with 1,000 anchors in under 40 minutes; compared with pooled-embedding approaches and, in challenging cases, Sequence Similarity Networks, PLMView provides finer and more biologically coherent functional resolution, while achieving more than 10-fold speed-up over SSN reconstruction on datasets of this scale. Applications to thioredoxins, visual opsins, and *Tara* Oceans environmental diatom cold-shock proteins show that PLMView can move from interpretable residue-level determinants in well-studied protein families to large-scale environmental functional discovery, linking molecular specialization to ecological distribution and transcriptional deployment across the global ocean.

## INTRODUCTION

Proteins are the molecular engines of life. They drive nearly every cellular process, from catalysis and signaling to structural maintenance and environmental adaptation. Understanding how protein sequences encode their myriad of functions remains one of the central challenges in biology. Despite decades of biochemical and structural characterization, our view of protein function is still partial: for most known sequences, their precise activity, interacting partners, and mechanisms remain uncharacterized. This gap limits our ability to interpret the rapidly expanding universe of genomic data and constrains the discovery of new biological principles.

Global sequencing initiatives, such as *Tara* Oceans, are delivering an unprecedented catalog of protein diversity across every branch of life^1–5^. However, the rate at which new sequences are produced vastly exceeds our capacity to characterize their functions experimentally. Computational annotation has therefore become indispensable, but it remains predominantly effective at a relatively coarse level, assigning proteins to broad families, domains, or generic biochemical activities. Such annotations are valuable, yet they frequently conceal the functional diversity that exists within homologous protein families.

This limitation reflects a fundamental resolution problem in protein function inference. Members of the same family may share a common fold, conserved motifs, and substantial sequence similarity while performing distinct specialized functions^6^. These fine-grained molecular roles can involve differences in substrate selectivity, interaction partners, regulatory mechanisms, subcellular localization, or pathway integration. Importantly, such specialization is often driven not by large structural changes or the appearance of new motifs, but by sparse and context-dependent substitutions distributed across the sequence^7^. Consequently, sequence similarity, domain architecture, and motif detection can identify broad molecular roles while failing to distinguish the biological specificities of closely related homologues.

The challenge is especially acute in large protein families, in which thousands of sequences may share a conserved structural framework yet occupy distinct functional niches. Conventional homology-based annotation tends to collapse this diversity into generic labels, obscuring biologically meaningful subdivisions. For enzymes, this limitation is illustrated by Enzyme Commission classifications, which describe overall reaction chemistry but do not always capture the molecular determinants of substrate selectivity, catalytic specialization, or regulatory context^8^. Comparable limitations apply to signaling, regulatory, and interaction-mediated proteins^9–12^. A more informative conception of functional annotation must therefore go beyond assigning sequences to broad classes: it should also connect inferred functional partitions to the sequence-level features that implement them. In this view, fine-grained classification is not complete unless the separation of functional groups can be traced to residues or regions whose variation encodes specialization.

Protein language models (PLMs), trained on millions of natural sequences, have transformed the computational analysis of proteins by learning representations that reflect biochemical, structural, and evolutionary constraints. PLM-based approaches have substantially improved broad protein function prediction and annotation transfer^13–19^. However, many existing methods compress residue-level representations into a single pooled sequence vector. Although effective for detecting broad relationships, this compression can obscure subtle, position-specific signals associated with specialized functions.

Recent studies have begun to reveal the biological organization contained within PLM representations. Sparse autoencoders applied to PLM embeddings can recover residue-level biochemical and structural features^20,21^, while local and global comparisons of residue embeddings correlate strongly with structural similarity^22–24^. These observations suggest that PLM embeddings encode information relevant not only to overall protein identity, but also to the contextual contributions of individual residues. At the same time, several approaches have used PLM representations to transfer or predict labels from established functional vocabularies, including Gene Ontology, Enzyme Commission, and KEGG Orthology ^14,16–18,25,26^. These methods considerably extend annotation coverage, including for proteins that are difficult to characterize by conventional homology-based approaches.

Nevertheless, label-driven approaches remain constrained by the completeness, granularity, and biases of the annotations on which they are trained. Experimentally supported annotations cover only a small fraction of known proteins and are strongly concentrated in model organisms, canonical pathways, and well-studied families^26,27^. Supervised predictors trained on GO, EC, or KEGG labels can therefore project proteins into existing functional categories, but they are less suited to identifying previously unrecognized subdivisions when the relevant labels are absent, incomplete, or too coarse. This is particularly problematic for specialized functions: closely related homologues may share the same broad annotation while differing in substrate specificity, regulatory control, interaction partners, or ecological deployment.

More generally, functional specialization is inherently relational. It cannot always be inferred from a sequence considered in isolation, because specialization is defined by the similarities and differences between that sequence and its homologues^28^. Current PLM-based methods, however, treat proteins as independent representations or transfer labels through pairwise similarity to annotated examples, without constructing a functional space in which homologous sequences collaboratively define one another’s positions. As a result, subtle sequence variations that distinguish specialized functions remain difficult to resolve, particularly within closely related subfamilies.

Here, we introduce PLMView, an unsupervised collaborative framework for fine-grained protein function inference. PLMView reframes functional classification as a relational problem: rather than representing each protein through an isolated pooled embedding, it positions sequences within an anchor-based, multi-view functional space defined by comparisons among homologous PLM representations. Each anchor provides a complementary functional perspective, and the collective pattern of agreement and divergence across anchors determines the position of a sequence in this space. This representation is designed to amplify subtle sequence–function relationships that may remain hidden in conventional sequence or embedding comparisons.

Importantly, PLMView uses this collaborative space not only to organize proteins into functionally coherent groups, but also to identify the residue-level determinants underlying their separation, namely sequence positions whose contextual variation is most strongly associated with functional specialization between the inferred groups. By decomposing embedding comparisons into positional contributions and contrasting these contributions across hierarchical functional partitions, PLMView links inferred classes to residues whose contextual signals drive functional divergence. It therefore moves beyond terminal label assignment toward a mechanistically interpretable representation of specialization. The framework requires neither labeled training data, family-specific model training, structural information, nor PLM fine-tuning, and can consequently be applied to protein families for which formal functional annotations are sparse or unavailable.

In this work, we evaluate PLMView through a progression from mechanistically interpretable protein families to environmental functional discovery. We first apply the method to protein families in which functional divergence is partly understood. Thioredoxins provide an experimentally grounded interpretability case, in which distributed residue-level determinants are linked to redox-related substrate specificity and overlap with positions capable of rewiring thioredoxin recognition. Visual opsins provide a complementary phenotype-linked benchmark, where functional divergence is associated with spectral tuning. Together, these examples establish that PLMView recovers biologically interpretable functional partitions and the residues that support them. We then extend the same framework to environmental diatom metagenomic and meta-transcriptomic sequences from the *Tara* Oceans dataset. Diatoms are a widely distributed group of eukaryotic phytoplankton that are particularly abundant in polar regions^29^. Using cold-shock DNA-binding proteins as a case study, we demonstrate the framework’s potential for environmental functional discovery. Together, these analyses test whether PLMView can resolve fine-grained functional diversity, identify the molecular determinants supporting that diversity, and extend interpretable function inference to large environmental sequence datasets. By scaling efficiently to large protein families, PLMView opens the way to the systematic exploration of functional determinants across the protein universe.

## RESULTS

### Collaborative PLM representations for fine-grained and interpretable function inference

PLMView is an unsupervised framework for fine-grained and interpretable function inference within homologous protein families. We first provide an intuitive introduction to its collaborative principle, the construction of its functional space, and its residue-level interpretation, before outlining the biological applications it enables, from well-characterized protein families to large environmental sequence collections (Fig. 1).

**Figure 1:**
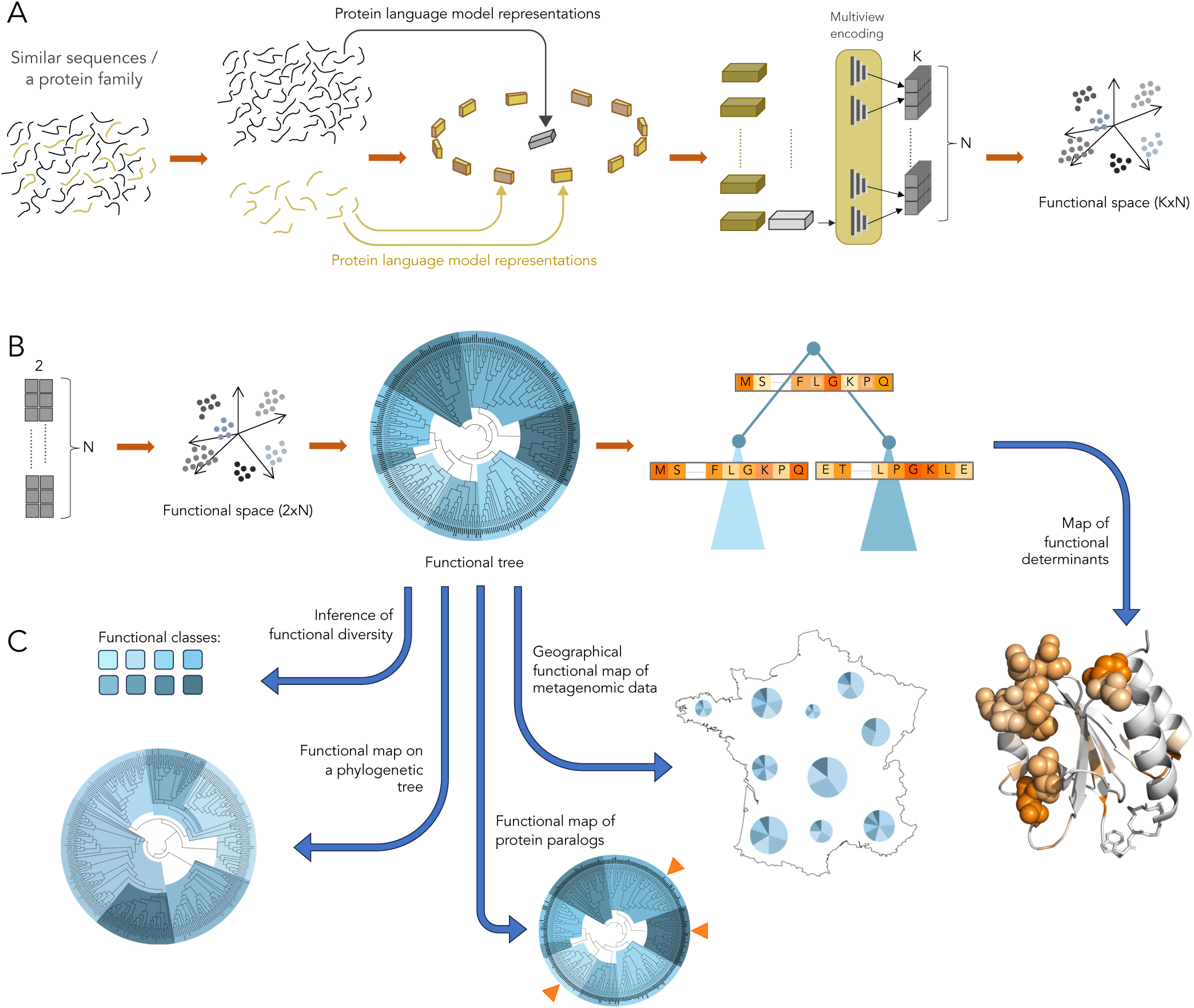
Overview of the PLMView method and its applications. **A.** Starting from an initial set of homologous sequences, PLMView selects a subset of anchor sequences (gold). All sequences are transformed into PLM representations, and each sequence to be classified (grey) is compared pairwise against all anchors. The anchors collaboratively evaluate the functional relationship of each sequence within the dataset. For each anchor–sequence pair, PLMView computes *K* comparison scores, so that each sequence is encoded by a vector of dimension *K × N*, where *N* is the number of anchors. The resulting *K × N*-dimensional functional space can then be clustered to identify functionally coherent groups. **B.** In the present implementation, PLMView uses two complementary embedding-comparison measures (*K* = 2), yielding a functional space of dimension 2 *× N*. Clustering within this space enables the construction of a functional tree. PLMView further decomposes the position of each sequence in this tree into residue-level contributions. A tree-split-based analysis of these residue-level weights provides a residue-resolved importance score for each cluster separation, defined as the difference in positional importance between sister subclusters forming the parent cluster. Thus, for each residue in a classified sequence, PLMView extracts a weight reflecting its contribution to the sequence classification. **C.** PLMView provides a global view of functional diversity within a protein family through its distinct subtrees in the functional tree, while also offering a comparative framework to analyze the functional relationships among two or more sequences. These capabilities support several applications: mapping inferred functional labels onto phylogenetic trees to compare functional and evolutionary organization; distinguishing specialized functions among paralogous sequences; projecting functional classes onto metagenomic and/or meta-transcriptomic data from independent geographical locations to generate geographical functional maps of protein families; and mapping residue-level weights for individual sequences onto predicted or experimentally determined protein structures.

### Collaborative PLM representations define a relational space for protein specialization

PLMView introduces a collaborative use of PLM representations for functional inference. A protein is not characterized by an embedding considered independently, but by the relationships that its residue-level embedding establishes with a panel of homologous reference sequences, termed anchors. We use “collaborative” in this precise representational sense: the coordinates assigned to a protein are jointly determined by comparisons with the complete anchor panel rather than being intrinsic to an independently considered embedding. Each anchor provides a complementary reference view, and the combined responses form a relational fingerprint of the protein within its homologous family (Fig. 1A).

Proteins are therefore compared through the similarity of their relational fingerprints rather than through a single global sequence or embedding score. Because PLMs represent each residue according to its sequence environment, these comparisons retain signals associated with biochemical, structural, and evolutionary constraints. Different anchors interrogate complementary aspects of this contextual representation: responses shared across the anchor panel capture properties common to the family, whereas differential responses reveal subtle sequence variation associated with functional specialization. Homologues with similar relational fingerprints consequently occupy nearby regions of the functional space, allowing the relational organization of the family to amplify distinctions that may remain weak when sequences or embeddings are examined individually or through pooled representations. This residue-resolved organization therefore enables both fine-grained functional discrimination and the subsequent identification of sequence positions associated with functional specialization.

The collaborative principle has a conceptual precedent in ProfileView^28^, where multiple profile HMMs were combined to resolve functional diversity. Because each profile requires a multiple-sequence alignment and a separate model-building step, this formulation incurs substantial pre-processing costs and scales less readily to large sequence collections. PLMView transfers collaborative inference to contextual PLM embeddings generated directly from individual proteins. By comparing the contextual residue-level embeddings of both anchors and queries, PLMView defines a relational space that carries richer contextual information than one constructed from sequence-to-profile comparisons. This representation preserves residue-specific sequence context and can be decomposed to identify the positions supporting individual functional separations. Thus, ProfileView provides the conceptual precedent, whereas scalable collaboration among residue-level PLM representations is the defining innovation of PLMView.

### Anchor comparisons generate the PLMView functional space

PLMView implements the collaborative principle by representing each protein through its relationships to a panel of anchor sequences. Starting from a set of homologs, the method selects *N* anchors and transforms both anchors and query sequences into residue-level PLM embeddings (Fig. 1A). For a protein of length *L*, the resulting representation is an *L × H* matrix, where each residue is associated with an *H*-dimensional contextual embedding (Fig. 2A). PLMView preserves this positional organization rather than compressing the protein into a single pooled vector.

**Figure 2:**
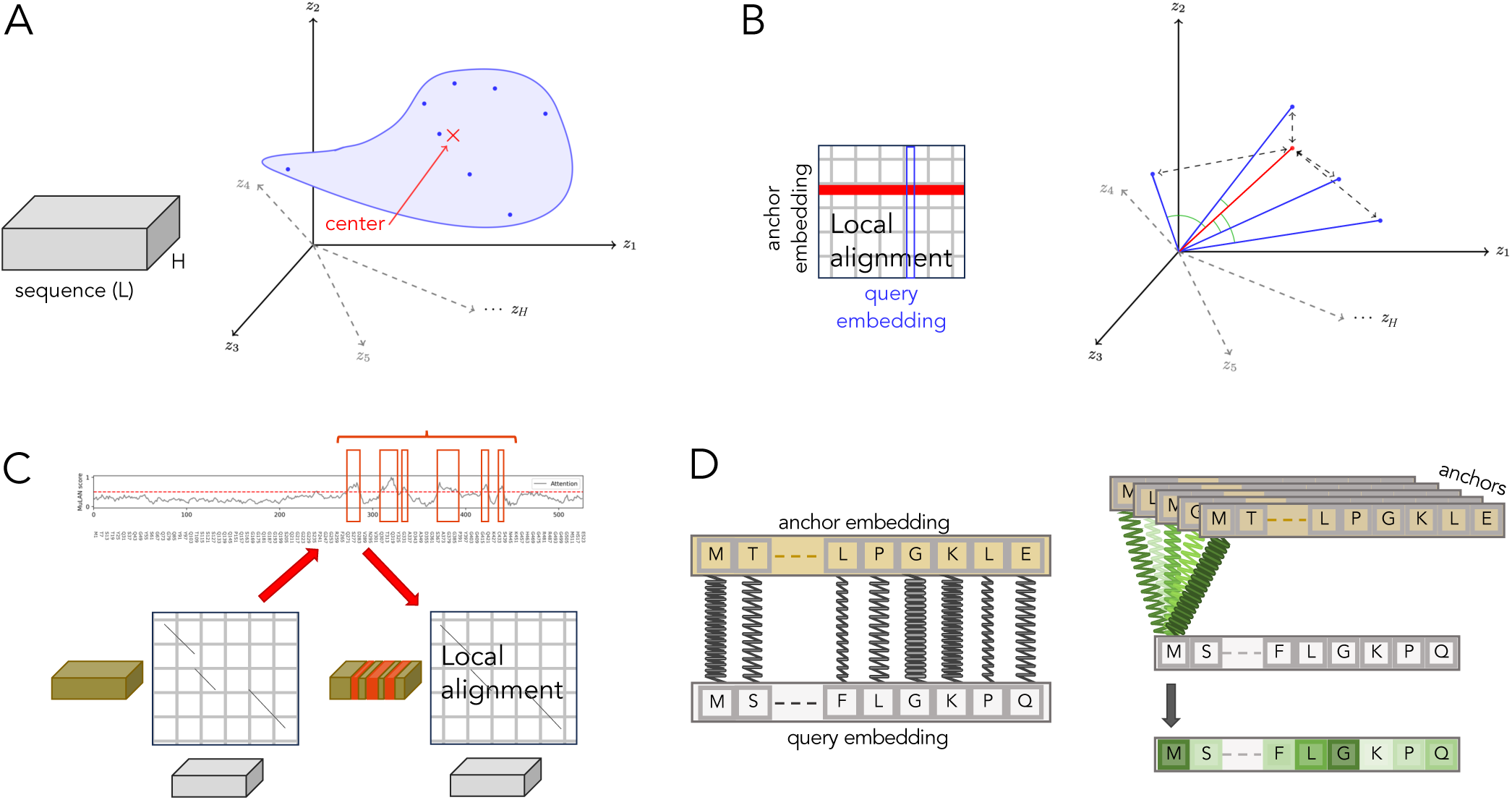
PLM embeddings, alignments, and residue weights. **A.** The space of a single embedding. An embedding of dimension *L × H* (Fig. 1), where *L* is the length of the sequence and *H* the dimension of the PLM model used to embed, is visualised in a multidimensional space of *H* dimensions as a set of points, one for each amino acid in the sequence. A red cross indicates the center of the cloud. **B.** The space of embedding pairs. In the alignment matrix representation, local alignment between anchor and query embeddings maps each position in the anchor sequence (red row) to positions in query sequence (blue column). The same local alignment can also be represented in an *H*-dimensional embedding-pair space (right), where anchor residue embeddings (red vector) are compared with query residue embeddings (blue vectors). Pairwise relationships are quantified using cosine similarity (green angles) and RBF similarity (dashed arrows). **C.** MuLAN profile analysis of an anchor sequence reveals regions of high functional importance (red rectangles); MuLAN is applied only to the anchor, not to the query sequence. **D.** Comparison between anchor and query embeddings determines the contribution (weight) of each residue to the positioning of a sequence in the multidimensional functional space. Using a mass–spring analogy, darker/thicker (lighter/thinner) springs represent higher (lower) similarity between residues in the anchor and the query embeddings, effectively pulling the query embedding toward the corresponding anchor-defined dimensions (left). Each query residue is assigned a weight based on its similarities to the corresponding aligned residues across the anchor ensemble (right).

Each query sequence is then compared with every anchor at the level of their residue embeddings. For each anchor–query pair, PLMView computes *K* complementary similarity scores, so that the complete panel of *N* anchors defines a *K × N*-dimensional representation of the query (Fig. 1A,B). In the current implementation, *K* = 2: a radial basis function (RBF) similarity derived from Euclidean distances between residue embeddings and a cosine similarity measuring their directional agreement. The two measures retain complementary aspects of the embedding geometry and provide two coordinates for each anchor (Fig. 2B).

To focus these comparisons on positions likely to carry functional information, PLMView incorporates MuLAN^30^, a residue-level importance estimator operating directly on single-sequence PLM embeddings. For each anchor, MuLAN produces a positional profile assigning larger weights to residues and regions predicted to be informative for protein function and smaller weights to less discriminative positions (Fig. 2C). Importantly, MuLAN-derived weights are computed only for the anchor sequence; no corresponding weights are assigned to the query sequence. During each anchor–query comparison, the anchor weights modulate the contribution of similarities involving the aligned query residues, allowing informative anchor positions to contribute more strongly to the final scores. These profiles may highlight positions located in disordered regions while down-weighting structurally conserved but functionally neutral regions. MuLAN therefore does not define functional classes directly but modulates the residue-level evidence used to compute the collaborative coordinates, thereby improving downstream interpretability.

The weighted residue similarities are aligned and aggregated to obtain the two sequence-level scores associated with each anchor. For a query sequence *B* and a panel of *N* anchors *A*_1_*,…, A_N_*, the two similarity scores obtained for each anchor are concatenated to define the relational representation

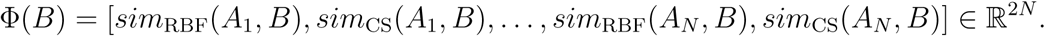

where *sim*_RBF_ and *sim*_CS_ denote the RBF-based and the cosine-based similarity scores, respectively. These complementary scores capture distance-based and directional relationships between the residue embeddings of each query–anchor pair (Fig. 2B).

Thus, the coordinates of a protein encode its pattern of relationships to the complete anchor ensemble rather than an intrinsic property of its embedding. The dimensionality of this representation is independent of protein length. Anchors displaying stronger functional similarity to the query contribute higher scores, whereas more distant anchors produce lower or distinct responses. The protein is thus positioned relative to the functional diversity of its homologous family rather than represented by an absolute embedding in isolation.

The collection of query vectors defines the PLMView functional space of a protein family (Fig. 3). Its dimensions correspond to complementary anchor-derived perspectives, and proteins with similar relational profiles occupy nearby regions. Signals shared across anchors reflect family-level properties, whereas differences among selected coordinates reveal specialization within closely related groups. Divisive hierarchical clustering of the functional space produces clusters representing the organization of the full set of sequences (Fig. 1B and Fig. 3A): higher-level partitions capture broad functional classes, whereas deeper partitions resolve progressively finer subfunctional classes (orange panels in Fig. 3B). Experimental annotations and quantitative phenotypes are not used in clustering but can subsequently be mapped onto clusters to evaluate or interpret the inferred organization (blue panels in Fig. 3B).

**Figure 3:**
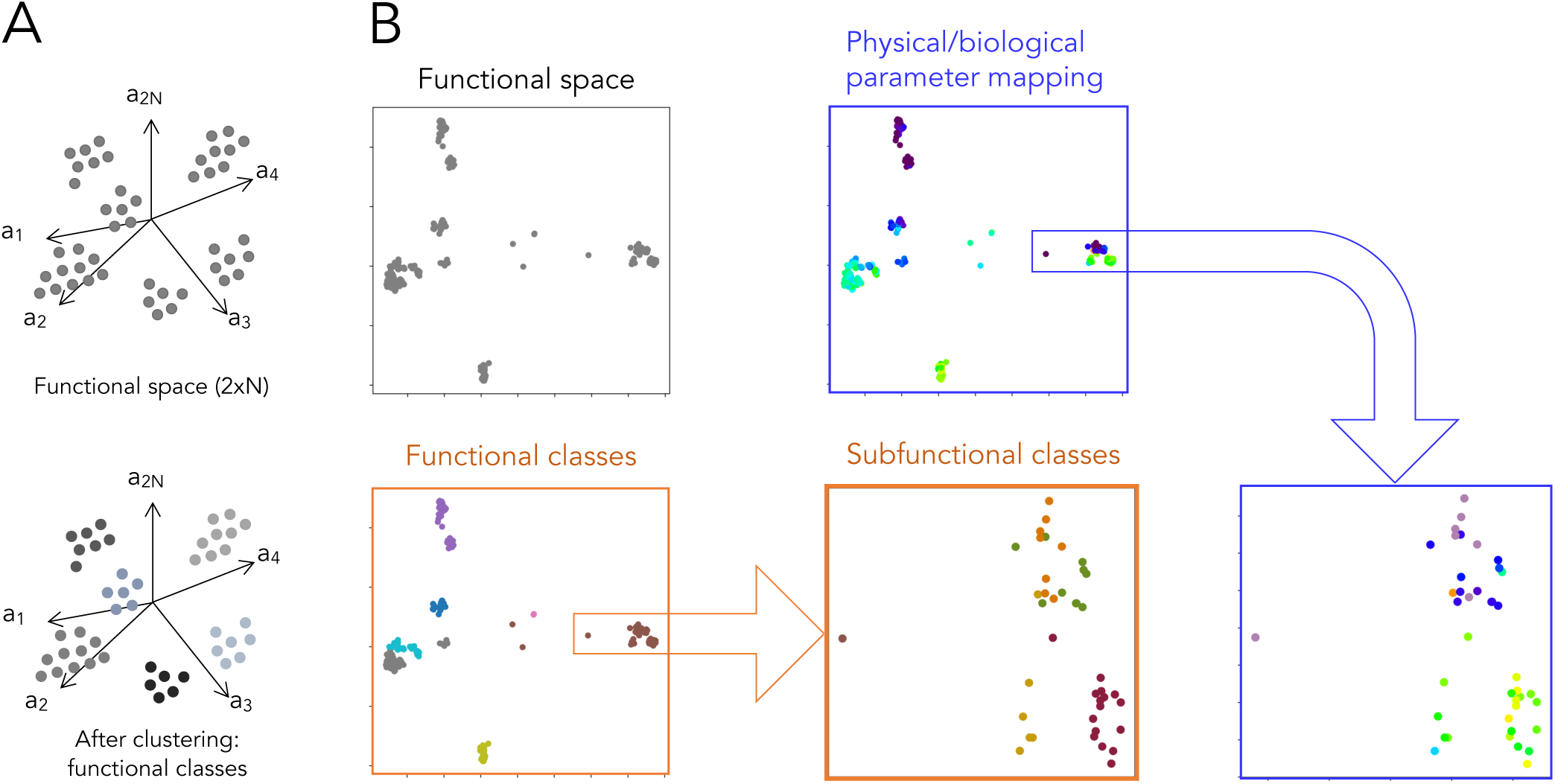
Functional embedding space, functional classes, and subfunctional classes. **A.** The schematic in Fig. 1, with (bottom) and without (top) clusters, is illustrated with an example in panel B. As explained in the text, *N* refers to the number of anchors and 2 to the number of similarity measures. **B.** Each point corresponds to a sequence represented as a vector in a 2*N* multidimensional functional space (grey panel, top). Clustering of the space (orange panels) reveals functional classes, shown as colored groups of related sequences. Zooming into selected regions, PLMView resolves subfunctional classes and supports finer-grained characterization of sequence diversity within broader functional groups. See zoom of the brown cluster (left) in the right orange panel. Biological or physical parameters (blue panels), if available, mapped onto the functional space can confirm within-class structure. Compare PLMView subfunctional classes (orange panel) with color distributions of physical parameter values (blue panel).

The complete construction is unsupervised and requires neither functional labels, structural information, handcrafted features, family-specific training, supervised learning, nor PLM fine-tuning. PLMView can therefore be applied directly to protein families for which experimental annotations are sparse or unavailable.

### Hierarchical contrasts identify residue-level determinants of specialization

The anchor-based coordinates used to construct the PLMView functional space retain a residue-level decomposition. Each anchor–query score is obtained by aggregating similarities between aligned residue embeddings, weighted by the MuLAN profile of the anchor. The position of a sequence in functional space is therefore the cumulative result of contributions from individual residues rather than an indivisible comparison between complete protein representations.

This principle is illustrated by the mass–spring analogy in Fig. 2D. Each aligned residue pair connects the query to an anchor-defined dimension, with a strength determined by the similarity of their embeddings and by the importance assigned to the anchor position. Strongly similar and highly weighted pairs exert a greater influence on the corresponding coordinate, whereas weakly related positions contribute less. The ensemble of these residue-level interactions determines the relational placement of the complete sequence.

PLMView exploits this decomposition to interpret the hierarchical partitions of the functional space. At each internal node of the functional tree, the sequence cluster represented by that node is divided into two daughter subclusters, corresponding to the two sister subtrees (Fig. 1B). Because the separation is defined from anchor-based similarity coordinates, it can be decomposed into position-specific contributions for every sequence. The magnitude of a residue contribution reflects its influence on the split, while its direction indicates which of the two daughter subclusters it supports.

To characterize the molecular signal associated with a given partition, representative sequences from the two sister subtrees are placed in a common multiple-sequence alignment. For each alignment position, PLMView compares the average residue contribution in one subcluster with that in the other. The normalized difference defines a split-specific functional score, which quantifies how strongly and in which direction that position contributes to the separation. Positions with the largest absolute differences are defined as functional determinants because their contextual PLM signals most clearly distinguish the two functional subclusters.

This definition is explicitly contrastive. A position is not identified as a functional determinant solely because it is conserved, variable, or assigned a high prior importance during the anchor comparison. The residue-level importance profile focuses the initial comparisons on potentially informative positions, whereas the functional score identifies those positions whose contributions differ systematically between the two groups produced by a specific split. Functional determinants can therefore include distributed residues outside conserved motifs and positions whose relevance depends on their sequence context.

The hierarchical organization of the tree makes it possible to examine determinants at different levels of functional resolution. Splits near the root identify positions associated with broad family-level divergence, whereas deeper splits reveal progressively more specific determinants of subfamily or sequence specialization. Following the path from the root to a selected sequence reconstructs how its functional signature is refined as it separates from increasingly similar sister subclusters. Alternatively, individual nodes can be selected to investigate a particular functional contrast.

When several nodes are considered jointly, PLMView assigns each alignment position a maximum functional score corresponding to its strongest contribution across the selected splits. This summary identifies residues that support functional discrimination at one or more hierarchical levels, while the individual split-specific scores retain information about the functional scale at which each determinant emerges. The resulting profiles can be visualized along the sequence or projected onto predicted or experimentally determined structures to generate mechanistic and experimentally testable hypotheses.

This residue-resolved framework supports the complementary applications summarized in Fig. 1C. The PLMView functional tree provides an unsupervised organization of homologous proteins into candidate functional classes. The resulting class assignments can then be projected onto phylogenetic trees to compare functional and evolutionary organization, examined among paralogous proteins to identify functional divergence in genomes, or mapped across sampling locations to generate geographical functional maps of metagenomic and metatranscriptomic data. Because each functional partition retains a residue-level decomposition, the determinants supporting individual class separations can also be mapped onto experimental or predicted protein structures to generate mechanistic and experimentally testable hypotheses.

The following case studies provide concrete implementations of these outputs across increasing levels of biological and data complexity. Thioredoxins provide an experimentally grounded system in which PLMView resolves functional subfamilies, relates their organization to phylogenetic structure, and maps distributed determinants of substrate specificity onto a protein structure; several highlighted positions overlap residues whose mutation can redirect TRX-h2 toward F-type substrate recognition. Visual opsins provide a complementary phenotype-linked benchmark, connecting inferred functional partitions and residue-level determinants to the measurable property of spectral sensitivity. Finally, cold-shock DNA-binding proteins from environmental metagenomes and metatranscriptomes illustrate how inferred functional classes can be projected across environmental samples to reveal their geographical distribution, transcriptional deployment and residue-level signatures. Together, these analyses show that PLMView not only resolves functional partitions, but also discovers functionally coherent sequence variants and their residue-level signatures in both curated multispecies protein-family datasets and large environmental sequence collections.

### Thioredoxins: resolving distributed determinants of redox specialization

Having established that PLMView links hierarchical functional partitions to residue-level contributions, we first asked whether this interpretability could recover experimentally meaningful determinants in a highly divergent protein family. Thioredoxins provide a stringent test case because they are broadly distributed oxidoreductases whose subfamilies differ in substrate repertoires, cellular localization, and regulatory roles despite sharing a conserved thioredoxin fold and catalytic motif. They therefore allow us to assess two complementary strengths of PLMView: its ability to resolve fine-grained functional subtypes within homologous proteins, and its ability to trace subtype discrimination back to specific sequence regions and residues that mechanistically support specialization. Rather than treating functional classification as a terminal label assignment, PLMView links the organization of thioredoxin subfamilies to distributed residue-level determinants, including subtle substitutions outside the conserved active site that may underlie specialized redox functions.

### From global thioredoxin diversity to algal substrate specificity

We applied PLMView to a curated dataset of thioredoxin sequences encompassing seven major types—F-, H-, M-, O-, X-, Y-, and Z—which represent distinct functional subfamilies in green plants. The resulting PLMView functional tree (Fig. 4A; see Fig. S2 for a larger view) resolves these classes into distinct subtrees despite substantial sequence divergence.

**Figure 4:**
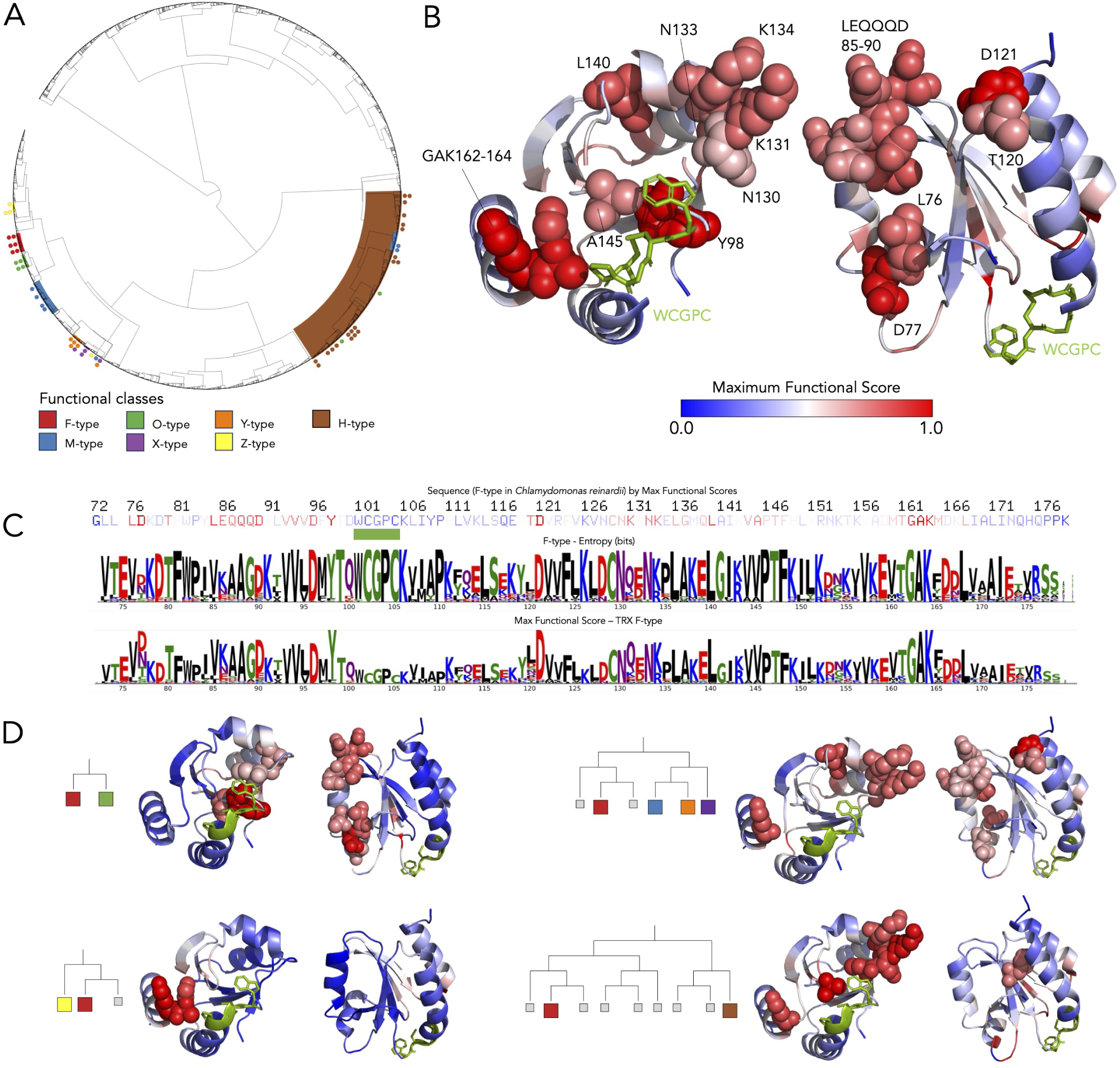
Functional analysis of thioredoxins. **A.** PLMView tree built from sequences of the thioredoxin protein family. Major subtrees are labelled according to the seven known functional classes following known annotations (outer circle): F- (red), H- (brown), M- (blue), O- (green), X- (purple), Y- (orange), and Z-types (yellow). Each color dot corresponds to an annotated sequence a functional class (Fig. S2 for a large view). **B.** Structure of the F-type thioredoxin from *C. reinhardtii* (PDB ID: 6I1C) shown in two orientations, rotated by 180*^◦^*. Residues with the highest PLMView functional scores are represented as spheres and localize predominantly around the conserved catalytic motif WCGPC (green), highlighting distributed determinants potentially involved in substrate recognition and interaction specificity (face, left). On the back of the protein (right), another group of residues with high maximum functional scores is identified. It indicates a potential interaction of TRX-f2 with some other partner. **C.** Residue-level analysis of the *C. reinhardtii* F-type thioredoxin sequence. Top: sequence representation colored according to Max Functional Scores (as in panel B). The WCGPC motif is underlined in green. Bottom: comparison between positional entropy profiles computed across F-, M-, O-, X-, Y-, Z-type thioredoxins and the PLMView Maximum Functional Score profile for the F-type. **D.** Residues identified at intermediary splits of the PLMView hierarchy. Residues and short sequence regions contributing to functional separation of F-type sequences from other TRX family types are shown for successive bifurcations from distant thioredoxin classes to sequence-specific specialization. The discriminating residues refine the functional signature associated with the target F-type thioredoxin sequence shown in panel B. Color scale as in panel B.

A TRX sequence may or may not contain an N-terminal transit peptide responsible for directing the protein to specific subcellular compartments. In plants, X-, Y-, F-, and M-type TRXs are generally targeted to the chloroplast, whereas O-type TRXs are addressed to the mitochondria. By contrast, H-type TRXs are predominantly localized in the cytosol. Because PLMView analyzes complete TRX protein sequences, including their targeting regions when present, the resulting functional tree reflects both sequence similarity and subcellular targeting information. Notably, the subtree containing the H-type TRXs also includes a limited number of O- and M-type TRX sequences that are known to lack transit peptides and therefore cluster closely with cytosolic isoforms. In contrast, TRX sequences carrying organelle-targeting peptides segregate into distinct subtrees, consistent with their specialized subcellular localization and evolutionary divergence (Fig. 4A).

A second analysis, performed after preprocessing the dataset to retain only the thioredoxin domain (Fig. S3), allowed us to test whether the PLMView topology on H- and O-types was driven primarily by targeting peptides or whether functional organization was still detectable within the conserved TRX domain itself. On the trimmed dataset, PLMView still separates chloroplastic thioredoxins (F-, M-, X-, Y-, and Z-types) from mitochondrial O-type and cytosolic H-type sequences, indicating that the classification was not induced solely by the presence or absence of N-terminal targeting regions.

The number of ambiguous placements decreases and remains very low after trimming (compare Fig. 4A and Fig. S3). The remaining cases comprise one O-type sequence, Q84XS0, placed within the H-type cluster, and one M-type sequence from the cyanobacterium *Synechocystis* placed within the Y-type cluster. Q84XS0 appears to have a shortened transit peptide relative to other plant mitochondrial O-type thioredoxins. By contrast, the cyanobacterial M-type sequence lacks a transit peptide, as expected because cyanobacterial proteins do not require organelle-targeting signals, and therefore differs in its N-terminal sequence architecture from plant chloroplast-targeted M-type thioredoxins. These placements may thus reflect genuine sequence and evolutionary heterogeneity rather than an artifact specific to PLMView.

### Phylogenetic context of thioredoxin functional specialization

Fig. S4 illustrates the phylogenetic tree of TRX sequences constructed on a sample of the sequences in Fig. 4A. The phylogeny reveals that predicted TRX functional classes are largely phylogenetically coherent, with most classes forming distinct clades consistent with shared evolutionary ancestry. Experimentally characterised proteins generally colocalise with PLMView predicted class assignments, supporting the biological relevance of the PLM-derived classification. The few misplaced sequences suggest possible functional divergence, convergent evolution, or secondary neofunctionalisation within specific TRX lineages. Although several PLM-derived functional classes display substantial phylogenetic coherence, the overall topology resembles a branching cascade rather than a set of sharply separated monophyletic groups. H- and O-types show diffuse or interleaved distributions across neighbouring clades, suggesting gradual functional diversification and partially continuous evolutionary transitions rather than strictly discrete functional partitions.

### Residue-level determinants explain substrate specificities

To investigate residue-level determinants underlying TRX functional organization, we focused on the F-type thioredoxin from *Chlamydomonas reinhardtii*. F-type thioredoxins are known to display specific substrate preferences associated with photosynthetic regulation and chloroplast redox control, distinguishing them from other thioredoxin classes such as the H-type Importantly, these functional differences are not primarily encoded by the catalytic redox-active motif, which generally follows a conserved CXXC architecture and frequently adopts the canonical WCGPC sequence in thioredoxins. Despite sequence variations across prokaryotic and eukaryotic thioredoxin-like proteins, this active-site region remains highly conserved between functional classes and therefore provides limited discriminatory power for functional specialization.

Instead, PLMView identifies distributed sequence determinants outside the catalytic core. As shown in Fig. 4D, high specificity scores localize to residues and short regions positioned around the thioredoxin scaffold rather than within the catalytic motif itself. Several of these residues correspond to positions implicated in substrate recognition and protein–protein interaction specificity in plant thioredoxins, including regions surrounding positions Y98, T99, L107, K131, N133, K134, the VAPT144–147 segment, and the GAK162–164 region^31,32^. These determinants likely contribute to shaping the interaction surface responsible for selective target recognition.

This distributed organization is further illustrated in the top of Fig. 4C, where the *C. reinhardtii* F-type thioredoxin sequence is represented together with residue-level functional scores. In this linear sequence representation, PLMView highlights multiple non-contiguous residues and short regions associated with functional specialization, emphasizing that the determinants of substrate specificity are distributed across the sequence rather than concentrated in a single motif. Fig. 4C also compares the positional entropy profiles of F-, M-, O-, X-, Y-, and Z-types thioredoxins (center) together with the PLMView maximum functional score profile (bottom). While entropy captures broadly conserved versus variable positions across subfamilies, the maximum functional score selectively enriches for residues associated with functional divergence, enabling the identification of candidate determinants that are not necessarily the most variable positions in the alignment.

Importantly, the functional score of each residue is not computed from a single comparison, but from the sequence’s trajectory through the PLMView hierarchy. At each bifurcation, from the root of the tree down to the sequence of interest, PLMView compares the residue-level contributions of the two sister subclusters and identifies positions that best explain their separation (Fig. 4D). Early bifurcations capture broad functional divergence and therefore involve many discriminating residues (bottom and top right), whereas lower bifurcations isolate progressively finer subfamily distinctions and retain a smaller, more specific set of positions (top and bottom left). The score assigned to a residue therefore reflects the strongest split along this hierarchical path at which that residue contributes to separating the target lineage from its sister lineage. Intuitively, the maximum functional score records the highest evidence, across all relevant bifurcations, that a residue acts as a functional discriminator for the sequence of interest (see Methods).

These results highlight that PLMView can recover mechanistically meaningful determinants even in highly divergent protein families where functional specificity emerges from subtle modulation of interaction surfaces rather than from discrete active-site substitutions. This conclusion is supported by previous mutational redesign experiments in which F-type determinants of the *C. reinhardtii* sequence were introduced into the TRX-h2 scaffold. Among the lysines tested to probe F-type versus H-type specificity, PLMView highlights K131, K134, and K164, with K164 lying in the GAK162–164 region implicated in TRX specificity. Introduction of F-type determinants into TRX-h2 generated a hybrid thioredoxin capable of recognizing F-type substrates, with enhanced recognition relative to the native H-type protein^32^. Thus, the residues prioritized by PLMView are not only computational markers of cluster separation, but experimentally validated determinants capable of rewiring TRX substrate specificity.

Thus, the thioredoxin analysis establishes that PLMView can recover functional partitions together with residue-level determinants that are distributed across the protein surface, providing mechanistic hypotheses for specialization rather than only assigning sequences to subfamilies.

### Visual opsins: recovering phenotype-linked determinants of spectral specialization

Having shown in thioredoxins that PLMView can link functional partitions to experimentally supported determinants of substrate specificity, we next asked whether the method could recover functional determinants in a protein family where specialized function is associated with a measurable phenotype. Visual opsins provide such a benchmark because differences in spectral sensitivity are largely determined by sequence variation around the retinal-binding pocket and have been experimentally characterized for several opsin classes. This system therefore tests whether PLMView can connect unsupervised sequence-derived functional partitions to known biophysical determinants of visual phenotype.

Opsins are light-sensitive proteins that play central roles in circadian regulation, phototaxis, and image-forming vision. Visual opsins form a subfamily of G protein–coupled receptors (GPCRs) that mediate phototransduction by binding a retinal chromophore in the photoreceptor membrane, thereby forming visual pigments capable of photon absorption. Visual opsins include several major classes, among them RH1 and RH2 opsins and short-, medium-, and long-wavelength-sensitive opsins (SWS, MWS, and LWS). Their spectral sensitivity is defined by the wavelength of maximal absorbance (*λ*_max_), which varies among opsins and determines their sensitivity to different regions of the visible spectrum. RH1 opsins mediate dim-light, or scotopic, vision, whereas RH2, SWS, MWS, and LWS opsins generally contribute to bright-light, or photopic, and color vision. Despite this functional diversification, visual opsins retain a conserved structural framework, making them particularly suitable for testing whether PLMView can recover functionally meaningful variation within a shared protein architecture.

We applied PLMView to sequences from the Visual Physiology Opsin Database (VPOD 1.3), comprising 364 animal opsin genotypes with corresponding *λ_max_* phenotypes. The dataset spans seven principal classes—RH1, RH2, MWS, LWS, SWS2, SWS1/UVS, invertebrate RH, and non-visual opsins—distributed across four phyla—Arthropoda, Annelida, Mollusca, and Chordata—including species such as *Homo sapiens*, *Diretmus argenteus*, and *Oryzias latipes*.

The resulting PLMView tree reveals distinct functional clustering (Fig. 5A; see Fig. S5 for an enlarged view). Arthropod sequences form a well-separated “Invertebrate” subtree. MWS and LWS opsins cluster proximally but segregate into distinct subtrees, consistent with their differing spectral properties, with a 17 nm difference in *λ*_max_ between the two groups. Notably, human MWS and LWS sequences are colocated, consistent with their recent divergence, high sequence identity (*≈*96%), and limited representation. SWS sequences cluster cohesively, with the established SWS1/SWS2 subdivision clearly resolved, as observed also for RH1. RH2 sequences also form a coherent cluster positioned near the RH1 sequences from *D. argenteus*.

**Figure 5:**
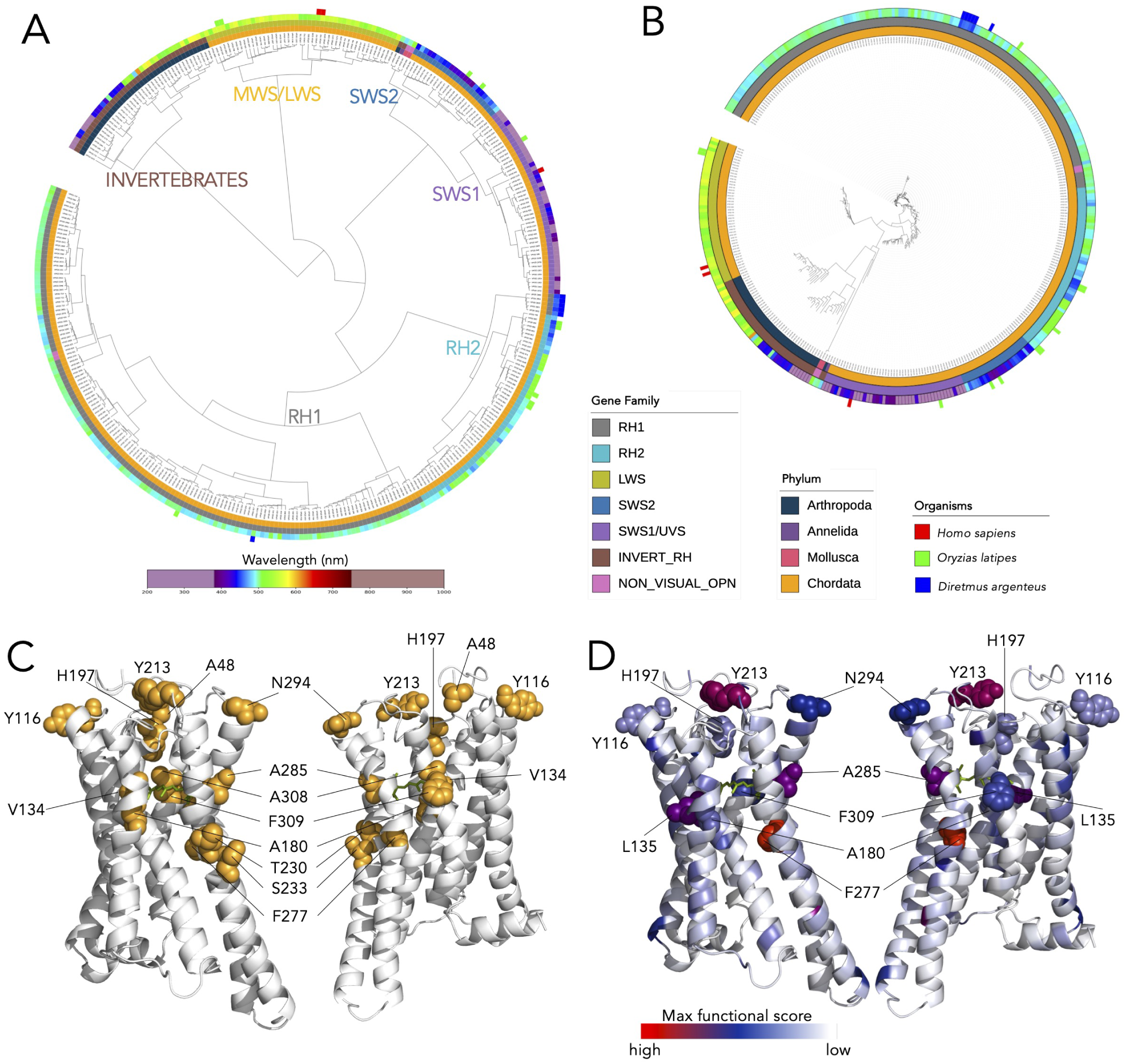
Functional analysis of visual opsins. **A.** PLMView tree built on sequences from the VPOD dataset. Subtrees are labelled by gene family names. Annotation rings from inner to outer correspond to “Phylum”, “Gene Family”, wavelength *λ_max_*, and organism species. **B.** Phylogenetic tree constructed from the same set of visual opsin sequences considered in panel A. **C.** Structure of the medium-wave-sensitive opsin from *Homo sapiens* (PDB ID: 8Y01). Orange residues correspond to experimentally validated positions associated with medium- and long-wavelength spectral tuning and red/green color discrimination. These residues include previously identified spectral tuning determinants from mutagenesis and comparative studies. The retinal chromophore (RET) is shown in green. **D.** PLMView maximal functional score, computed across 4 splits in the MWS/LWS subtree, is mapped onto the same structure. The color scale ranges from white (low specificity) to red (high specificity), with intermediate values shown in blue. Residues represented as spheres belong to the top 10% of positions with the highest PLMView maximal functional score (*≥* 0.23). Several of these positions overlap the experimentally validated spectral tuning determinants highlighted in panel C. Residue L135 is additionally highlighted because it exhibits one of the strongest maximum functional scores despite not having been previously reported in the literature. Interestingly, it is located adjacent to the experimentally characterized tuning residue V134 (see panel C). The RET is shown in green, as in panel C.

The deep-sea fish *D. argenteus* exhibits lineage-specific diversification of RH1 opsins associated with adaptation to low-light environments, characterized by loss of color-discriminating opsins and enhanced sensitivity to bioluminescent wavelengths. In the PLMView tree, the *D. argenteus* RH1 sequences form a distinct subtree positioned between the main RH1 and RH2 subtrees. They cluster near the RH2 sequences of *O. latipes*, reflecting their similar spectral properties, while remaining separated into two independent subtrees that preserve their respective RH1 and RH2 identities.

In contrast, the corresponding phylogenetic tree (Fig. 5B; see Fig. S6 for an enlarged view) does not recover these functional subdivisions; all *D. argenteus* sequences cluster within a single phylogenetic subtree, underscoring the limitations of purely evolutionary reconstructions for resolving functional differentiation.

### Functional determinants explain visual opsins subfamilies

To test the ability of PLMView to identify residue-level determinants that mechanistically explain subfamily differences, we evaluated whether it could distinguish the medium- and long-wavelength-sensitive opsin subfamilies, MWS and LWS.

The analysis is associated with the node in Fig. 5A labelled MWS/LWS. Fig. 5C highlights residues previously demonstrated experimentally to contribute to spectral tuning and red/green color discrimination in vertebrate visual opsins. These include the classical “five-sites” residues A180, H197, F277, A285, and A308^33^, together with additional experimentally validated tuning positions including residues Y213 and N294^34^, A48 and V134^35^, and Y116, T230, S233, and F309^36^. Collectively, these residues are distributed across multiple transmembrane helices surrounding the retinal-binding pocket and are known to modulate wavelength sensitivity through local electrostatic and steric effects on the chromophore environment.

We therefore computed residue-level functional specificity scores across selected splits within the MWS/LWS subtree, chosen based on differences in opsin *λ*_max_. As shown in Fig. 5D, PLMView identifies a concentrated set of high-scoring residues that spatially colocalize with known spectral tuning positions. Remarkably, eight experimentally validated residues shown to explain major shifts in *λ*_max_ between MWS and LWS pigments are recovered among the strongest specificity signals: Y116, A180, H197, Y213, F277, A285, N294, and F309.

PLMView also identifies residue L135 among the positions with the highest functional scores. Although this residue has not previously been reported in the literature, it localizes adjacent to the experimentally characterized tuning residue V134, suggesting that PLMView may capture previously unrecognized local determinants contributing to spectral specialization.

Importantly, the functional score does not simply highlight the most conserved positions in the family; rather, it identifies residues whose contributions to embedding alignment differ most strongly between sister subclusters along the hierarchical path from the root to individual sequences, thereby capturing positions specifically associated with functional divergence. The enrichment of experimentally validated tuning residues among the highest-scoring positions, together with the identification of neighboring residues and structural regions surrounding the retinal-binding cavity, indicates that PLMView recovers both established determinants and distributed contextual effects underlying spectral specialization. These results suggest that the geometric organization of sequences in PLMView functional space reflects the biophysical mechanisms governing opsin wavelength sensitivity.

Together, the thioredoxin and opsin analyses show that PLMView captures complementary modes of functional specialization: distributed determinants of substrate specificity and localized, context-dependent determinants of a measurable phenotype. These benchmarks support its application to environmental protein families, where annotations are sparse and the aim is to discover sequence-defined variants and their candidate molecular determinants.

### Cold-shock proteins: from protein-family specialization to ocean-scale functional discovery

We applied PLMView to the *Tara* Oceans metagenomic (MetaG) and metatranscriptomic (MetaT) datasets. These resources provide a global view of marine protein diversity and are commonly used to characterize the distribution of functional potential and activity across the ocean through protein-domain abundance^29^. Although effective for identifying broad ecological trends, this approach treats each domain as a homogeneous functional unit, implicitly assuming that all domain instances contribute equivalently to biological function.

Here, we move from domain counting to functional classification of domain instances. Rather than asking only how abundant a domain is in MetaG or MetaT data, PLMView asks which sequence-defined functional variants of that domain are present, where they occur, and which of them are transcriptionally deployed. As a proof of concept, we consider cold-shock DNA-binding domain proteins in diatoms. PLMView partitions PF00313-containing sequences into fine-grained potential functional subtypes that can be quantified across *Tara* Oceans stations, organismal size fractions, and MetaG versus MetaT datasets. Even when the precise biochemical roles of individual variants remain unknown, their coherent sequence organization, environmental distribution, transcriptional activity, and residue-level signatures can provide concrete hypotheses about functional diversification and ecological specialization.

The underlying datasets encompass more than 200 sampling stations across major oceanic basins and latitudinal gradients, enabling the global distribution of sequence-defined cold-shock domain variants to be examined across stations, size fractions, and MetaG and MetaT datasets. An example of these distributions is shown for the MetaT mesoplankton-size fraction in Fig. 6A; the full planetary-scale analysis is presented below.

**Figure 6:**
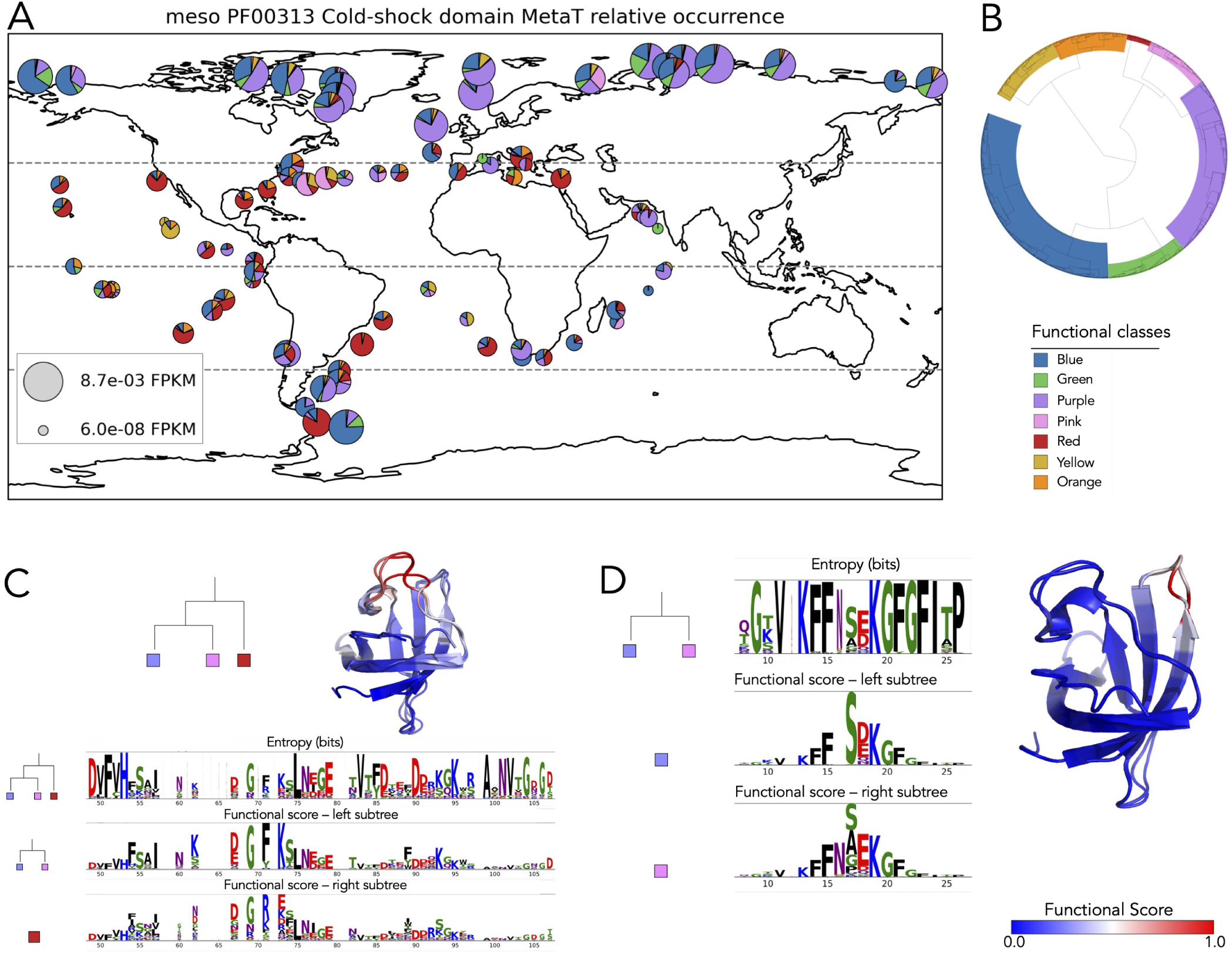
Mapping of a cold-shock DNA binding domain in diatom sequences from the *Tara* Oceans expedition. **A.** Map of functional metatranscriptomic abundance of cold-shock DNA binding domain-containing sequences derived from the *Tara* Oceans data for meso-scale size diatoms. Each sampling station is described by a pie chart reporting sequence abundance for seven functional classes. Colors are as in B. **B.** PLMView of cold-shock sequences as from A. Seven major functional subtrees are identified with PLMView using a tree depth of 3 and colored accordingly. **C.** Residue-level functional determinant analysis for the bifurcation separating the combined Purple+Pink partition from the Red partition. Functional scores (see bottom right for color scale) are mapped onto representative AlphaFold 2^37^ model structures predicted from the Purple, Pink, and Red functional classes. Below, positional entropy profiles and subtree-specific functional score profiles compare the Purple+Pink and Red partitions, highlighting residues that contribute specifically to their functional separation. **D.** Residue-level functional determinant analysis for the bifurcation separating the Purple and Pink functional partitions. Functional scores (see bottom right for color scale) are mapped onto representative AlphaFold 2^37^ model structures predicted from the Purple and Pink functional classes. On the left, positional entropy is compared to subtree-specific functional score profiles. This analysis identifies a refined set of residues discriminating closely related Purple and Pink variants, consistent with functional specialization within the Purple/Pink branch.

### A planetary-scale map of cold-shock DNA-binding domain variants in diatoms

To reconstruct the global functional landscape of PF00313 cold-shock DNA-binding domain-containing proteins in diatoms, PLMView partitioned the cold-shock domain sequences into seven distinct sequence-defined functional subfamilies (Fig. 6B; see Fig. S7 for an enlarged view). Their phylogenetic organization (Fig. S8) does not fully mirror their proximity in the PLMView tree, most notably because sequences assigned to the Yellow/Orange partition were distributed across separate phylogenetic subtrees.

The PLMView subfamilies were then used as functional classes to assign diatom sequences and quantify their relative abundance at each *Tara* Oceans station. We mapped the global distribution of the seven classes in both MetaG (Fig. S9) and MetaT (Fig. S10) datasets to compare genomic potential with transcriptional activity. To examine the ecological organization of these variants across organismal size fractions, we repeated the analysis separately for diatom sequences in pico-, nano-, micro-, and meso-plankton-size fractions. The corresponding MetaG maps are shown in Fig. S11, Fig. S12, Fig. S13, and Fig. S14, respectively, whereas transcriptional activity maps are shown in Fig. S15, Fig. S16, Fig. S17, and Fig. 6A (Fig. S18).

Across metagenomes, all partitions are broadly distributed and frequently co-occur within the same sampling stations, indicating that the cold-shock repertoire of diatoms is globally conserved at the DNA level. However, metatranscriptomic profiles display sharper spatial structuring (compare Fig. S9 and Fig. S10). The Yellow/Orange partition is abundant and geographically widespread in metagenomes: Yellow variants are preferentially enriched in temperate and subtropical regions, whereas Orange variants extend toward temperate and polar waters. Despite their broad genomic occurrence, both partitions remain only weakly represented in metatranscriptomic datasets across most ocean basins and size fractions. This suggests that a substantial fraction of PF00313 sequence diversity corresponds to latent or conditionally deployed functional potential rather than constitutively active transcriptional programs.

In contrast, the Blue and Purple partitions dominate transcriptional activity, especially at high latitudes. Arctic and Antarctic stations are consistently enriched in Blue- and Purple-associated transcripts, both in total-community maps and within individual size fractions, suggesting that these variants likely encode the primary cold-responsive functional states actively deployed under chronically low-temperature conditions. This polar enrichment is particularly pronounced in picoplanktonic and nanoplanktonic size fractions, where Blue and Purple variants frequently account for the majority of transcriptional signal (Fig. S15, Fig. S16). More generally, meta-transcriptomes exhibit substantially reduced functional diversity relative to metagenomes, with transcription converging toward a limited subset of highly active partitions despite broader underlying genomic repertoires.

At equatorial and subtropical stations, the relative contributions of Green-, Red-, and Yellow-associated transcripts increase, although the pattern varies among size fractions and Blue remains prominent at many stations. Green and Yellow variants make substantial contributions to several picoplankton MetaT profiles (Fig. S15), whereas Red variants are more prominent in larger size fractions (Fig. S17 and Fig. 6A, respectively). Nano- and microplankton size fractions exhibit mixed functional compositions, with variable contributions from Green, Pink, and Red variants (Fig. S16 and Fig. S17). Mesoplankton size fractions show particularly pronounced geographic turnover, shifting from predominantly Blue- and Purple-associated transcription at high latitudes toward greater representation of Red, Yellow, Orange, and Pink variants at temperate and subtropical stations (Fig. 6A). These patterns are consistent with larger-size diatom communities deploying a broader repertoire of cold-shock-domain variants under warmer or more variable environmental conditions.

Together, these patterns extend the PLMView hierarchy beyond sequence-level organization and reveal a nested functional architecture that links molecular diversification to plankton size structure and global environmental gradients. Cold-shock domain diversification in diatoms therefore appears to reflect not only sequence-level variation but also ecological specialization across thermal regimes and organismal scales. Integrating genomic occurrence with transcriptional deployment reveals how distinct functional states are selectively activated across the global ocean, providing a systems-level view of how diatoms partition and regulate cold-associated molecular functions. Having established this ecological stratification, we next examined the residue-level determinants that distinguish the major functional partitions.

### Extraction of functional determinants for cold-shock proteins in the ocean

Cold-shock domains are conserved nucleic-acid-binding folds that recognize single-stranded RNA and DNA through a shared binding surface, generally with limited sequence or substrate-type selectivity^38,39^. In prokaryotic and eukaryotic proteins, this surface includes exposed residues and loop regions that contact the nucleic-acid substrate. Several of the strongest determinants identified by PLMView map to these recognition regions, suggesting that the inferred functional partitions reflect differences in nucleic-acid binding rather than changes in the overall CSD fold.

Functional determinants within the PF00313 family were extracted by analyzing successive bifurcations in the PLMView hierarchy and computing residue-level functional scores between sister subtrees. For the separation of the combined Purple & Pink partition from the Red partition, Fig. 6C highlights multiple non-contiguous residues and short sequence regions—particularly within the *β*_3_–*β*_4_ loop (red residues in the structure), which has been implicated in substrate discrimination between ssDNA and dsDNA^40^. Whereas positional entropy captures globally variable alignment positions, PLMView selectively identifies residues associated with the functional separation of these partitions. At a finer hierarchical level, Fig. 6D highlights residues separating the Purple and Pink classes, located within the *β*_1_ *− β*_2_ loop (red residues in the structure). Although a direct role for this loop in substrate discrimination has not been experimentally established, structural studies have identified conformational changes in the *β*_1_–*β*_2_ loop upon ssDNA binding, alongside changes in the *β*_3_–*β*_4_ loop^39^.

Together, these analyses reveal distinct molecular signatures associated with broad functional divergence and closely related specializations. PLMView thus progressively resolves functional variation across hierarchical scales, from major family-level partitions to fine-grained sub-family distinctions. Importantly, these determinants emerge from an unsupervised analysis of protein language-model representations, without reliance on prior structural or biochemical annotations, enabling the exploration of molecular specialization in environmental protein families that remain largely uncharacterized.

### A sequence-to-ecology framework for functional discovery

Across these three applications, PLMView supports a unified sequence-to-ecology workflow. In thioredoxins, it identifies distributed determinants of redox-related specialization. In opsins, it recovers residues associated with experimentally characterized spectral tuning. In *Tara* Oceans cold-shock proteins, it extends the same residue-resolved analysis to environmental metagenomic and metatranscriptomic sequences, revealing functionally distinct variants with different geographic and transcriptional profiles. These results show that PLMView can be used not only to classify homologous proteins, but also to discover functional structure within protein families and project that structure onto ecological data.

### Comparison with Sequence Similarity Networks

Sequence Similarity Networks (SSNs) are commonly used to infer functional relationships among homologous protein sequences. Details on the construction of SSNs are given in Methods.

Despite their widespread use, SSNs present important limitations when applied to large-scale protein family datasets. SSNs rely directly on sequence alignment scores and therefore primarily capture evolutionary proximity, which may limit their ability to resolve subtle functional specialization among closely related homologs. This limitation may become particularly pronounced in highly diverse protein families, where sequence similarity alone may fail to distinguish proteins with distinct biochemical or ecological functions. To examine this limitation, we analysed three protein families using SSNs and compared the resulting reconstructions with PLMView trees.

On the TRX dataset, SSNs and PLMView showed broadly comparable local class coherence, with sequences of the same functional type generally grouping together (Fig. S19 and Fig. S20). However, the SSN reconstruction placed the H-type cluster within the O-type cluster and less clearly separated cytosolic, mitochondrial, nuclear, and chloroplastic thioredoxins. PLMView instead placed cytosolic H-type thioredoxins on a distinct subtree and chloroplastic thioredoxins in a separate functional region. After sequences were trimmed to the thioredoxin domain (Fig. S3), PLMView retained this overall separation, while the few ambiguous placements were also observed in the SSN dendrograms, suggesting that they might reflect intrinsic sequence similarity rather than a PLMView-specific artifact.

On the VPOD dataset, SSNs perform comparably to PLMView, with the main difference concerning the placement of the *D. argenteus* RH1 sequences. In the SSN dendrograms (Fig. S21 and Fig. S22), canonical RH1 sequences cluster closer to RH2 than to *D. argenteus* RH1. By contrast, PLMView places the latter between RH1 and RH2 (Fig. 5A), more consistently reflecting their distinct wavelength sensitivities.

To examine a case in which sequence similarity is particularly limiting, we analysed 307 Cryptochrome/Photolyase Family (CPF) sequences previously classified in^28^. CPF proteins retain a conserved FAD-binding core despite performing diverse functions, including DNA repair, photoreception, circadian regulation, signalling, and transcriptional regulation. Together with the presence of fragmented sequences, this conservation can produce two opposing problems in SSNs: functionally distinct lineages may remain collapsed, whereas coherent classes may become fragmented.

We therefore applied PLMView using anchors trimmed to the FAD-binding domain (PF03441), while analysing query sequences at full length. The resulting tree resolves experimentally characterized CPF proteins into eleven sharply delimited functional subtrees (Fig. S23), more clearly than the corresponding SSN reconstructions (Fig. S24 and Fig. S25). The contrast is most pronounced around the (6-4)-photolyase-related subclass: PLMView separates transcriptional regulators, animal photoreceptor CRYs, and (6-4)-photolyases, whereas the SSN partially merges these groups. Conversely, Class I CPD photolyases are fragmented across several SSN sub-trees but form a coherent PLMView group. Thus, CPF provides a stringent example in which PLMView resolves specialized lineages that sequence similarity either collapses or fragments.

Beyond differences in functional resolution, the two approaches also differed substantially in computational scalability. Because pairwise similarities must be computed for all sequence pairs, SSN construction scales quadratically with the number of sequences (*O*(*n*^2^)), resulting in substantial computational and memory costs for large protein families or metagenomic datasets. In practice, this often requires aggressive filtering, thresholding, or graph-sparsification strategies that can obscure weak but biologically meaningful relationships. The large TRX dataset provides a concrete illustration of this computational difference. For more than 30,000 TRX sequences, data preparation—including domain annotation and trimming with MetaClade2^41^—required 1 h 35 min. PLMView then classified the trimmed sequences in 50 min, whereas SSN reconstruction on the same dataset required approximately 14 h, followed by an additional 1 h to construct the linkage dendrogram. This corresponds to an approximately 18-fold speed-up in classification and tree construction. The difference is largely explained by the substantially smaller number of pairwise comparisons performed by PLMView: approximately 40 million embedding comparisons, compared with 965 million sequence comparisons for the SSN. Thus, while PLMView recovered a biologically coherent classification comparable to, and in some aspects topologically clearer than, SSN-based reconstruction, it remained substantially faster and more scalable for large protein-family analyses.

### Comparison with pooled PLM embedding representations

Representing protein sequences by averaging residue-level PLM embeddings across sequence length has become a widely used strategy for protein classification^13–15^. We therefore compared the functional organization obtained from pooled ESM2-3B and Ankh-Large embeddings with that recovered by PLMView for two protein families analysed in this study, thioredoxins and visual opsins, and extended the comparison to the CPF family.

The thioredoxin dataset provides a stringent test case because TRX functional subclasses are closely related and can differ by both thioredoxin-domain determinants and subcellular targeting signatures. Fig. S26 and Fig. S27 show classifications obtained directly from pooled ESMB2-3B and Ankh-Large embeddings alone, respectively, without the collaborative multi-view architecture of PLMView. Because these analyses were performed on trimmed TRX sequences, they test whether pooled embeddings can recover functional organization from the thioredoxin domain after removal of transit peptide regions.

The dendrogram built from averaged ESM2-3B representations (Fig. S26) isolates most H-type thioredoxins, corresponding to the cytosolic class, from the other sequences, with the exception of one O-type sequence. However, the remaining classes are less clearly resolved, with several misplacements involving M-, F-, O-, X-, Y-, and Z-type thioredoxins. Thus, pooled ESM2-3B embeddings capture a coarse H-type versus non-H-type distinction but provide limited resolution among chloroplastic and mitochondrial TRX subclasses.

The Ankh-Large dendrogram in Fig. S27 makes the same broad separation between H-type thioredoxins and the rest of the family, but displays greater difficulty in separating the remaining TRX types. Although some local groupings are recovered, F-, M-, O-, X-, Y-, and Z-type sequences remain partially intermixed. This suggests that, for TRXs, part of the functional signal encoded in raw PLM embeddings is lost when residue-level representations are averaged across the domain. In contrast, the PLMView reconstruction (Fig. 4A) retains higher functional resolution by comparing sequences through anchor-based residue-level similarities, thereby better preserving the distributed determinants that distinguish closely related TRX subclasses.

We next analysed visual opsins, where functional specialization is associated with spectral sensitivity and therefore provides a phenotype-linked benchmark. Fig. S28 and Fig. S29 show classifications obtained directly from pooled ESM2-3B and Ankh-Large embeddings, respectively. These reconstructions reflect the intrinsic geometry of the pooled embedding spaces rather than the relational, anchor-based functional organization recovered in Fig. 5A.

Consistent with the embedding analyses presented in the Supplementary Methods, the ESM2-3B classification appears comparatively compressed and less discriminative, likely due to the strong global similarity and anisotropy of ESM embeddings, which reduce contrast between residue and sequence representations. This is reflected in the topology of Fig. S28, where the dendrogram separates the dataset into four main regions but only partially recovers the expected functional organization. One subtree groups invertebrate sequences split into two separated groups and interleaved with non-visual Chordata opsins and RH1 Mollusca opsins. A second subtree is enriched in RH2 sequences but also contains RH1 and SWS1 sequences. A third correctly groups the MWS/LWS and SWS1/SWS2 subtrees, whereas a fourth collects most of the remaining RH1 sequences. Thus, pooled ESM2-3B embeddings capture broad opsin-family structure, but they do not fully resolve the functional subdivisions recovered by PLMView.

By contrast, the Ankh-Large-based reconstruction (Fig. S29) preserves sharper local organization and clearer functional separation, consistent with the broader variance distribution and stronger residue-level localization observed for Ankh embeddings (Supplementary Methods). One subtree correctly groups invertebrate sequences, whereas a neighboring subtree collects non-visual Chordata opsins together with Rh1 Mollusca opsins. A second region groups the MWS/LWS and SWS1/SWS2 subtrees, while a third groups RH1 and RH2 sequences, although RH1 sequences remain split into two independent subtrees. Thus, Ankh-Large improves local coherence relative to ESM2-3B, but the pooled representation still fails to reproduce the full functional hierarchy obtained with PLMView, particularly for closely related opsin classes associated with subtle spectral-tuning residues.

Finally, we considered the CPF protein family, which extends the comparison to a functionally diverse family sharing a common FAD-binding cryptochrome/photolyase core. The ESM2-3B dendrogram (Fig. S30) shows stronger fragmentation of functionally characterized sequences sharing the same annotation than the Ankh-Large dendrogram (Fig. S31). The ESM2-3B tree recovers some broad CPF organization, but several functional groups are split across neighboring branches or positioned at interfaces between subtrees. Ankh-Large displays more coherent sectors and reduced fragmentation, indicating a stronger functional signal for CPF classification. Nevertheless, both pooled-embedding reconstructions remain less sharply organized than the PLMView CPF tree, where functionally characterized CPF lineages form more clearly delimited subtrees.

Overall, these comparisons indicate that raw PLM embedding geometry already captures part of the functional organisation, especially for Ankh-Large. Across the TRX, visual opsin, and CPF datasets, however, pooled embeddings remain less functionally resolved than PLMView. The limitation is not the absence of functional information in the PLM representations, but the loss of residue-level and relational information caused by averaging embeddings into a single sequence-level vector. PLMView substantially enhances this signal by integrating residue-level weighting, anchor-based comparisons, and hierarchical multi-view clustering. This collaborative representation yields a more interpretable functional hierarchy and improves the resolution of biologically meaningful specialization across all three protein families.

### Tree reconstruction using FANTASIA embedding-similarity metrics

In the previous section, averaged embeddings were clustered using Ward’s method (Methods). Here, we instead reconstruct the trees using the embedding-similarity measures implemented in FANTASIA^14^, namely cosine similarity and RBF similarity, this latter derived from Euclidean distance. FANTASIA uses similarity in embedding space to identify reference proteins from which Gene Ontology annotations can be transferred. Because FANTASIA and PLMView were developed for different purposes, this analysis is not intended as a direct method comparison: FANTASIA targets large-scale functional annotation transfer, including for proteins that are difficult to annotate using conventional homology-based approaches, whereas PLMView aims to reconstruct fine-grained, family-specific functional organization from sequence relationships.

We applied the FANTASIA similarity metrics to the VPOD dataset. Fig. S32 and Fig. S33 show the classifications obtained using FANTASIA V1/ProtT5 and FANTASIA V2/ProstT5 representations, respectively. Both dendrograms recover some coarse structure, most notably the broad separation of a subset of invertebrate opsins. However, the remaining sequences are less consistently organised accordingly to the functional annotation rings than in the PLMView reconstruction (Fig. 5A). In particular, RH1, RH2, SWS, MWS/LWS, non-visual opsins, and phylum-specific groups remain partially intermixed rather than forming compact and interpretable functional regions.

These reconstructions confirm that the embedding similarities used in FANTASIA contain functional signal, but provide limited resolution of the fine subdivisions associated with visual-opsin specialization. This difference is consistent with the different design goals of the methods: FANTASIA similarities support broad annotation transfer, whereas PLMView reorganizes PLM information into a family-specific, anchor-based functional space and uses residue-level weighting to distinguish closely related opsin classes.

## DISCUSSION

This study presents PLMView as a flexible framework for functional discovery from protein sequences. Rather than treating function as a label transferred from characterized homologs, PLMView reconstructs the internal functional architecture of protein families by comparing sequences in a collaborative PLM-derived space. This makes the method particularly suited to environmental genomics, where sequence diversity is vast, experimental annotation is sparse, and biologically meaningful differences often occur below the level of conventional domain, family, or ontology-based annotations.

The three applications illustrate complementary aspects of this framework. In thioredoxins, PLMView resolves functional organization within a divergent redox protein family and identifies distributed residue-level determinants of substrate specificity. The overlap between PLMView determinants and residues previously used to redesign TRX-h2 toward F-type substrate recognition shows that the method can recover experimentally actionable positions, not only statistical markers of cluster separation. In visual opsins, PLMView recovers residues associated with an experimentally measurable phenotype, spectral tuning, including determinants located around the retinal-binding pocket. Finally, in *Tara* Oceans cold-shock DNA-binding proteins, the same residue-resolved framework is extended to environmental metagenomic and metatranscriptomic sequences, where PLMView identifies sequence-defined functional variants with distinct geographic distributions, transcriptional profiles, and candidate molecular determinants. Together, these examples show that PLMView can connect residue-level specialization, protein-family organization, and environmental deployment within a single analytical framework.

This perspective differs from conventional annotation pipelines, which often operate through label projection. Homology-based transfer, curated ontologies, supervised PLM predictors, and structure-based annotation methods are powerful when reference functions are available, but they necessarily organize new sequences around pre-existing functional categories. As a result, they tend to perform best in well-characterized regions of sequence space and may compress unexplored diversity into broad or generic labels. This limitation is particularly important in environmental datasets, where many sequences belong to recognizable domains but may represent uncharacterized functional variants. PLMView addresses this gap by resolving functional structure within protein families before formal labels are available, thereby shifting annotation from retrospective label assignment toward prospective hypothesis generation.

The contrast is especially clear in large-scale environmental analyses. Many global metaG and metaT studies quantify the abundance of protein domains to infer functional potential and activity across ecosystems. This strategy has revealed major ecological trends, but it treats each domain as a homogeneous functional unit. PLMView adds a complementary layer of resolution by classifying individual domain instances into functional subtypes directly from sequence. In the *Tara* Oceans cold-shock protein analysis, this shift from domain counting to functional classification makes it possible to ask not only how abundant a domain is, but which functional variants of that domain are present, where they occur, and which variants are transcriptionally deployed. This distinction is essential for separating genomic functional potential from realized activity and for detecting ecological structure hidden within broadly defined functional categories.

A central advantage of PLMView is that functional partitions are interpretable at the residue level. The method does not stop at assigning sequences to subfamilies; it traces the separation of subfamilies back to positions whose contributions differ across hierarchical bifurcations. This is important because functional specialization is often encoded by sparse, context-dependent substitutions rather than by whole-domain changes or obvious motif replacement. In thioredoxins, these determinants are distributed across interaction surfaces rather than confined to the conserved catalytic motif. In opsins, high-scoring residues include experimentally characterized tuning sites and neighboring positions around the retinal-binding cavity. In cold-shock proteins, PLMView identifies residues and short sequence regions that distinguish environmental variants. In each case, the method links geometric separation in functional space to candidate molecular mechanisms.

This residue-level interpretability also distinguishes PLMView from approaches based only on global sequence similarity, pooled PLM embeddings, or structural clustering. Sequence similarity and phylogeny provide important evolutionary context, but they may not resolve the specific substitutions that underlie functional divergence. Similarly, structural similarity can group proteins with related folds, but proteins sharing a fold may still differ in substrate specificity, interaction partners, localization, regulation, or ecological role. PLMView complements these approaches by identifying functionally coherent partitions within homologous families and by highlighting the residues that support those partitions. In this sense, the method is not intended to replace phylogenetic, structural, or ontology-based analyses, but to add a discovery-oriented layer between raw sequence diversity and formal functional annotation.

The unsupervised nature of PLMView is particularly valuable in sparse-label regimes. Supervised predictors, including PLM-based methods trained or fine-tuned on curated functional labels, inherit the biases of existing databases: well-studied proteins, canonical pathways, and experimentally characterized model systems are overrepresented, whereas lineage-specific, environmentally restricted, or weakly characterized variants remain underrepresented. PLMView avoids this dependency on predefined labels by using intrinsic structure in PLM-derived sequence relationships. This allows it to identify coherent functional subgroups even when their precise biochemical roles are not yet known. Such subgroups can then be prioritized for experimental characterization, ecological interpretation, or integration into future annotation schemes. At the same time, PLMView should be viewed as a framework for generating functional hypotheses rather than as a definitive substitute for biochemical validation. The method requires a homologous family context and a sufficiently informative set of sequences from which anchor-based relationships can be constructed. It is therefore best suited to families where multiple related sequences are available, rather than to isolated orphan proteins with no meaningful comparative context. Moreover, sequence-defined variants inferred by PLMView may reflect differences in biochemical activity, interaction specificity, localization, regulation, or ecological deployment; distinguishing among these possibilities requires additional experimental, structural, or systems-level evidence. The strength of the approach lies in narrowing this search space by identifying coherent variants and the residues most likely to underlie their specialization.

Taken together, our results support PLMView as a general framework for moving from protein sequence variation to functional discovery. In well-studied or experimentally tractable families, it recovers interpretable determinants that agree with known mechanisms or redesign experiments. In phenotype-linked systems, it identifies residues associated with measurable functional properties. In environmental datasets, it reveals hidden functional structure within protein domains and projects this structure onto geographic and transcriptional landscapes. By linking sequence-defined variants, residue-level determinants, and ecological distributions, PLMView provides a route toward discovery-oriented annotation of the expanding protein universe, with particular relevance for large-scale environmental and evolutionary genomics.

By transforming learned protein representations into interpretable, relational functional maps, PLMView bridges modern representation learning with classical comparative biology. Rather than treating PLM embeddings as isolated sequence descriptors, PLMView organizes proteins through their relationships to homologous anchors, thereby converting dense representations into functional landscapes that can be explored hierarchically and interpreted at residue-level resolution. This relational perspective enables scalable discovery of functional specialization across the protein universe while preserving a direct link between functional partitions and candidate molecular determinants. At larger scales, however, the challenge is not solely computational but also representational, as trees containing hundreds of thousands or millions of leaves become impractical to navigate and interpret. Scaling functional classification also requires preserving the distinction between sequences that can be confidently assigned to established classes and those that fall outside the functional diversity represented by the reference set. This issue is particularly important in metagenomic datasets, where divergent and poorly characterized proteins may encode previously unrecognized functional specializations. Methods such as SPIN^42^ provide a framework for extending classification to very large sequence collections while retaining the ability to identify sequences that are not adequately explained by existing classes. Such sequences should not be obscured by large-scale assignment, as they represent important targets for subsequent functional investigation.

An important next step for PLMView will be to connect residue-level determinants to more explicit biological features encoded within PLMs. Recent interpretability approaches based on sparse autoencoders (SAEs)^20,21^ have shown that dense PLM embeddings can be decomposed into sparse, more interpretable feature spaces. In such representations, each residue is encoded by a high-dimensional sparse vector whose active dimensions can often be associated with recognizable biochemical, structural, or functional properties. PLMView and SAE-based interpretability are therefore highly complementary: PLMView identifies which residues contribute to functional separation within a protein family, whereas SAEs could help explain why these residues are important by associating them with interpretable latent features.

Coupling PLMView functional scores with SAE-derived sparse features would thus add an additional layer of mechanistic interpretation, linking hierarchical functional partitions not only to specific residues, but also to the biological attributes represented at those positions. In practice, this would allow one to ask whether residues highlighted by PLMView correspond to sparse features related to catalytic motifs, binding interfaces, structural environments, localization signals, or other functional properties. At the time of writing, however, no sparse autoencoder has yet been trained on Ankh-Large, the model underlying most of our analyses. Extending SAE-based interpretability to Ankh-Large, or applying PLMView to PLMs for which SAEs are already available, therefore represents a promising direction for future work.

## METHODS

### Protein Language Model selection and embedding representation

Protein language models (PLMs) leverage large corpora of natural protein sequences to learn the underlying statistical rules that govern protein structure and function. Analogous to natural language models, PLMs use Transformer-based architectures^43^ to infer the contextual relationships among amino acids along the sequence. Through pretraining on masked residue prediction tasks, they capture both local and long-range dependencies, producing rich residue embeddings that implicitly encode biochemical, structural, and evolutionary information.

These embeddings have been successfully applied to diverse problems—from structure prediction^44^ and mutational effect estimation^45,46^ to functional site identification^47^. In this study, we primarily used Ankh-Large, a PLM shown to outperform other models in capturing alignment-like relationships^24,48^. Each sequence is represented as a matrix of high-dimensional vectors, where each vector corresponds to an amino acid residue. We refer to this representation as the sequence embedding, and to each individual vector as a residue embedding.

Detailed comparisons between Ankh-Large and ESM2-3B embeddings are provided in the Supplementary Methods (see the supplementary text). Briefly, our analysis reveals marked differences in the geometric organization of the embedding spaces. ESM2-3B embeddings exhibit strong global components and in-sequence anisotropy, resulting in elevated baseline cosine similarity across residue representations. In contrast, Ankh-Large embeddings display greater variability and stronger localization of information at the residue level, properties that are advantageous for residue-resolved functional comparisons in PLMView.

### Collaborative learning architecture

PLMView is designed to classify specialized protein functions—the distinct biochemical or regulatory roles that emerge among homologous sequences—directly from PLM embeddings. The method integrates complementary PLM representations, identifies informative residues, and clusters sequences hierarchically in a collaborative embedding space. Detailed procedures for anchor selection, PLM embedding generation, multi-view similarity computation, collaborative embedding representation, functional similarity metrics, and hierarchical clustering, are provided in the Supplementary Methods.

### Weighting of functionally informative residues

Functional specialization often arises from a small subset of critical residues. To highlight such positions in anchors, PLMView integrates the MuLAN attention mechanism^30^, which provides a rapid estimate of residue-level functional importance. For each anchor sequence, MuLAN scores are computed directly from the PLM embeddings and normalized to yield interpretable positional weights. These weights are then used to emphasize functionally relevant positions when comparing anchor-query sequences. Importantly, weights are associated to all residues in an anchor sequence, in such a way to guarantee the emerging of possible residues as important even if MuLAN does not detect them as such. Formal definitions describing the MuLAN’s score normalisation are given in Supplementary Methods.

### Functional similarity metrics

To compare a query sequence with an anchor sequence, PLMView computes residue–residue similarity matrices from their PLM embeddings using both a radial basis function (RBF) kernel and cosine similarity (CS). MuLAN importance scores are used to weight the anchor positions. The resulting weighted matrices serve as scoring matrices in a fast sequence-alignment procedure with a gap penalty of zero. The aligned residue pairs are then aggregated into two complementary functional similarity measures: an RBF-based score, denoted *sim_RBF_*, and a cosine-based score, denoted *sim_CS_*. Detailed mathematical definitions of the similarity metrics and the anchor-query embedding alignment algorithm are provided in the Supplementary Methods, together with a discussion of the effect of setting the gap penalty to zero in the pairwise alignment algorithm.

### Hierarchical clustering, functional classification, and residue-level determinants

The multi-view similarity matrix is analyzed using the SCUT divisive hierarchical clustering algorithm^49^, which is well suited to high-dimensional data because it preserves both global and local structure. SCUT recursively partitions the PLMView functional space and produces an interpretable dendrogram in which subtrees correspond to functionally coherent sequence clusters. In PLMView, these clusters define the functional classification of the input sequences.

Importantly, because SCUT is recursive and graph-based, each split in the dendrogram can be associated with feature weights that trace back to specific anchors and residue-level similarity contributions. PLMView leverages this structure to decompose the functional placement of each sequence into residue-level importance scores, quantifying how strongly each position contributes to a given split or classification. By aggregating these scores across sequences within sister clusters, PLMView identifies residues whose contributions differ most strongly between subfamilies. These residues are interpreted as candidate functional determinants, providing a direct link between hierarchical functional classification and experimentally testable hypotheses about sequence positions driving functional divergence.

The formal definition of the functional score for a residue position within a multiple alignment of sequences in a cluster, the normalised functional score, and the maximum functional score for a residue position across different clusters are given in Supplementary Methods.

### Hierarchical clustering of comparison representations

Comparison dendrograms were reconstructed using agglomerative hierarchical clustering, with the linkage criterion selected according to the representation and comparison framework. For SSNs, average-linkage was applied to NBS-based distances, whereas single-linkage was used for e-value-based relationships. Dendrograms constructed directly from pooled sequence-level ESM2-3B and Ankh-Large embeddings were generated using Ward’s minimum-variance method, which iteratively merges clusters so as to minimize the increase in within-cluster variance in Euclidean embedding space. By contrast, dendrograms derived from averaged FANTASIA V1/ ProtT5 and FANTASIA V2/ProstT5 representations were reconstructed using Euclidean and cosine distances, preserving the pairwise ranking induced by the similarities, and single-linkage clustering. Ward, average-, and single-linkage were therefore used as alternative clustering criteria matched to the corresponding representation and comparison framework: single linkage for reconstructions based on threshold-defined relationships (see Supplementary Methods); average linkage when inter-cluster dissimilarity was defined by the mean pairwise distance between their members; and Ward’s method for direct clustering in the Euclidean space of pooled PLM embeddings.

### Construction of Sequence Similarity Networks

We constructed complementary BLAST-based SSNs using normalized bit scores (NBS) and e-values, with particular emphasis on the transformation of pairwise similarities into hierarchical representations.

For the NBS-based network, directed bit scores were normalized by query length and subsequently converted into a symmetric distance that also accounts for the self-similarity of each sequence pair; this distance was then clustered by average linkage to recover cohesive groups while reducing sensitivity to isolated high-scoring alignments. Unless otherwise stated, the SSN dendrograms shown in this study were derived from this NBS-based formulation. In parallel, e-value-based similarities were represented using single-linkage clustering, such that cutting the resulting dendrogram at a given threshold exactly recovers the connected components of the corresponding thresholded SSN. Very small or zero e-values were additionally transformed for visualization while preserving their similarity ordering.

The definitions of the two distance measures, the rationale for the linkage criteria, the treatment of multiple alignments and zero e-values, and the proof relating single-linkage dendrograms to SSN connected components are provided in the Supplementary Methods.

### Construction of phylogenetic trees

Phylogenetic trees for TRX (Fig. S4), opsins (Fig. 5B), and cold shock domain (Fig. S8) families were constructed using MAFFT^50,51^ and IQ-TREE2^52^. For opsins and cold-shock domain proteins, the best fitting substitution models were selected using ModelFinder^53^: LG+F+R7 for opsins, and LG+R9 for cold shock domain proteins. For TRX, sequences were aligned using MAFFT E-INS-i to accommodate large insertions, ModelFinder selected JTT+R8 as the best-fitting standard model. Refer to^28^ for the phylogenetic tree of the CPF family.

### Evaluation datasets

We evaluated PLMView across multiple protein families and experimental datasets that represent diverse biological contexts—from well-characterized enzymatic families to sensory and metabolic proteins with specialized functions. All datasets of sequences and their functional annotation is available at https://zenodo.org/records/20555097

### Plant Thioredoxin family (TRX)

We retrieved all UniProt sequences containing the thioredoxin domain (PF00085) and taxonomically annotated as Viridiplantae (taxonomy id: 33090), in total 30516 sequences. After filtering out sequences with length *>*3000aa, 30513 sequences remained.

### Visual physiology opsin database

The Visual Physiology Opsin Database (VPOD)^54^ (https://github.com/VisualPhysiologyDB/ visual-physiology-opsin-db/tree/main/vpod_data/VPOD_1.3) provides 364 wild-type visual opsin sequences with experimentally measured absorption maxima (*λ_max_*), reflecting the wavelength of peak photon absorption. These sequences are phylogenetically and functionally annotated and were used to assess PLMView’s ability to capture sequence determinants of spectral tuning, a key example of functional specialization at the residue level.

### Cold-shock proteins from Tara Oceans data and preprocessing

From the *Tara* Oceans Expeditions sampled MetaG and MetaT data, we retrieved diatom unigenes containing the Cold-shock DNA binding domain (PF00313). After filtering and data cleaning, we obtained 3287 domain sequences.

In this preprocessing step, Metagenome-Assembled Genomes (MAGs)^55^(downloaded at https://www.genoscope.cns.fr/tara/#SMAGs) from diatoms were annotated using MetaClade2^41^ MAG genes containing cold-shock DNA-binding domains were then used as queries against the full *Tara* Oceans diatom unigene catalog^29^ (https://www.genoscope.cns.fr/tara/#MATOU-1.5). Candidate matches were first identified using tblastn-evalue 1e-1^56^, followed by refinement with the exonerate tool^57^ to recover the longest coding sequences, which were subsequently translated into amino acid sequences.

The resulting protein sequences were annotated again with MetaClade2 to identify domain boundaries. For each sequence, only the longest domain hit was retained, provided that it satisfied the following criteria: e-value *≥* 1e-2, coverage *≥* 0.8, and domain length within 10% of the reference cold-shock domain length (64 amino acids). In addition, each retained domain was required to contain at least one hit with e-value *≥* 1e-8.

### Cryptochrome/Photolyase Family (CPF)

To compare PLMView with other computational approaches to functional classification, we used the CPF family, consisting of 307 protein sequences retrieved from the curated set described in^28^ (www.lcqb.upmc.fr/profileview/documentation/cpf-usecase/). The sequences originate from UniProt, JGI genome projects (genome.jgi.doe.gov), and OIST marine genomics projects (marinegenomics.oist.jp). This dataset was designed to combine functional characterization with broad taxonomic coverage. It includes CPF sequences for which specific functions have been assigned based on experimental evidence reported in the literature, together with sequences selected to span the diversity of the tree of life. Overall, the dataset covers 146 species, 74 classes, and 40 phyla. All sequences contain the FAD-binding cryptochrome/photolyase domain PF03441.

Because CPF proteins can differ in length and several sequences in the dataset are fragmented, we based the PLMView analysis on the conserved FAD-binding domain. Anchor sequences were trimmed to the corresponding PF03441 domain boundaries, whereas the sequences to be classified were analysed at full length. Domain boundaries were identified by aligning all sequences with MAFFT G-INS-i and aligning the resulting multiple sequence alignment to the Pfam HMM profile PF03441. Columns 1 to 189 (included) of the HMM alignment were retained for the domain-level representation of the anchors. For the SSN comparisons, CPF sequence relationships were reconstructed using the same sequence set, allowing direct comparison with the PLMView and pooled-embedding classifications.

### Rapid scalability and computational efficiency

A central design goal of PLMView is computational efficiency. The framework separates the computationally intensive generation of PLM embeddings from the comparatively lightweight construction and clustering of the collaborative functional space. For a family comprising 8,022 proteins with a mean length of 296 amino acids and 1,006 anchors, the complete workflow produced the functional tree in 28 min 41 s using Ankh embeddings^58^ computed in fp32. Embedding generation accounted for 17 min 22 s of this runtime, whereas the subsequent PLMView analysis required approximately 11 min.

Using bfloat16 (bf16) reduced the total runtime to 16 min 14 s, primarily by decreasing embedding-generation time to 4 min 50 s; the duration of the downstream analysis remained essentially unchanged. The resulting functional trees showed no substantial differences from those obtained with fp32 embeddings. Thus, PLM inference represents the largest single computational cost, while a comparable amount of time is collectively required for the collaborative comparisons, construction of the functional space, and hierarchical clustering.

The analyses were ran on an computer with an AMD Ryzen Threadripper PRO 5955WX 16-Cores CPU and NVIDIA RTX A4000 GPU (16GB VRAM).

### Hardware acceleration

To enable large-scale analyses, the computationally intensive components of PLMView were optimized for both CPU and GPU execution. PLM sequence embedding generation was handled using native PyTorch^59^/HuggingFace^60^ PLM implementation. A Cython^61^ implementation accelerates the core sequence–embedding alignment on CPUs, while a CUDA^62^ implementation, enables massively parallel computation on GPUs.

The CUDA version employs a parallelized Needleman–Wunsch algorithm similar to previous GPU formulations^63,64^, but adapted to operate directly on high-dimensional PLM embeddings. The GPU implementation achieves up to a 10× speed-up compared to the sequential Cython version—approximately 5× from CUDA parallelization and an additional 2× from half-precision (16-bit) computation.

In practice, the principal memory constraint stems from the embedding generation step of the PLMs rather than the downstream alignment or clustering stages.

In a separate benchmark, PLMView processed 10,011,456 comparisons between 8,022 GH2 sequences and 1,248 anchors in 807 s, compared to 2,784 s for hmmsearch (HMMER3) on a single CPU core and 495–709 s on 8–4 cores, respectively. Although PLMView was approximately 6–7× slower than HH-suite3’s hhblits on a single core, it achieved this performance while aligning high-dimensional sequence embeddings without prefiltering.

Once embeddings are available, PLMView can therefore analyse thousands of homologous proteins within minutes, enabling iterative exploration of protein families and large-scale functional surveys without sacrificing fine-grained resolution. This computational profile makes the framework applicable to the rapidly expanding sequence collections generated by genomic, metagenomic, and metatranscriptomic studies.

## RESOURCE AVAILABILITY

The datasets used for evaluation were obtained from publicly available resources. All data generated by the PLMView analyses are available at https://zenodo.org/records/20555097. The PLMView code is available at https://gitlab.lcqb.upmc.fr/Vinh-Son/PLMView under the CC BY-NC-SA 4.0 licence.

## ACKNOWLEDGMENTS

We thank Juan José Pierella Karlusich for advice on the use of *Tara* Oceans data; Julien Henri for assistance in interpreting thioredoxin functional determinants; Seth A. Frazer for helpful discussions concerning the VPOD dataset; Maya Czeneszew for her valuable suggestions on the manuscript.

## AUTHOR CONTRIBUTIONS

Conceptualization, VSP and AC; methodology, VSP and AC; software implementation, VSP; testing, VSP and ANB; data analysis, VSP, ANB, MS, CB and AC; writing, VSP and AC; editing, VSP, ANB, CB and AC; funding acquisition, AC; supervision, AC.

## FUNDS

This work was performed using HPC resources from GENCI–IDRIS and Sorbonne University GPU clusters from SCAI and the LIP6 laboratory (UMR 7606, Sorbonne University-CNRS). Financial support from PostGenAI@Paris within the framework of France 2030, ANR-23-IACL-0007 (AC); ANR DEFINE ANR-24-CE45-7686 (AC); Institut Universitaire de France (AC); PhD fellowship from Ministère de l’Enseignement Supérieur et de la Recherche, Sorbonne University (VSP); Horizon Europe project BlueRemediomics, grant agreement number 101082304 (CB).

## DECLARATION OF INTERESTS

The authors declare no competing interests.

## Supplementary Information

## Supplementary Methods

### Anchor selection

**Table S1:** Easy-cluster hyperparameters.

| Dataset | Dataset characteristics |  | Parameters |  |
| --- | --- | --- | --- | --- |
|  | # of input sequences | # of anchors | -c | --min-seq-id |
| TRX | 30,513 | 3,064 | 0.5 | 0.6 |
| TRX - trimmed | 31,080 | 1,289 | 0.5 | 0.6 |
| VPOD | 364 | 203 | 0.8 | 0.95 |
| Cold shock domain PF00313 | 3,287 | 1,356 | 0.5 | 0.9 |
| CPF | 307 | 307 (trimmed) | 1.0 | 1.0 |

Given a set of homologous sequences *B* to classify, PLMView first selects a subset of representative sequences *A ⊂ B*, referred to as anchors. Anchors are identified using MMseqs2^65^, specifically the easy-cluster procedure, typically with the default parameters -c 0.8, which sets the minimum coverage, and --min-seq-id 0.6, which sets the minimum sequence identity within clusters.

The anchor set defines a reduced, sequence-diverse reference space that spans the variability of the protein family while substantially limiting the number of pairwise comparisons required during downstream analysis. This strategy reduces computational complexity while preserving the sequence variation necessary for fine-grained functional discrimination. Anchors do not need to originate from the target dataset itself and may instead consist of an external collection of representative homologous sequences defining a more general reference functional space.

### Guidelines for anchor selection

The clustering thresholds can be adjusted according to the length, completeness, and diversity of the sequences under study. The coverage parameter -c controls the minimum fraction of a sequence that must be included in a match. Lower values provide greater tolerance for short domains, partial proteins, and fragmented metagenomic sequences. Accordingly, we used -c 0.5 for the TRX and cold-shock domain datasets, whereas the default value of -c 0.8 is generally suitable for complete protein sequences, like for the VPOD sequences.

The parameter --min-seq-id primarily controls the number of clusters and, consequently, the number of anchors. Lower values merge more sequences into the same cluster and produce a smaller anchor set, thereby reducing runtime. Higher values retain more representatives and may improve resolution in closely related or functionally diverse families. As a practical guideline, for datasets containing several thousand sequences, the anchor set can generally be kept substantially smaller than the full dataset, often representing only a few percent to roughly one tenth of the sequences, depending on family diversity and the desired resolution. For smaller datasets, all or most sequences may instead be retained as anchors. We used --min-seq-id 0.95 for the VPOD dataset and --min-seq-id 1.0 for the CPF dataset.

For small datasets, clustering may be omitted and all sequences used directly as anchors. This strategy was feasible for the CPF family, which comprised 307 sequences. Because all sequences contained the FAD-binding domain but not necessarily the photolyase-related domain region, anchors were trimmed to the FAD-binding domain, and each full-length query sequence was compared with all anchor FAD regions.

### MuLAN’s score normalisation

PLMView uses residue-level importance scores derived from MuLAN attention maps to emphasize functionally informative regions of anchor sequences. Let *C*_1_ and *C*_2_ denote the outputs of the MuLAN convolutional layers applied to the residue embeddings of a sequence of length *L*.

The raw MuLAN score vector *rm* is defined as

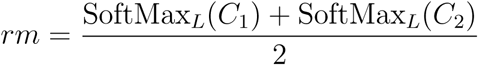

where SoftMax*_L_* denotes the softmax operation over sequence positions.

To obtain interpretable positional weights independent of sequence length, we define normalized MuLAN scores *m* as:

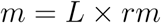

Consequently:

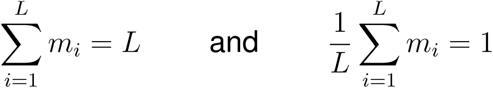

The normalized MuLAN score therefore has unit average across the sequence, allowing residues with scores above 1 to be interpreted as positions of above-average functional importance.

Intuitively, MuLAN scores identify regions likely involved in molecular interactions or functional specialization. In PLMView, these scores are used to weight residues in anchor sequences so that functionally informative regions contribute more strongly to sequence comparisons.

### Functional similarity functions

To compare a query sequence *B ∈ B* with an anchor sequence *A ∈ A*, PLMView computes residue-level similarities between their PLM embeddings. Let 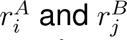 denote the embeddings of residue *i* in *A* and residue *j* in *B*, respectively. We use two complementary similarity functions:

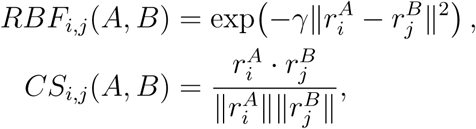

where *γ >* 0 controls the scale of the radial basis function and ||·|| denotes the Euclidean norm. In our implementation, *γ* = 1, unless otherwise specified. These two functions capture distinct geometrical properties of the embedding space. Cosine similarity (CS) measures angular agreement between embeddings and is therefore insensitive to their norms. It captures whether two residues point in similar directions in representation space, a property that has been related to structural similarity in PLM embeddings^24,48^. In contrast, the RBF similarity is distance-based: it assigns high similarity only to residues that are close in Euclidean distance and decreases exponentially as they move apart. Thus, while cosine similarity captures directional relationships, the RBF kernel captures local neighborhoods around residues in embedding space^22^.

To emphasize residues of the anchor sequence that are expected to be functionally informative, both similarity matrices are weighted by the MuLAN scores of the anchor. Let *m^A^* denote the MuLAN score of residue *i* in anchor *A*, and let

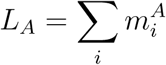

be the effective weighted length of the anchor. The residue-level alignment score matrix *SM* is defined as

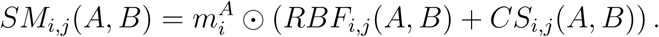

where *⊙* denotes the element-wise product. The MuLAN weights act row-wise on the anchor residues, increasing the contribution of positions predicted to be functionally relevant.

The matrix *SM* (*A, B*) is then used as an alignment cost matrix. Sequence alignment is performed by dynamic programming with zero gap penalty, so that insertions and deletions do not contribute to the score. Under this condition, Needleman–Wunsch and Smith–Waterman become equivalent up to free end gaps, since all insertions and deletions carry no penalty and alignment scores are driven entirely by residue similarity. The recursive dynamic programming step is therefore

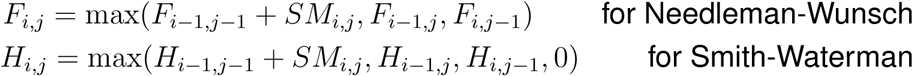

with *F_i,_*_0_ = 0, *F*_0*,j*_ = 0 and *H_i,_*_0_ = 0, *H*_0*,j*_ = 0 respectively.

Because PLMView only requires the final alignment score and not the explicit alignment path, no backtracking step is performed, reducing computational cost.

Using the resulting alignment, PLMView computes two sequence-level similarity scores between anchor *A* and query *B*:

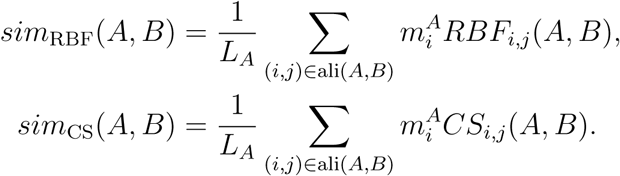

Thus, each anchor contributes two coordinates to the PLMView representation of a query sequence: one radial coordinate and one angular coordinate. For a set of anchors *A* = *{A*_1_*,…, A_N_ }*, the functional representation of *B* is therefore

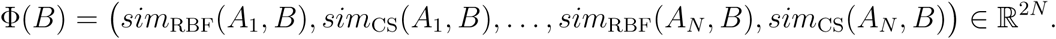

Using both similarity functions is important because they define complementary views of functional relatedness. Cosine similarity provides an interpretable angular comparison: for a fixed query embedding *x*, a weighted combination of cosine similarities to anchors *a_i_* satisfies

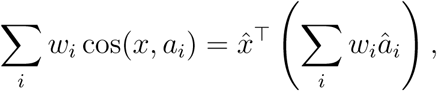

where *x̂* and *â_i_* denote normalized vectors. When the weighted anchor sum is non-zero, this is equivalent, up to a positive scaling factor, to a single cosine similarity between *x* and an effective anchor direction. Therefore, the cosine component captures angular trends and, when the weights are non-negative, defines cone-like regions generated by anchor directions. This geometry is efficient and interpretable, but it is essentially directional.

The RBF component complements this angular view by introducing a non-linear, distance-dependent notion of similarity. It can distinguish residues or sequences that have similar directions but lie at different distances in embedding space, and it can capture localized neighborhoods that are not well represented by a single effective direction. Combining cosine and RBF similarities therefore allows PLMView to retain the robustness and interpretability of angular comparisons while also detecting local, non-linear structure in the embedding space. Together, the two measures provide a richer functional space than either measure alone, with each query sequence positioned relative to anchors through both directional and distance-based relationships.

### On Needleman–Wunsch versus Smith–Waterman pairwise alignment

In PLMView, pairwise alignment is used only to compute an optimal residue-level similarity score between an anchor sequence and a query sequence; the explicit alignment path is not required. Since the gap penalty is set to zero, insertions and deletions do not decrease the alignment score. As a consequence, terminal gaps are effectively free, and any unfavorable residue match can be bypassed without penalty.

Under these conditions, Needleman–Wunsch and Smith–Waterman yield the same optimal score for our purposes. Although Smith–Waterman is usually formulated as a local alignment algorithm and Needleman–Wunsch as a global alignment algorithm, the distinction disappears here because zero-cost gaps allow the score of any optimal local match to be propagated to the full sequence boundaries without loss. Conversely, the global alignment is not forced to accumulate negative contributions, since such positions can be skipped through zero-cost gaps. This point is relevant because the PLMView similarity matrix includes cosine similarity, which can in principle introduce negative entries. However, negative values do not break the equivalence in this setting: with zero gap cost, they are simply avoided when they would reduce the score. Empirically, across millions of sequence pairs, the combined alignment matrix remained strongly dominated by positive values, with a minimum of *−*0.08, a maximum of 3.19, and a mean of 0.3. Thus, in the PLMView setting, Needleman–Wunsch and Smith–Waterman are effectively equivalent, and the choice of either algorithm does not affect the resulting sequence-level similarity scores.

### Hierarchical clustering

For every anchor–sequence pair (*A, B*) *∈ A×B*, PLMView computes the two similarity measures *sim_RBF_* and *sim_CS_*, yielding matrices

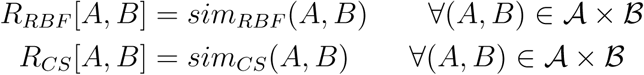

The final multi-view representation is obtained by concatenation:

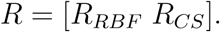

Each sequence (row) is therefore represented by its similarities to all anchors (columns) across multiple embedding-derived views. This representation is independent of sequence length and encodes the functional position of a sequence relative to the anchor set.

Hierarchical organization of the sequences is then reconstructed using the divisive clustering framework SCUT^49^. SCUT was chosen because it preserves both global and local organization of the data while providing an interpretable hierarchical partitioning. In this setting, the matrix *R* can be interpreted as the weighted adjacency matrix of a bipartite graph linking sequences and anchors, matching the SCUT formulation.

Because the hierarchy is constructed recursively through successive binary splits, each partition is directly associated with the features contributing most strongly to the separation, enabling downstream interpretation of functional determinants.

### Extraction of functional determinants

PLMView enables residue-level interpretation of functional partitions by decomposing the placement of a sequence within the hierarchical functional space into contributions from individual residues.

At each node *T* of the hierarchy, sequences are separated by a linear projection in the multi-view similarity space. Let *p_T_* (*B*) denote the projection score associated with sequence *B* at node *T*:

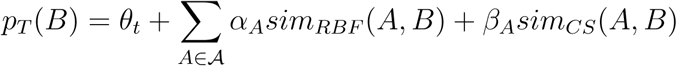

where *A* is the set of anchors, *α_A_* and *β_A_* are anchor-specific coefficients, and *θ_T_* is the node-specific offset.

The coefficients *α_A_* and *β_A_* are learned during the recursive clustering procedure defining the split at node *T*. Operationally, the clustering algorithm determines a separating hyperplane satisfying *p_T_* (*B*) = 0, while *θ_T_* corresponds to the low-density region separating the two daughter subtrees.

Because both similarity measures are computed as sums over aligned residue pairs, the projection function itself can be decomposed into residue-level contributions.

For the first similarity metric,

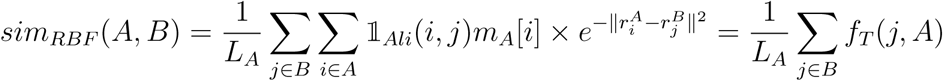

where 1*_Ali_*(*i, j*) = 1 when residues *i* and *j* are aligned and 0 otherwise, *r_i_* and *r_j_* denote residue embeddings, and *ms_A_*[*i*] is the normalized MuLAN weight associated with residue *i* of anchor *A*. Note that *f_T_* (*j, A*) quantifies the contribution of residue *j* in sequence *B* to the similarity between *A* and *B* for the first metric.

Similarly, for the cosine similarity metric,

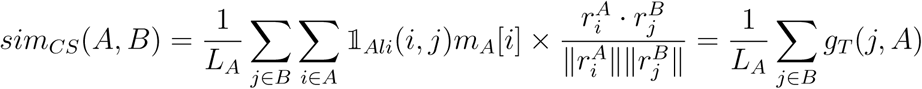

where the summing term over all positions of *A* is renamed *g_T_* (*j, A*). Substituting these expressions into the projection function yields

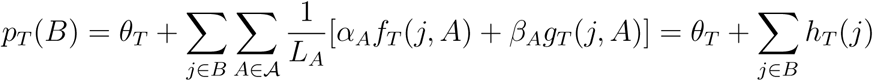

where the quantity *h_T_* (*j*) measures how strongly residue *j* contributes to the placement of sequence *B* within the functional split defined at node *T*. Positive and negative contributions respectively favor one or the other daughter subtree. In this way, the geometric organization of sequences in PLMView functional space can be directly interpreted in terms of residue-level contributions.

### Functional score and maximum functional score

Residue-level contributions can be aggregated across sequences belonging to sister subtrees in order to identify residues associated with functional divergence at a given bifurcation of the hierarchy.

Consider a node *T* with left and right daughter subtrees denoted *T_L_* and *T_R_*. To characterize the functional split associated with *T*, we sample representative sequences from both subtrees and construct a multiple sequence alignment, denoted *MSA_T_*, over the combined set of sampled sequences. In the case studies analyzed in this work, approximately 64 sequences were sampled from each of *T_L_* and *T_R_*. This sampling strategy limits alignment size while preserving representative sequence diversity within each subtree. When a set *E* of subtrees was analyzed jointly, the sampled sequences from all corresponding subtree analyses were pooled to construct the alignment *MSA_E_*.

For each alignment position *j* in *MSA_T_*, we compute the average residue contribution within each subtree of *T*:

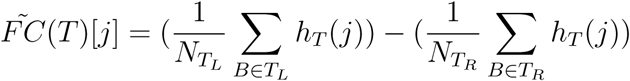

where *N_TL_* and *N_TR_* denote the numbers of sampled sequences in the left and right subtrees of *T*, respectively. The quantity *F̃C*(*T*)[*j*] therefore measures how differently alignment position *j* contributes to the placement of sequences in the two sister subtrees. Positions with large absolute values correspond to residues whose embedding contributions strongly discriminate the two functional groups. Gapped positions contribute zero to the corresponding terms in the alignment averages.

To facilitate comparisons across splits, we normalize the positional scores

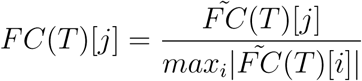

where the normalization runs over all alignment positions of *MSA_T_*. By construction, *FC*(*T*)[*j*] *∈* [*−*1, 1].

The normalized functional score *FC*(*T*)[*j*] therefore quantifies both the strength and direction of the contribution of alignment position *j* to the functional bifurcation defined by node *T*.

Functional determinants can be analyzed across multiple hierarchical levels of a PLMView tree simultaneously. Given an ensemble of nodes *E* corresponding to different functional splits in the hierarchy, we define the maximum functional score:

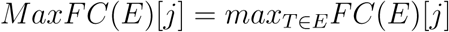

The quantity *MaxFC*(*E*)[*j*] identifies alignment positions exhibiting strong functional contributions across one or several hierarchical scales. This enables reconstruction of residue-level functional determinants progressively refined from broad family-level partitions to highly specific sequence specializations.

### Ankh-Large vs ESM2-3B comparison

Because PLMView relies on residue-level embedding similarities to infer functional relationships between homologous proteins, we sought to characterize the geometric and statistical properties of the embedding spaces produced by different PLMs.

We generated residue-level embeddings using Ankh-Large and ESM2-3B for triplets of sequences sampled from 11,950 Pfam domains. For each domain, we computed pairwise similarity matrices for the three distinct sequence pairs using both cosine similarity *CS* and radial basis function similarity *RBF*. Reported values correspond to averages over all 3 *×* 11,950 sequence pairs.

Two notable properties emerge from this analysis. First, ESM2-3B exhibits a surprisingly high average cosine similarity between residue embeddings (*∼* 0.75), even when compared residues do not correspond to aligned or functionally equivalent positions (see panels AB in the illustration below). This suggests the presence of strong global embedding components that reduce contrast between residue representations. Indeed, the effect is substantially reduced after subtracting the mean pooled embedding from each residue embedding (panel C). Such behavior may arise from attention-induced token mixing or from anisotropy in the ESM2-3B embedding space, whereby embeddings are concentrated along a limited number of dominant directions, artificially increasing cosine similarity between otherwise unrelated residues. However, because pooled embeddings are known to retain biologically meaningful functional information^13–15^, we chose not to center the embeddings in subsequent analyses. Second, the average *RBF* similarity for ESM2-3B is close to zero, indicating that Euclidean distances between residue embeddings remain comparatively large despite their high cosine similarity (panels DE). Neither of these behaviors is observed for Ankh-Large, whose embeddings display both lower global cosine similarity and more balanced distance distributions.

**Figure S1:**
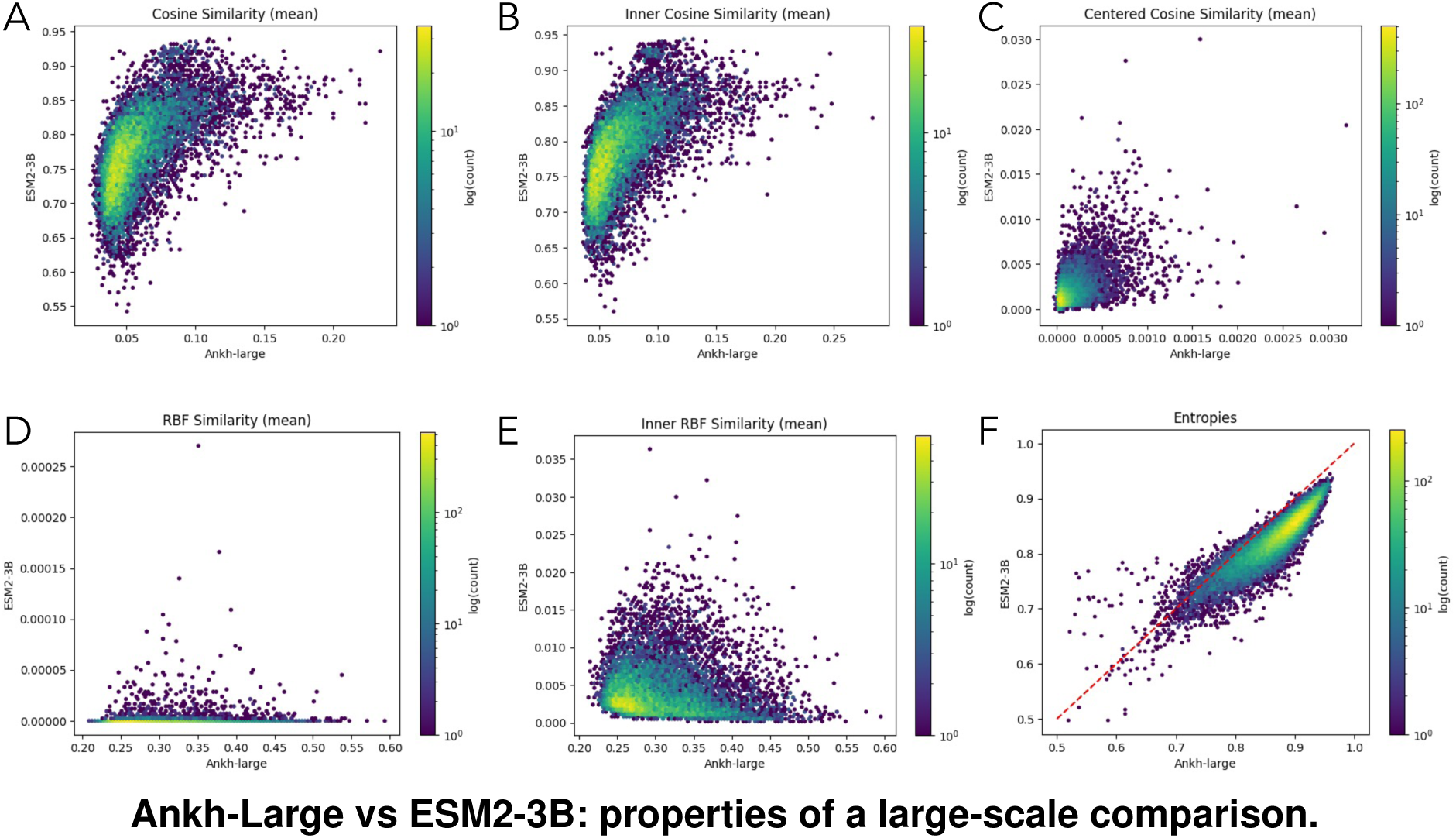
The plots contain 11,950 points corresponding to different Pfam domains (see text). They compare ESM2-3B (y-axis) vs Ankh-Large (x-axis) scores averaged over 3 different sequences chosen to represent each of the 11,950 domains. Comparison is based on: A. cosine similarity; B. inner cosine similarity; C. centered cosine similarity; D. RBF similarity; E. inner RBF similarity; F. Normalized effective rank (entropy-based).

To further characterize how information is distributed across embedding dimensions, we performed singular value decomposition (SVD) on centered residue embeddings for all previously sampled sequences. Squared singular values were interpreted as an explained-variance distribution, analogous to PCA, and the entropy of this distribution was computed^66^, and divided by the maximum achievable entropy. We found that ESM2-3B embeddings concentrate explained variance in relatively few dimensions, whereas Ankh-Large embeddings exhibit a substantially broader and more even variance distribution (panel F). This trend was highly consistent across Pfam domains: Ankh-Large showed higher explained-variance entropy than ESM2-3B in 98% of the 11,950 domains considered. Although a broader variance distribution could in principle indicate noisier embeddings, the strong correlation of Ankh-Large embeddings with protein structure^48^ instead suggests that they encode richer and more distributed representations.

### Construction of Sequence Similarity Networks

We constructed two types of Sequence Similarity Networks (SSNs): one based on normalized bitscores (NBS) and one based on BLAST e-values. These two measures capture complementary aspects of sequence similarity. The NBS-based network relies on an information-theoretic measure derived from pairwise sequence alignments, whereas the e-value-based network uses the statistical significance of the alignment. In both cases, networks were reconstructed using blastp, with protein sequences represented as nodes and weighted edges encoding pairwise sequence similarity.

For the NBS-based SSN, the weight of the directed edge from sequence *i* to sequence *j* was defined as

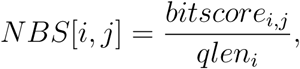

where *bitscore_i,j_* is the BLAST bit score obtained when sequence *i* is used as the query against sequence *j*, and *qlen_i_* is the length of the query sequence. When multiple alignments were detected between the same pair of sequences, only the alignment with the highest bit score was retained.

The directed NBS matrix was then converted into a symmetric distance matrix *D*:

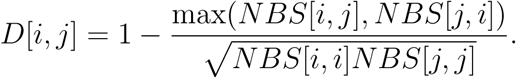

The denominator normalizes pairwise similarities by the self-similarity scores of the two sequences, so that higher sequence similarity corresponds to shorter distances.

The resulting NBS distance matrix *D* was clustered using average-linkage hierarchical clustering, also known as UPGMA. This choice summarizes the continuous sequence-similarity relationships encoded by the NBS matrix: clusters are merged according to their average pairwise distance, so that groups of sequences are joined when they are similar as a whole. Compared with linkage criteria driven by individual sequence pairs, average linkage is less sensitive to isolated high-scoring alignments and favors more cohesive sequence groups. The NBS-based dendrograms therefore provide a hierarchical visualization of the global similarity structure of the protein family. Unless otherwise stated, the SSN dendrograms presented in this article were constructed from this NBS-derived distance matrix.

The second type of SSN was constructed from BLAST e-values, following^67^. Each sequence set was compared against itself using blastp. To retain weak pairwise similarities and avoid truncation of the hit list, we used -max target seqs 1000000 and -evalue 10000. We then defined a symmetric e-value-based distance matrix *D̃* as

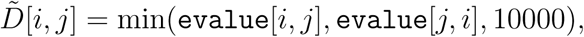

with *D̃*[*i, i*] = 0.

Because e-value thresholds are commonly used to define connected components in SSNs, the e-value-based matrix was clustered using single-linkage hierarchical clustering. This choice provides a hierarchical representation of the connected components obtained by varying the e-value threshold: at any given threshold, cutting the single-linkage dendrogram recovers the corresponding SSN connected components. A proof of this equivalence is provided in the Supplementary Text.

For visualization of the e-value-based dendrograms, very small e-values reported by blastp as zero were transformed as

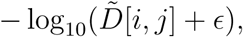

where *ɛ* was chosen as the smallest positive terminal branch length in the dendrogram, or as 10^−180^ when no positive value was available. This transformation avoids undefined values while preserving the ordering of sequence similarities.

These two linkage choices reflect two complementary uses of SSNs: average linkage provides a hierarchy of cohesive sequence groups from continuous similarity scores, whereas single linkage provides an exact hierarchical representation of threshold-based connected components. The former is closer in spirit to community-detection approaches such as Markov Clustering, although it should not be interpreted as formally equivalent to them.

### Equivalence between single-linkage clustering and thresholding a sequence similarity network

Let *G* = (*V, E, w*) be a connected edge-weighted graph representing a sequence similarity network, where *w*(*e*) denotes the dissimilarity associated with edge *e*. For a threshold *t*, define the thresholded graph

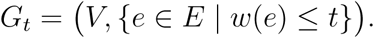

The connected components of *G_t_* define the partition of the network obtained at threshold *t*.

Let *L* = (*e*_1_*, e*_2_*,…, e_m_*), with *m* = *|E|*, be the edges of *G* sorted in non-decreasing order of dissimilarity. Increasing the threshold *t* from small to large is equivalent to starting from the graph (*V, ∅*) and adding the edges of *L* one after another. When an edge is added, two cases are possible: either its endpoints lie in two distinct connected components, in which case the edge merges these components and changes the induced partition; or its endpoints already belong to the same component, in which case the partition remains unchanged. Thus, only edges connecting distinct connected components affect the evolution of the threshold-induced partitions.

Single-linkage clustering defines the dissimilarity between two clusters *C* and *C^′^* as

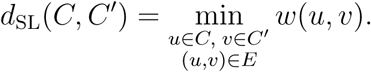

At each step, single linkage merges the two clusters with the smallest inter-cluster dissimilarity. In graph terms, this is exactly the operation of adding the smallest-weight edge whose endpoints belong to two distinct connected components. Edges whose endpoints already lie in the same component are ignored, since they do not change the current partition.

Therefore, single-linkage clustering follows the same sequence of partition changes as the thresholding process applied to *G*. Equivalently, the single-linkage dendrogram encodes the connected-component structure of the sequence similarity network for every possible threshold *t*. Cutting the dendrogram at height *t* yields the same partition as taking the connected components of *G_t_*.

## Supplementary Figures

**Figure S2:**
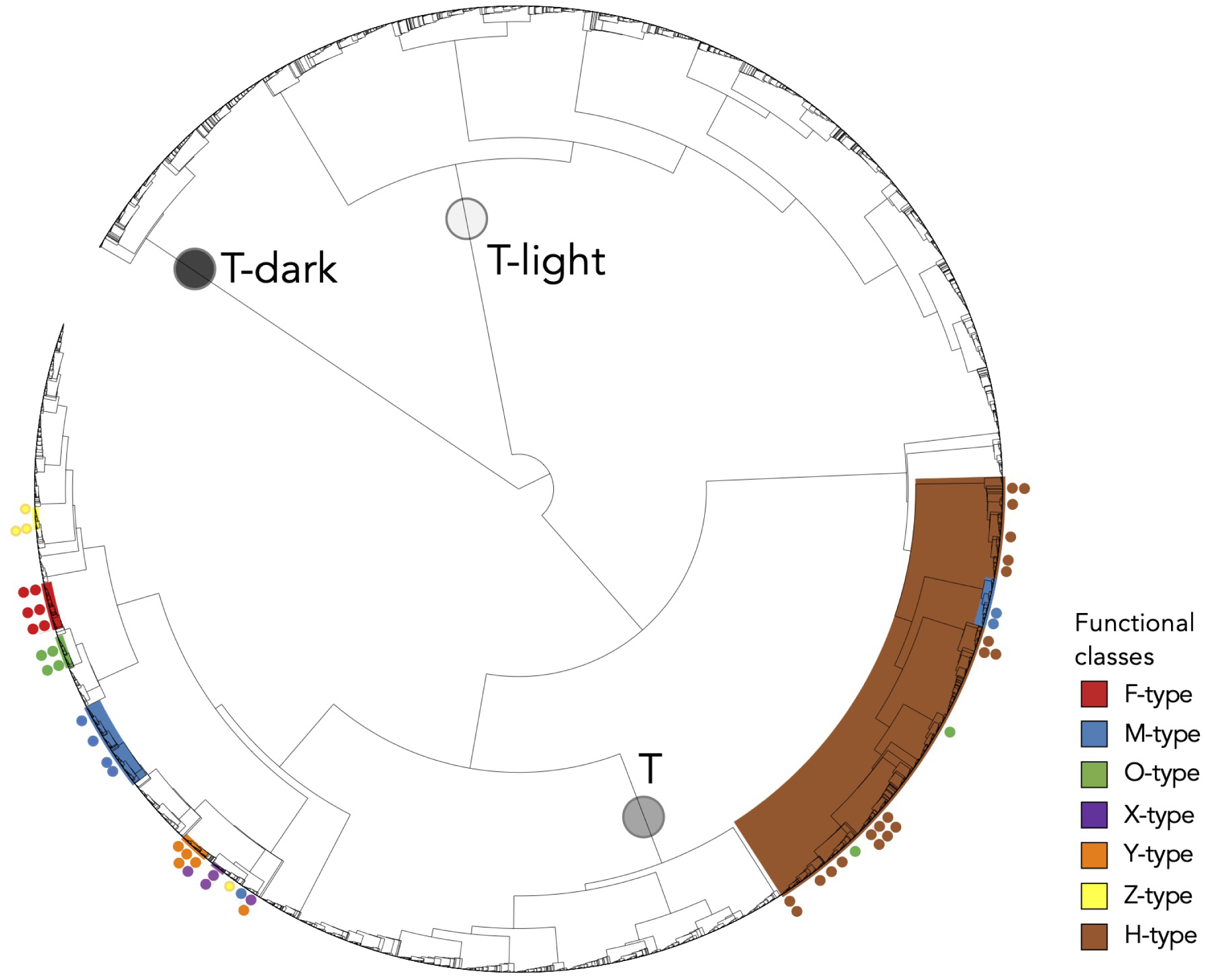
Large view of Fig. 4A. Grey circles (dark grey, grey, light grey) on the branches of the tree highlight distinct subtrees (T-dark, T, T-light, respectively) where no functional annotation is available. The same coloring is used to identify sequences in the corresponding phylogenetic tree of Fig. S4.

**Figure S3:**
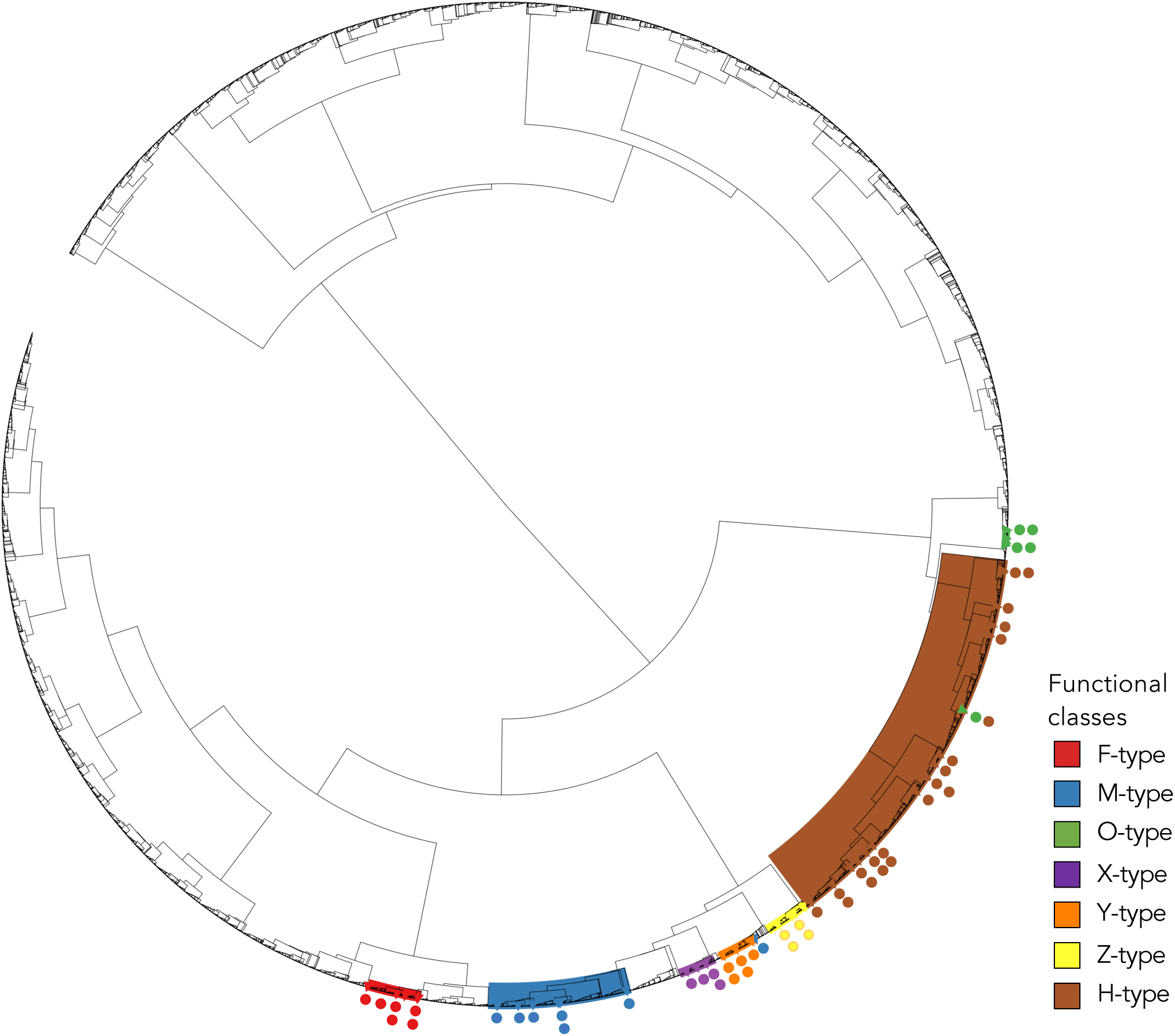
TRX PLMView tree based on trimmed TRX sequences. PLMView tree for sequences in Fig. 4A where the transit peptide subsequences, when present, have been trimmed off.

**Figure S4:**
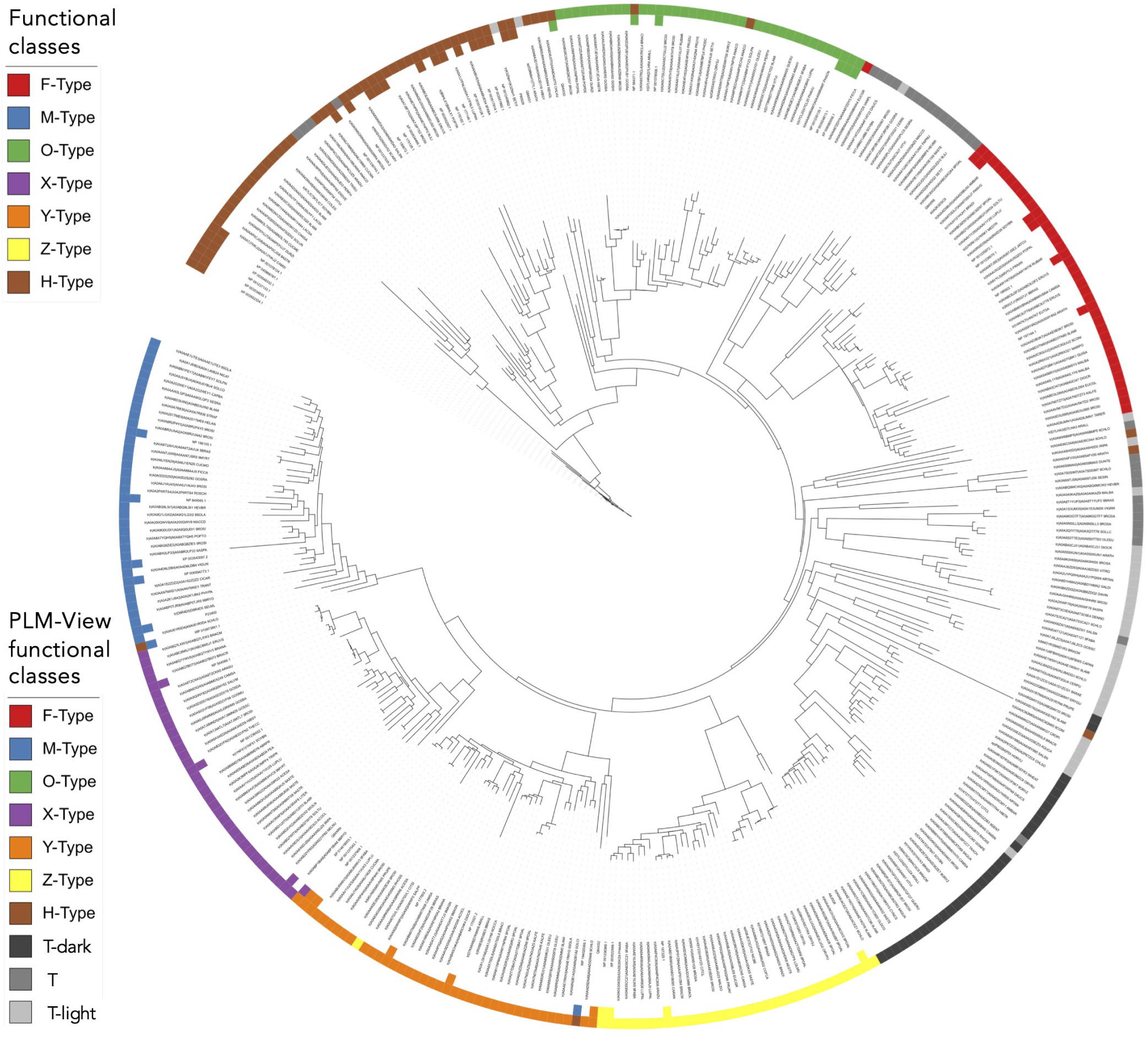
Phylogenetic tree of TRX sequences. A sample of 372 TRX sequences occurring in the PLMView functional tree in Fig. 4A is organised in a phylogenetic tree. The outer circle colours sequences based on PLMView functional partition as in Fig. 4A (bottom inset legend). Grey colors are explained in Fig. S2. The inner circle colors sequences with experimental functional characterization (top inset legend).

**Figure S5:**
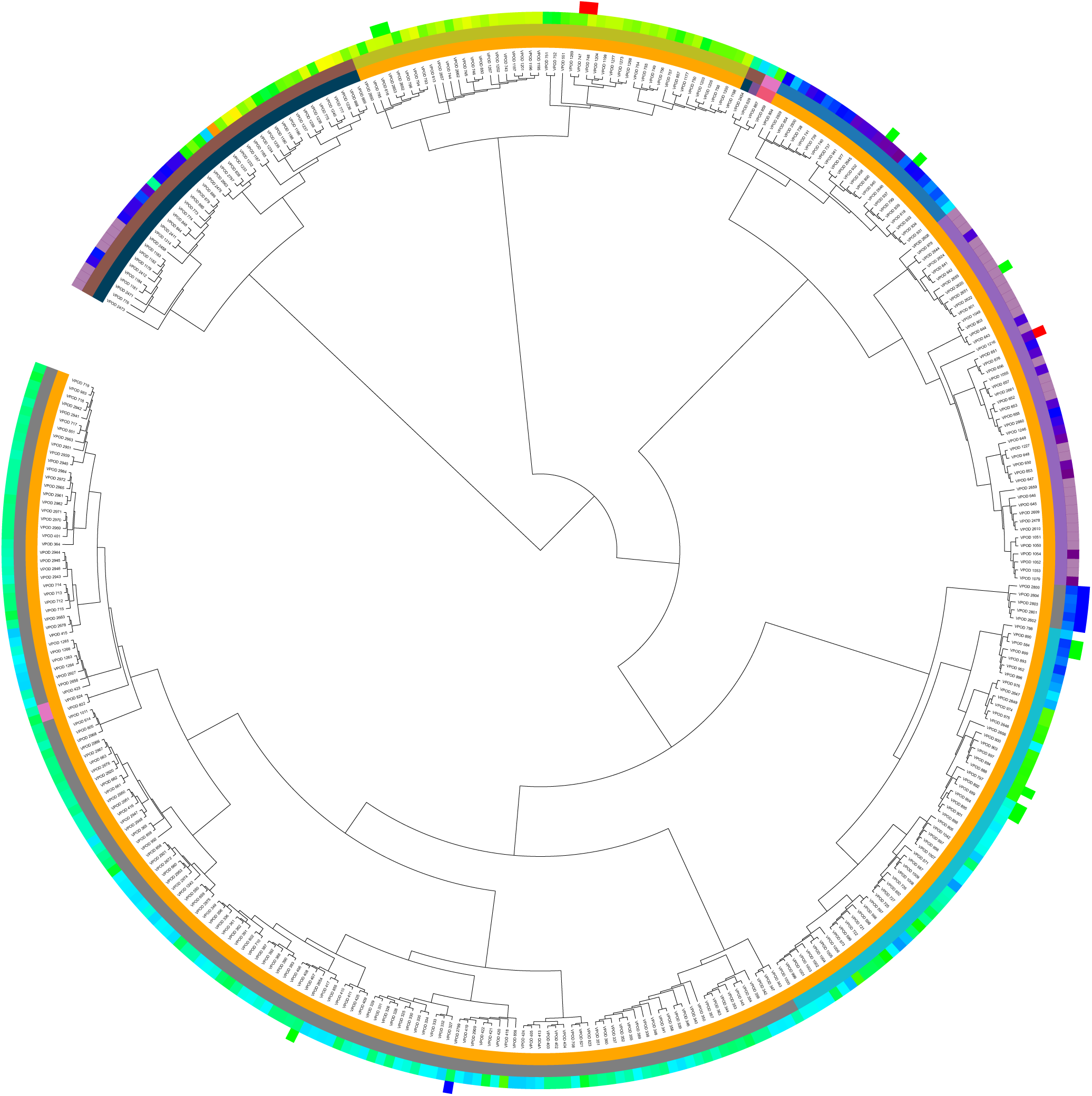
PLMView tree of the visual opsin sequences. Enlarged view of Fig. 5A. Color legend as in Fig. 5AB.

**Figure S6:**
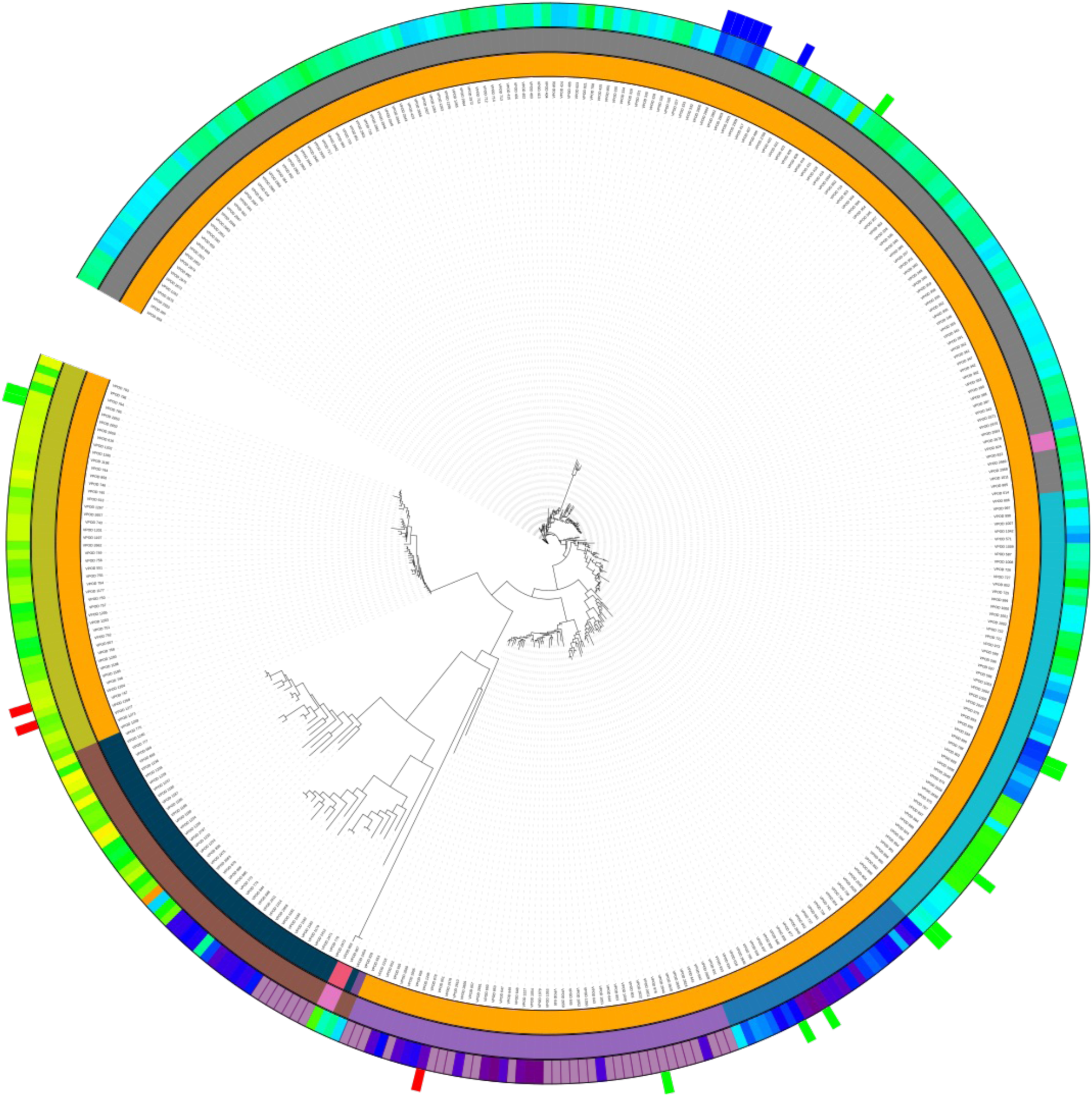
Phylogenetic tree of the visual opsin sequences. Enlarged view of Fig. 5B. Color legend as in Fig. 5AB.

**Figure S7:**
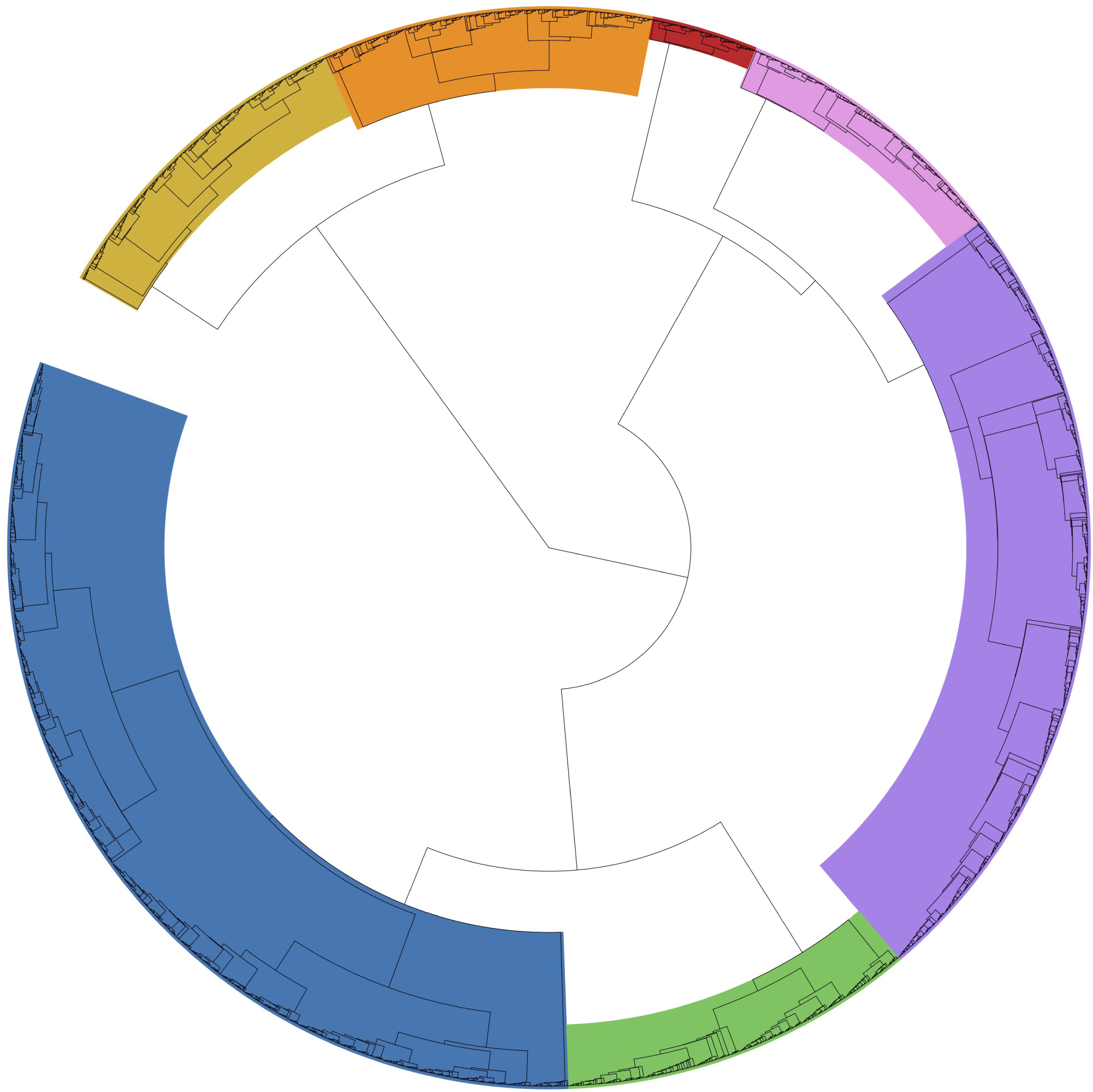
PLMView tree of the cold-shock DNA binding domain sequences. Enlarged view of Fig. 6B. Color legend as in Fig. 6B.

**Figure S8:**
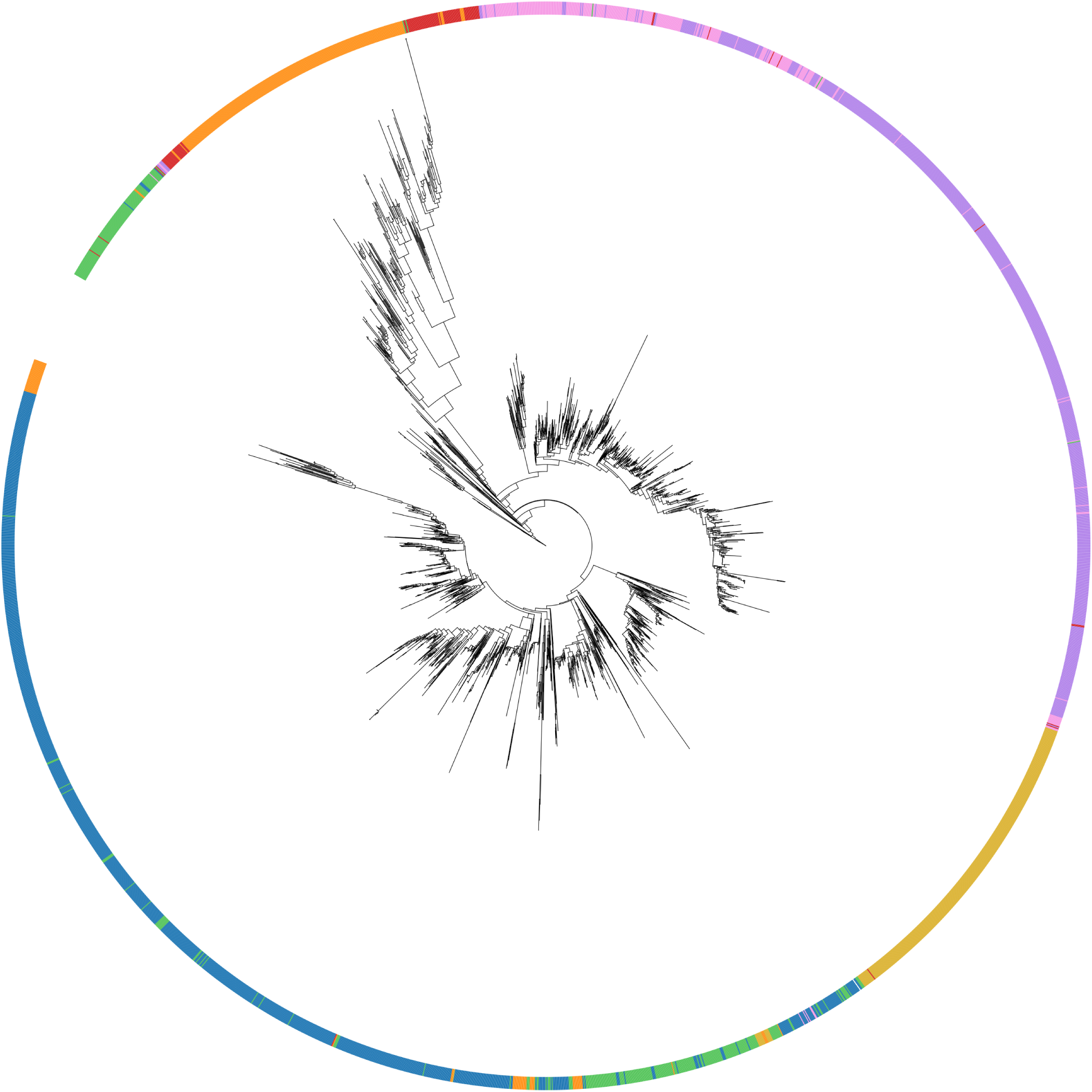
Phylogenetic tree of cold shock domain (PF00313) sequences. The set of 3,287 cold shock sequences considered in the PLMView functional tree in Fig. 6B is organised in a phylogenetic tree. The outer circle colours sequences based on PLMView functional partition, as in the bottom inset legend of Fig. 6B.

**Figure S9:**
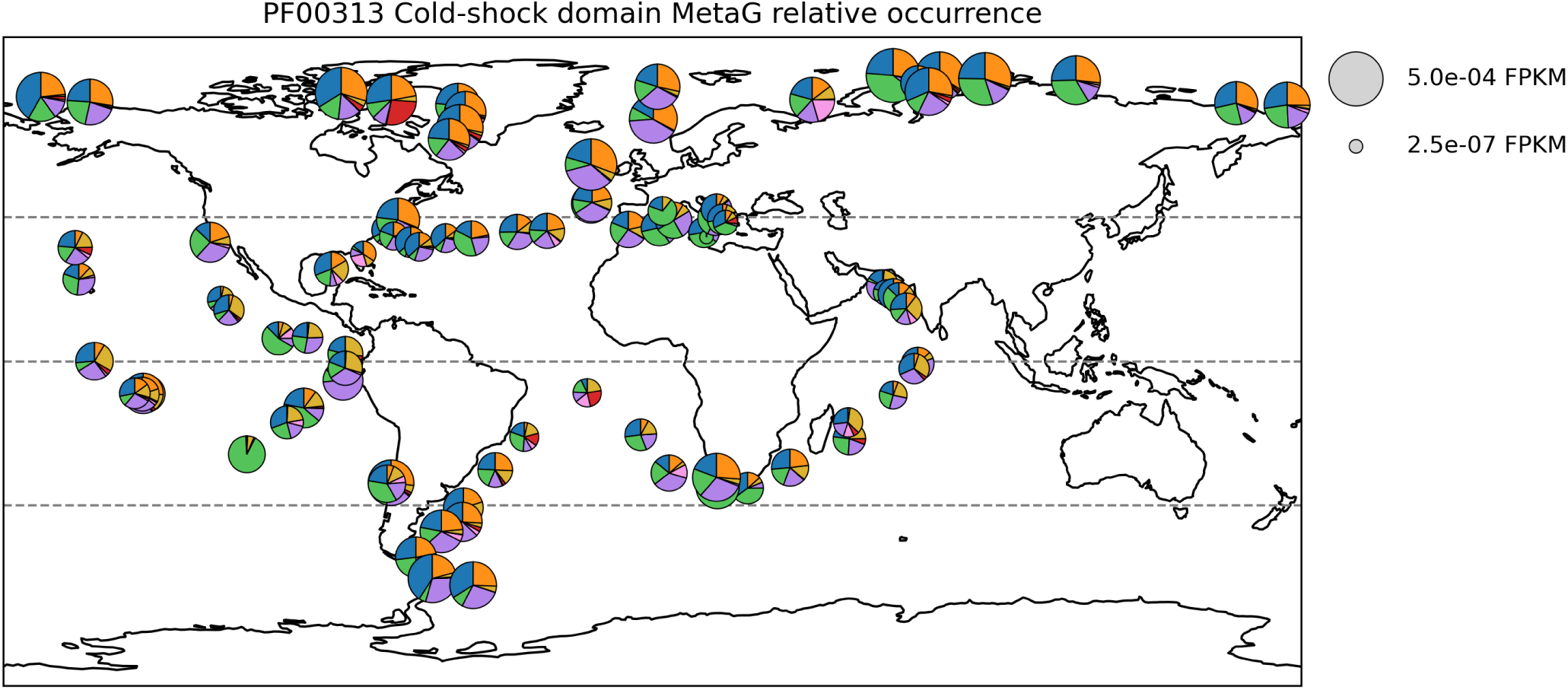
Global distribution of functional metagenomic abundance per sampling station. Colors correspond to the functional partitions defined in Fig. 6B.

**Figure S10:**
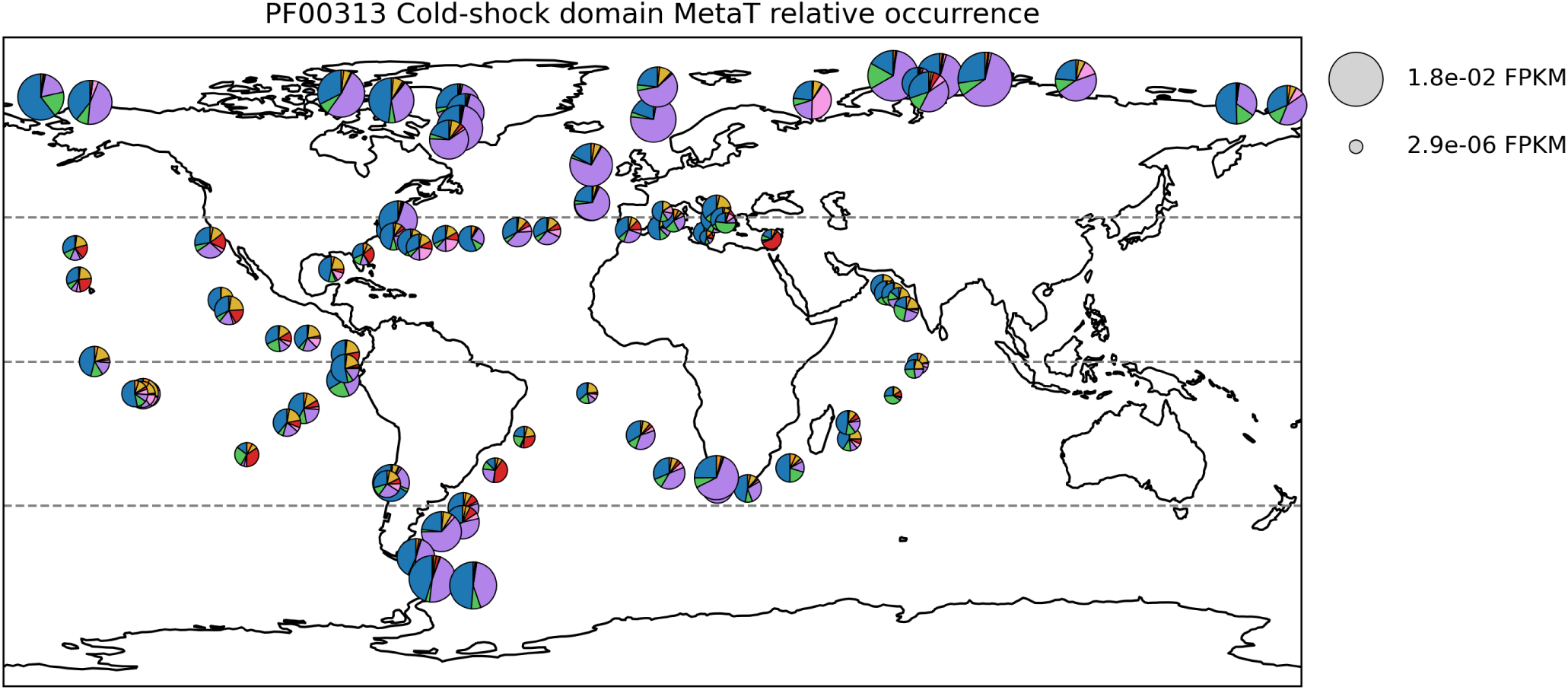
Global distribution of functional metatranscriptomic abundance per sampling station. Colors correspond to the functional partitions defined in Fig. 6B.

**Figure S11:**
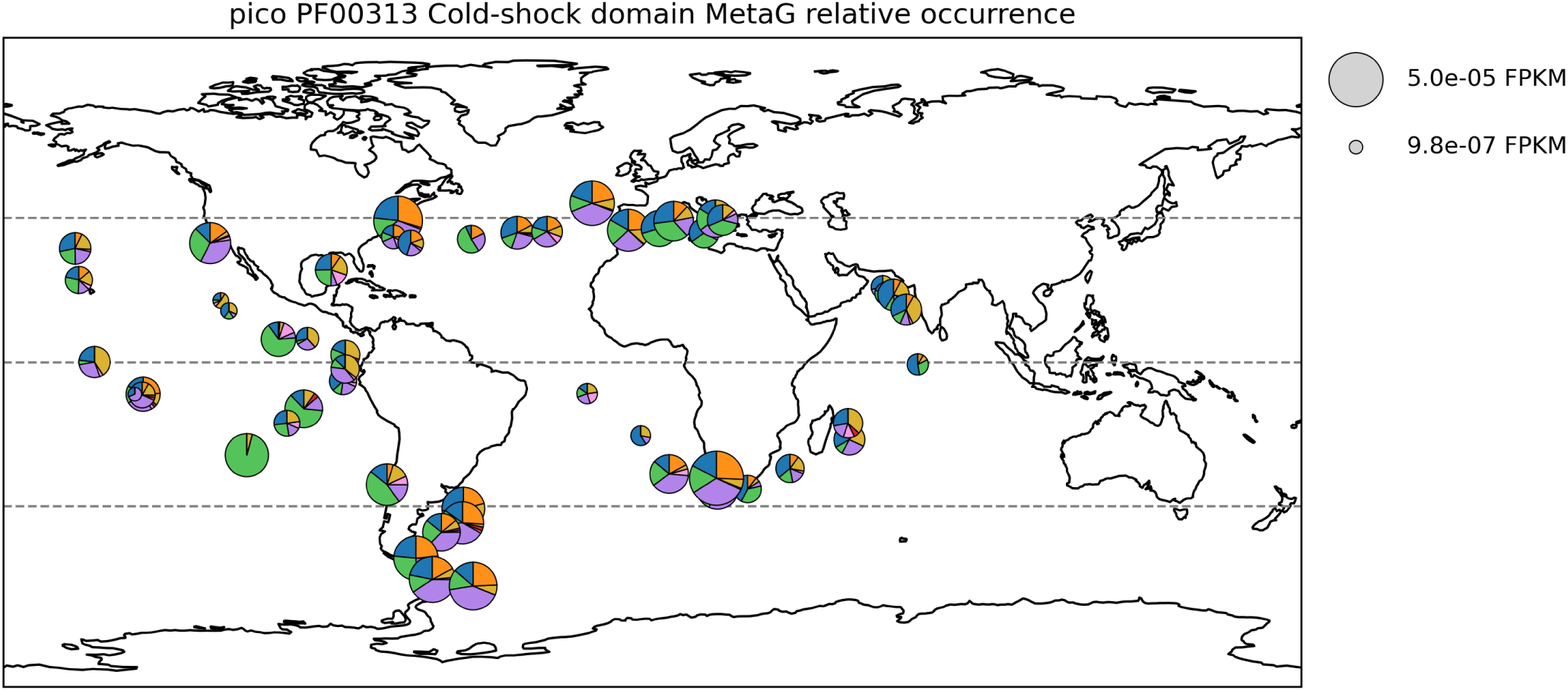
Global distributions of diatom-assigned PF00313 cold-shock domain-encoding sequences in metagenomes from picoplankton size fractions. Colors correspond to the functional partitions defined in Fig. 6B.

**Figure S12:**
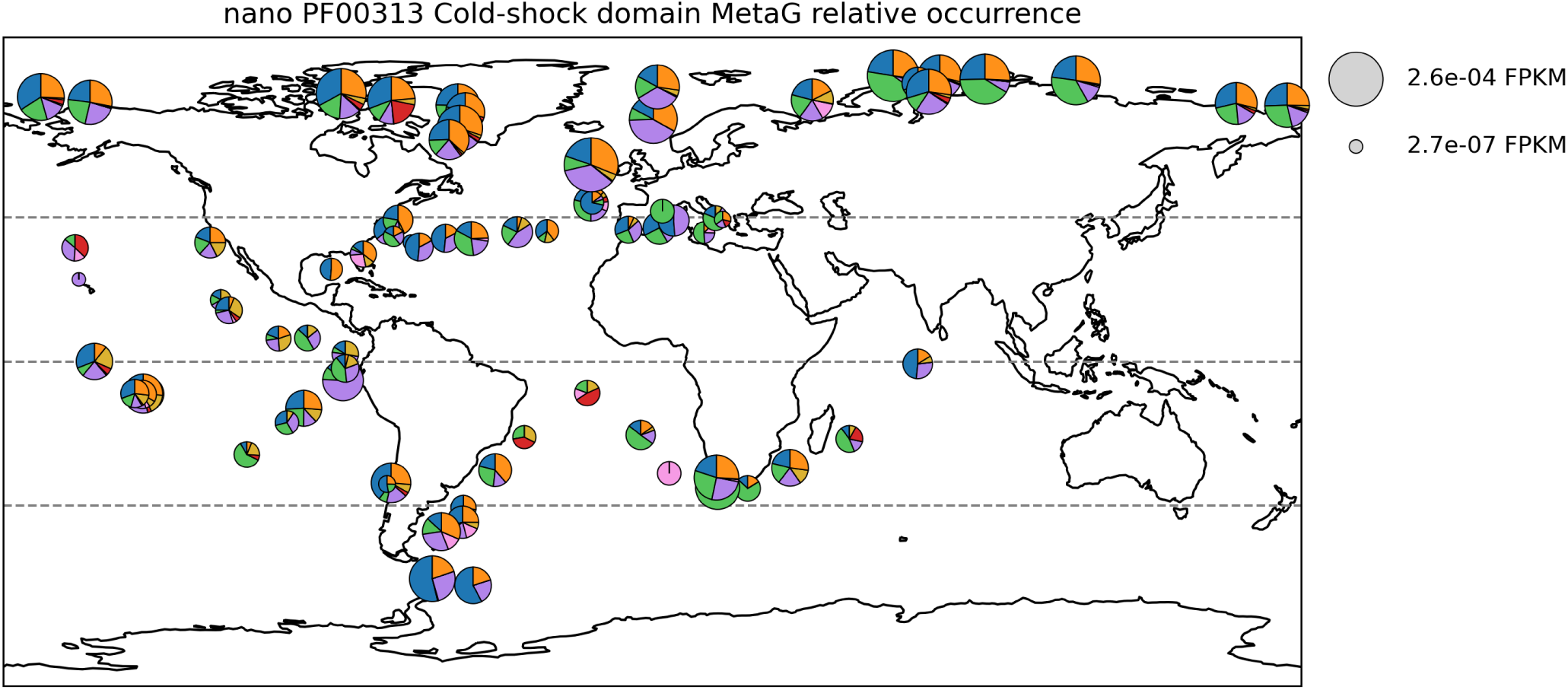
Global distribution of diatom-assigned PF00313 cold-shock domain-encoding sequences in metagenomes from nanoplankton size fractions. Colors correspond to the functional partitions defined in Fig. 6B.

**Figure S13:**
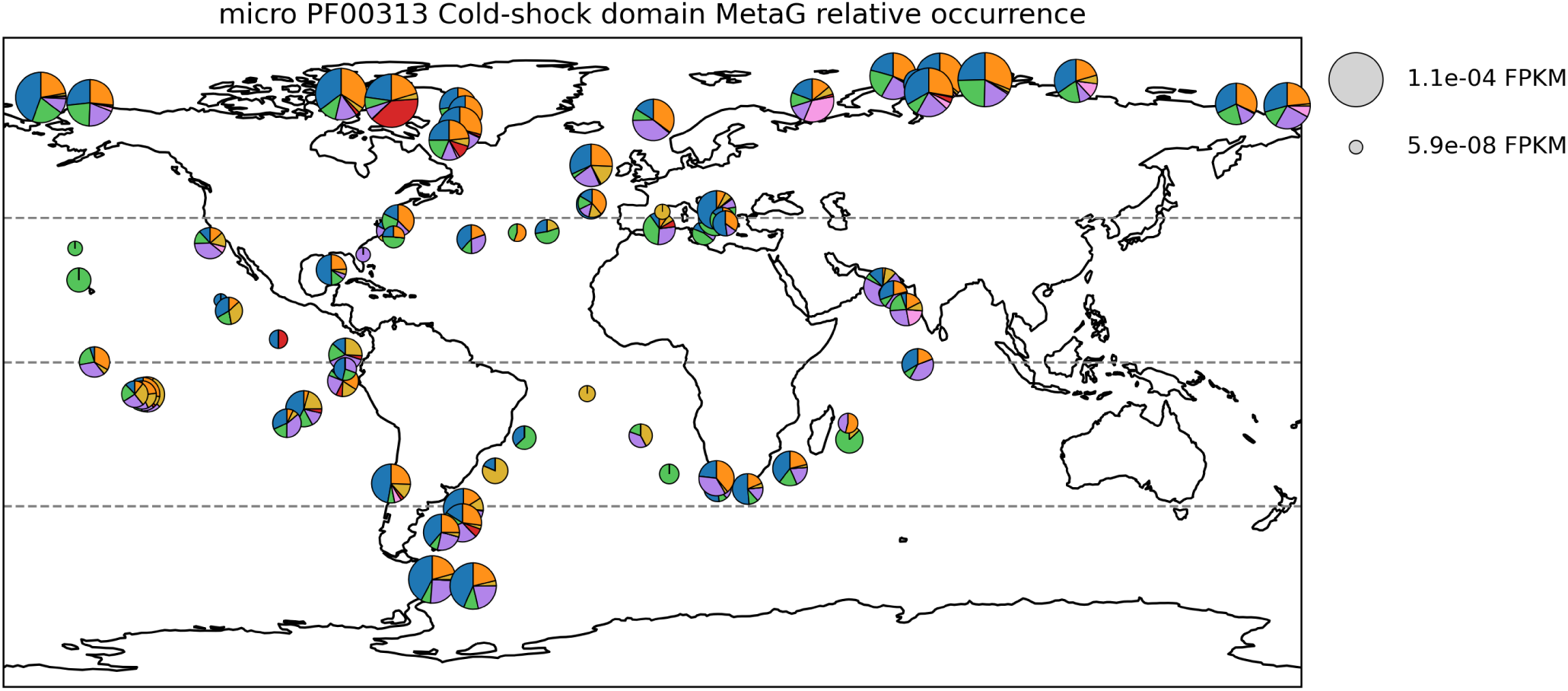
Global distribution of diatom-assigned PF00313 cold-shock domain-encoding sequences in metagenomes from microplankton size fractions. Colors correspond to the functional partitions defined in Fig. 6B.

**Figure S14:**
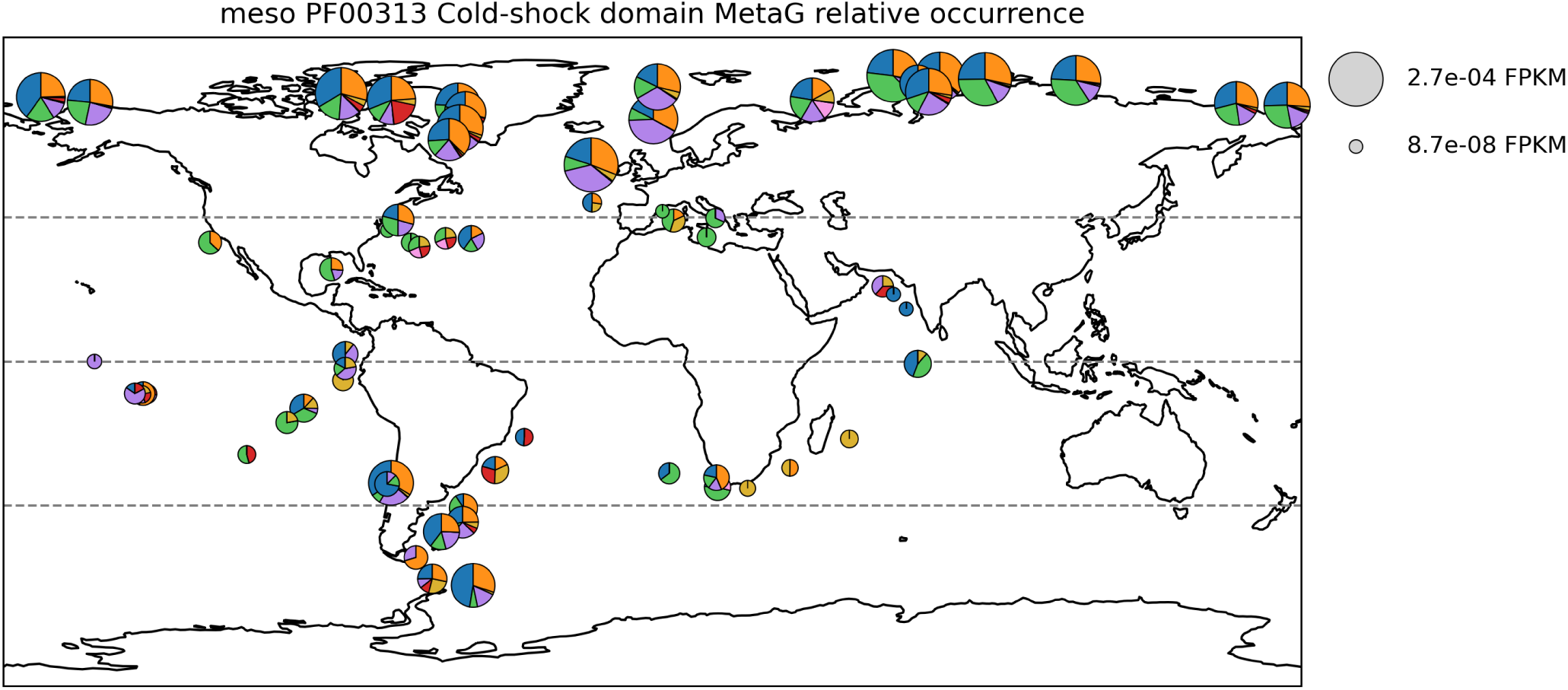
Global distribution of diatom-assigned PF00313 cold-shock domain-encoding sequences in metagenomes from mesoplankton size fractions. Colors correspond to the functional partitions defined in Fig. 6B.

**Figure S15:**
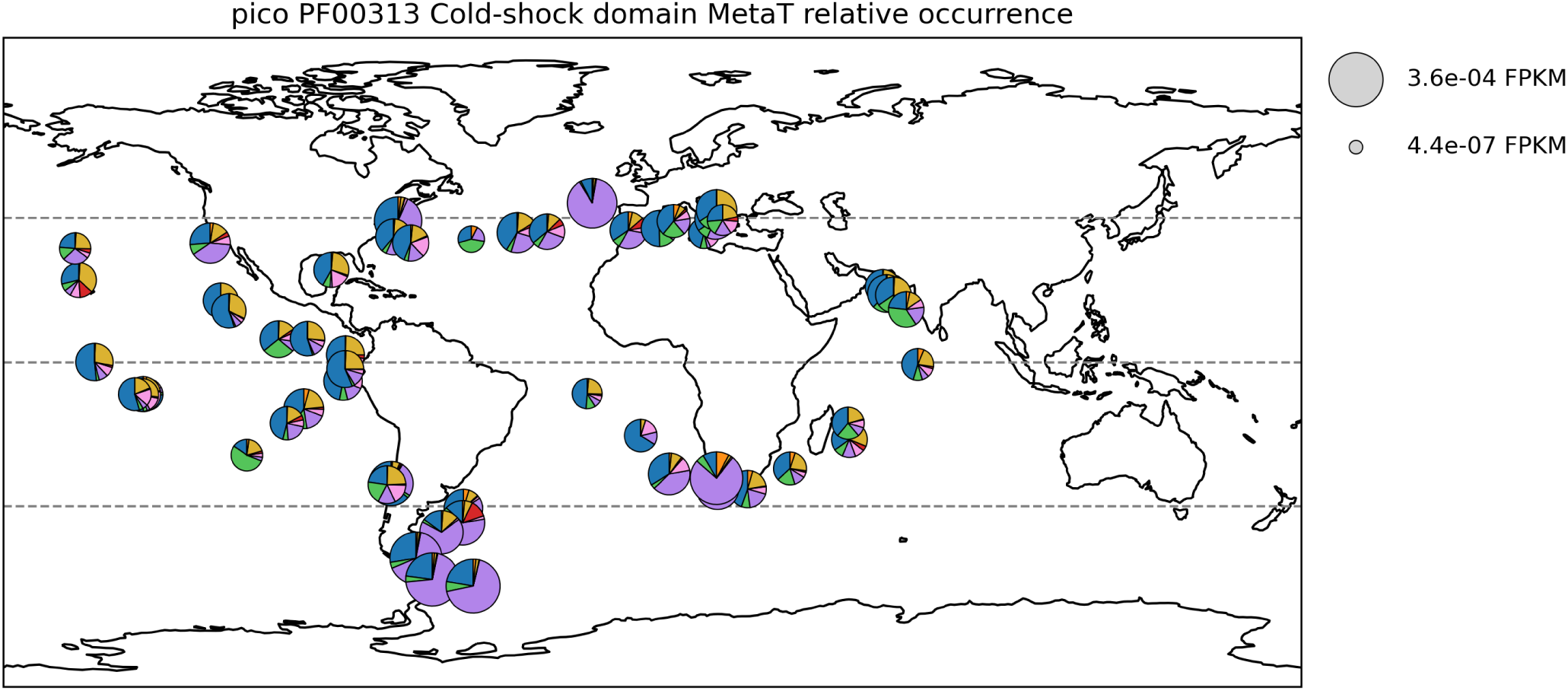
Global distribution of diatom-assigned PF00313 cold-shock domain-encoding sequences in metatranscriptomes from picoplankton size fractions. Colors correspond to the functional partitions defined in Fig. 6B.

**Figure S16:**
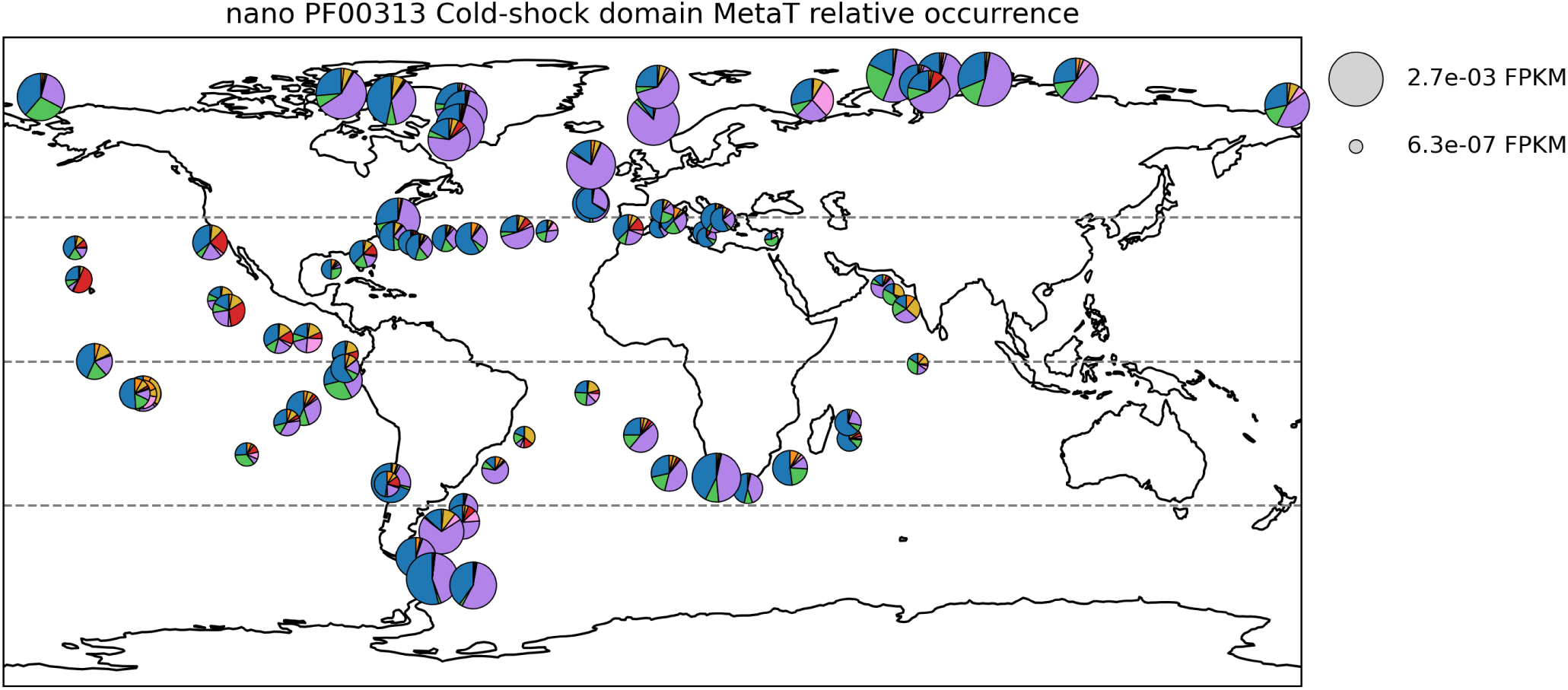
Global distribution of diatom-assigned PF00313 cold-shock domain-encoding sequences in metatranscriptomes from nanoplankton size fractions. Colors correspond to the functional partitions defined in Fig. 6B.

**Figure S17:**
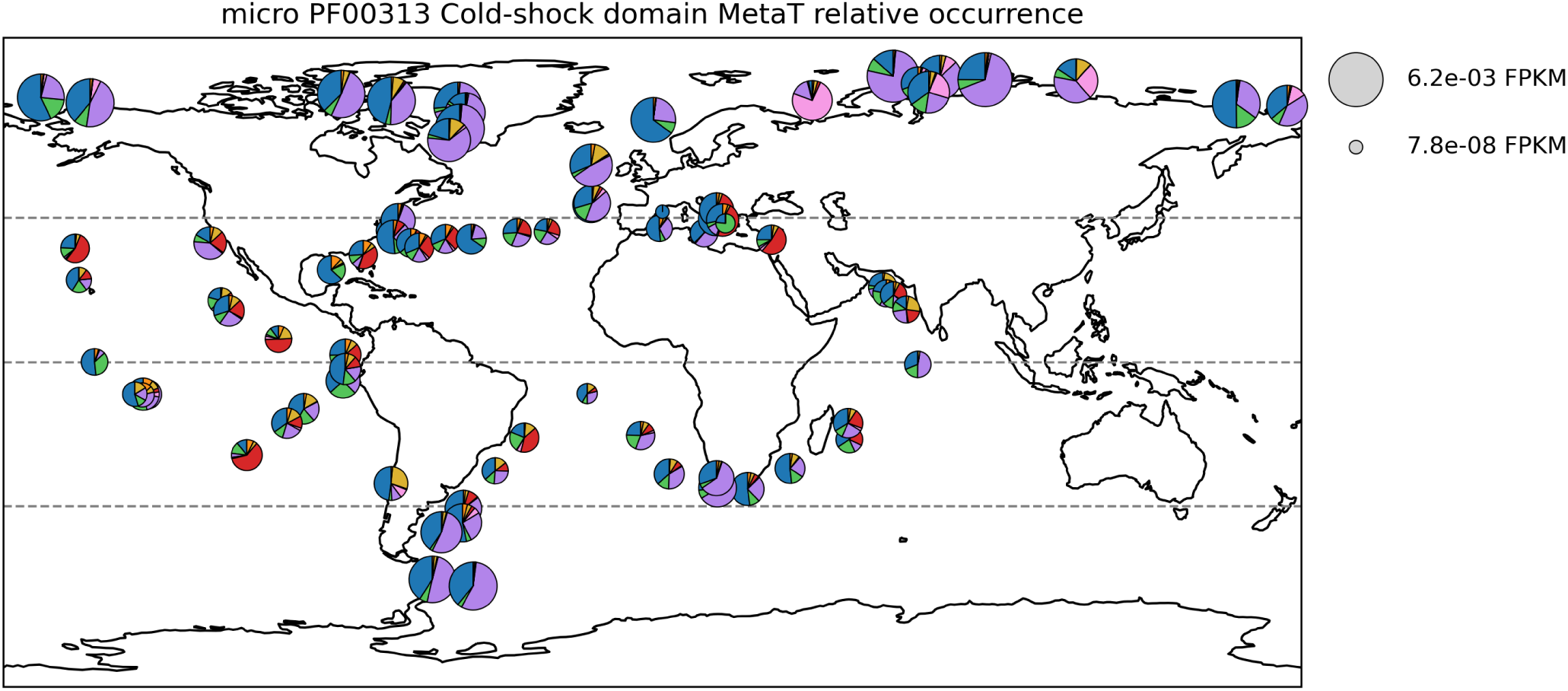
Global distribution of diatom-assigned PF00313 cold-shock domain-encoding sequences in metatranscriptomes from microplankton size fractions. Colors correspond to the functional partitions defined in Fig. 6B.

**Figure S18:**
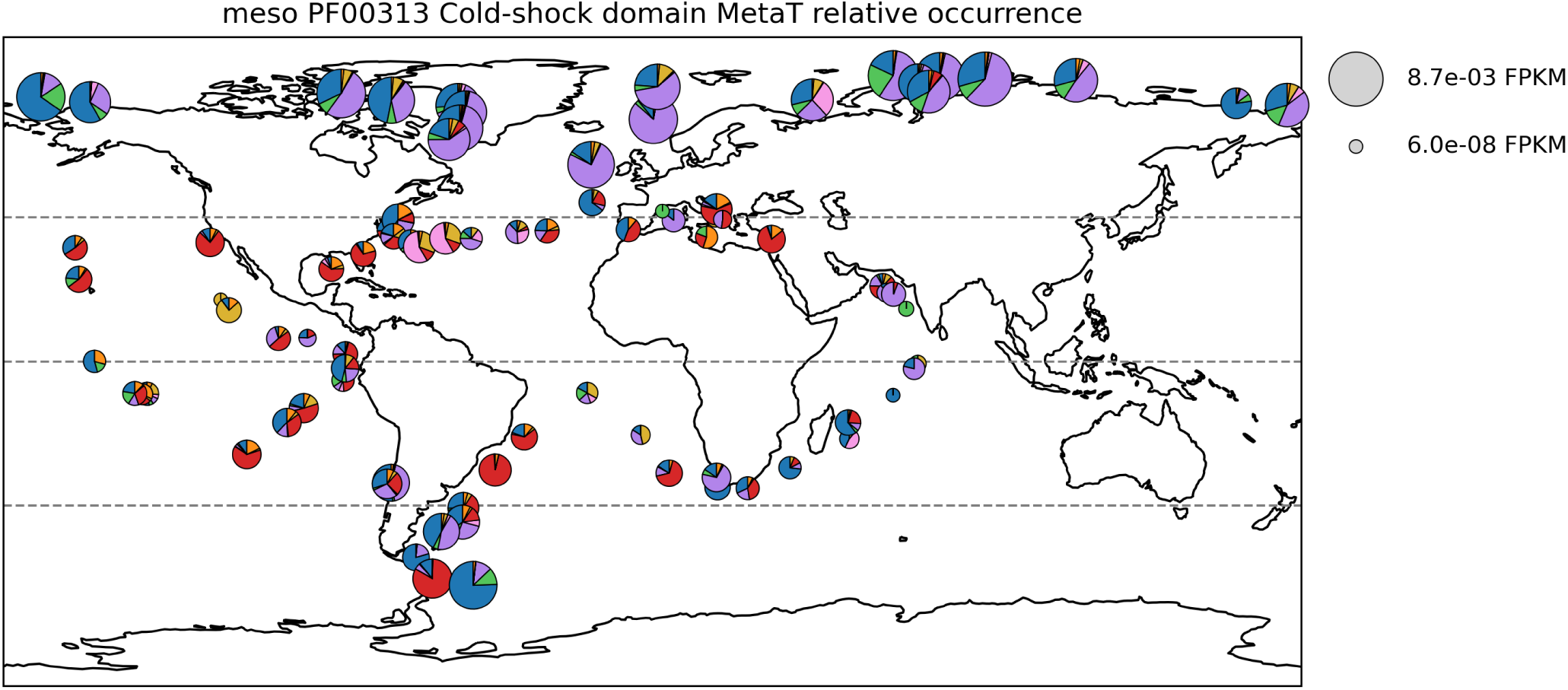
Global distribution of diatom-assigned PF00313 cold-shock domain-encoding sequences in metatranscriptomes from mesoplankton size fractions. Colors correspond to the functional partitions defined in Fig. 6B. This panel corresponds to an enlarged view of the mesoplankton metatranscriptomic map shown in Fig. 6A.

**Figure S19:**
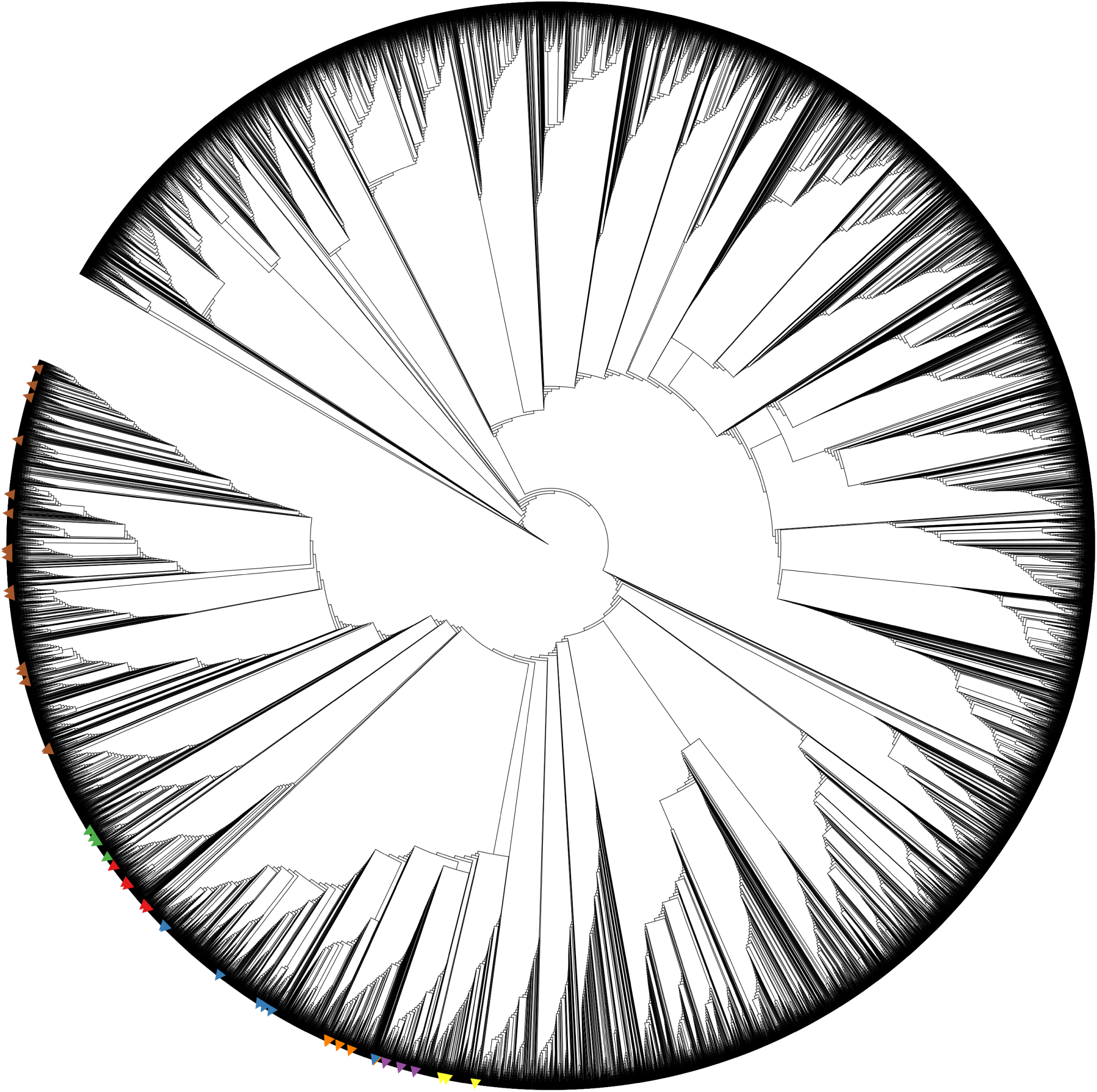
Sequence Similarity Network of trimmed TRX sequences based on NBS distances. Sequences are the same as those used in the PLMView reconstruction shown in Fig. S3A. Annotated sequences are indicated by colored triangles corresponding to known TRX classes. The SSN was constructed from trimmed sequences using NBS distances, and clustering was performed using the average-linkage clustering method. Compare to Fig. S20

**Figure S20:**
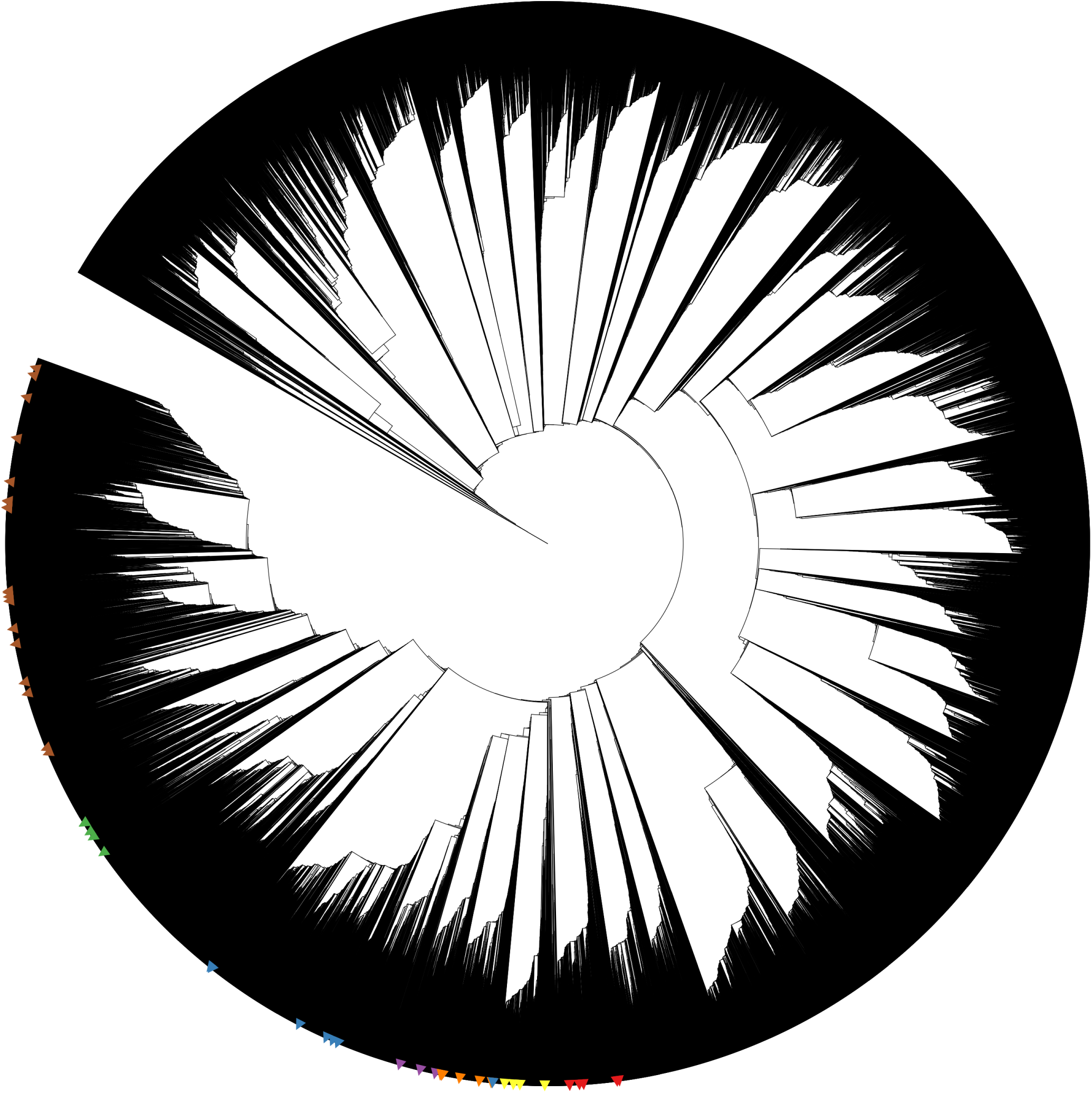
Sequence Similarity Network of trimmed TRX sequences based on e-value distances. Sequences are the same as those used in the PLMView reconstruction shown in Fig. S3A. Annotated sequences are indicated by colored triangles corresponding to known TRX classes. The SSN was constructed from trimmed sequences using e-value distances, and clustering was performed using the single-linkage clustering method. Compare to Fig. S19.

**Figure S21:**
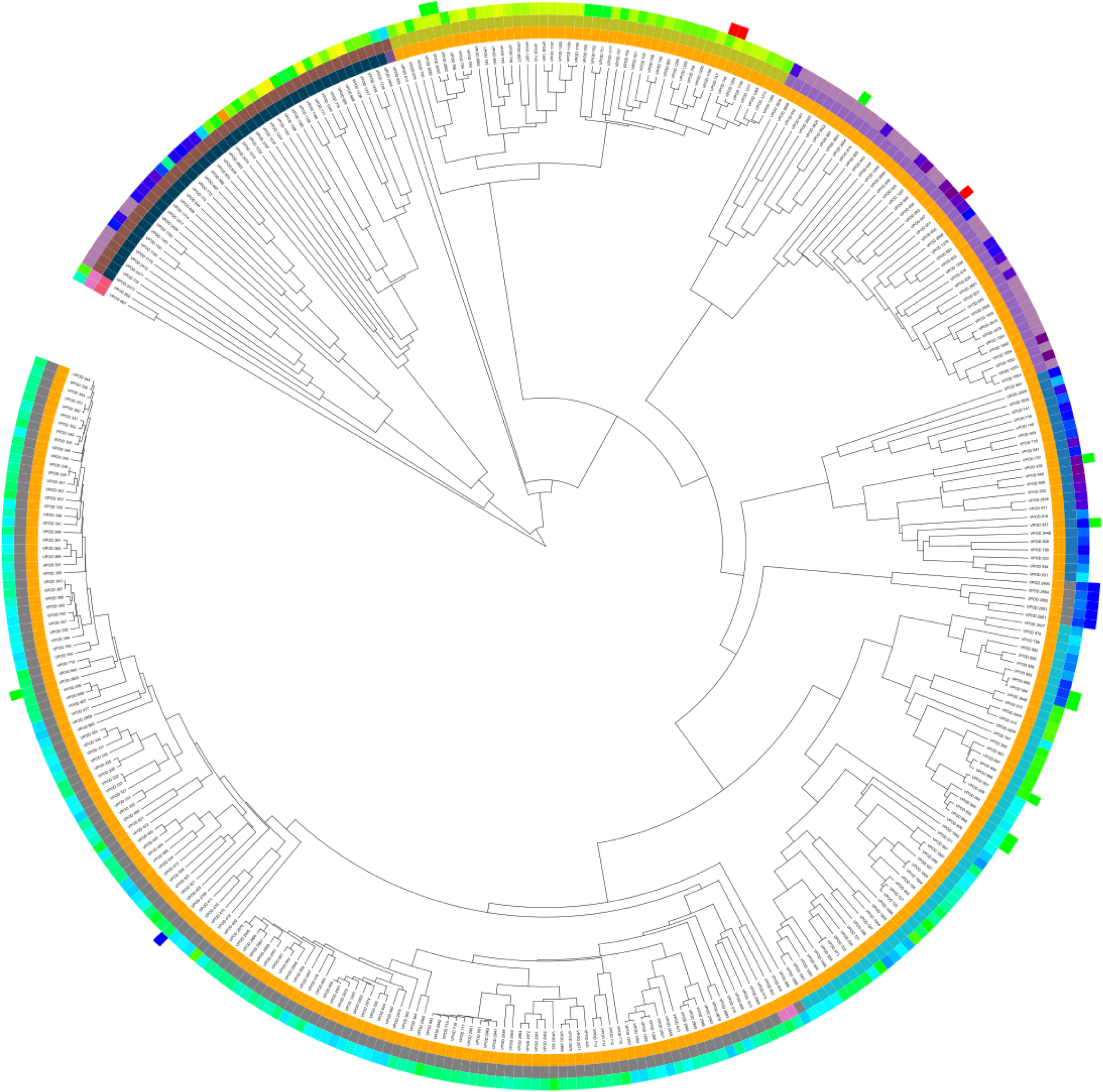
Sequence Similarity Network of Visual Opsin sequences, based on NBS distance. Sequences are the same as in PLMView reconstruction of Fig. 5A. See legend of Fig. 5A for color coding and description of the rings. The analysis is based on NBS distances and the tree has been constructed with the average-linkage clustering method. Compare to Fig. S22.

**Figure S22:**
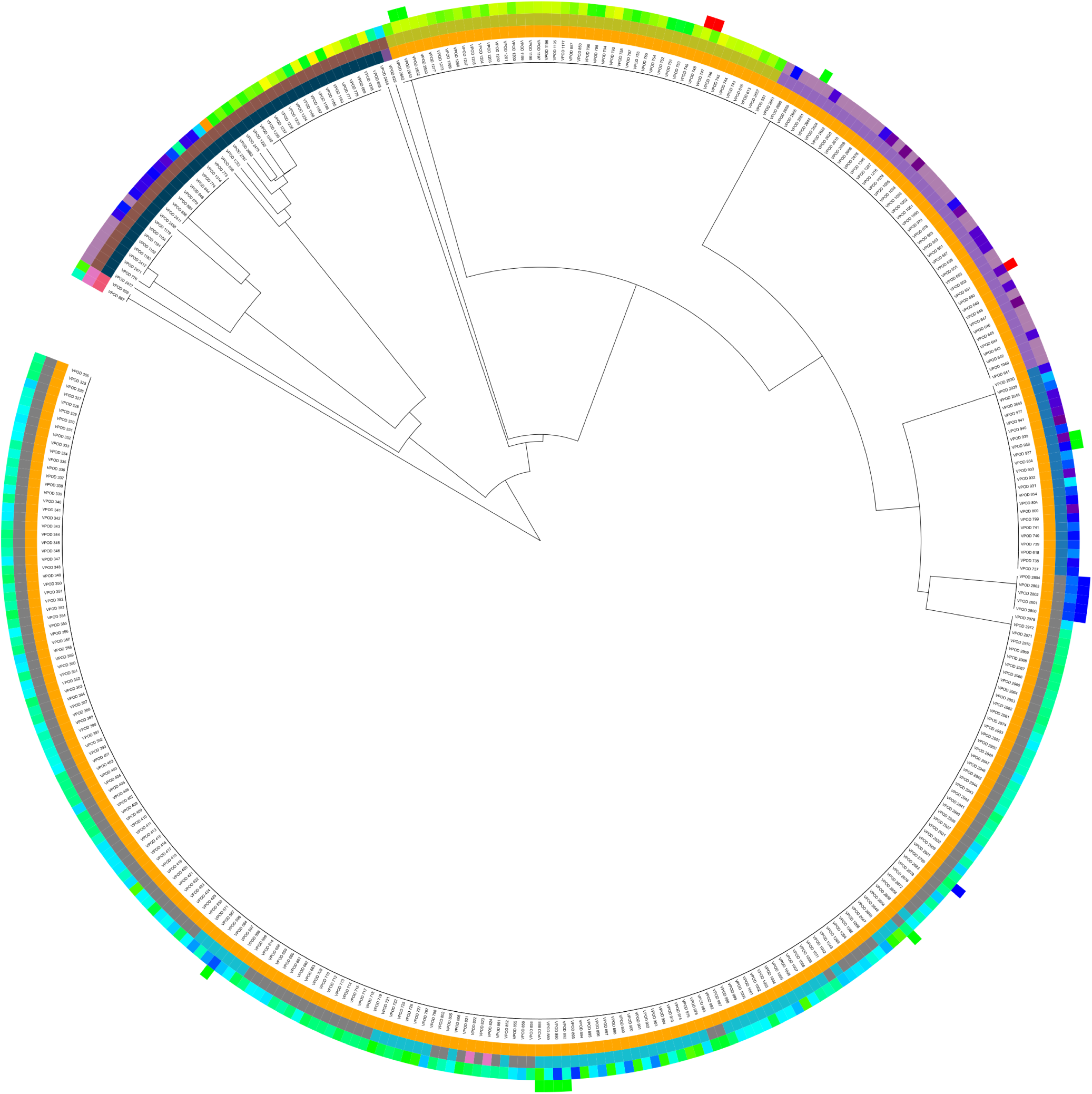
Sequence Similarity Network of Visual Opsin sequences, based on e-value distance. Sequences are the same as in PLMView reconstruction of Fig. 5A. See legend of Fig. 5A for color coding and description of the rings. The analysis is based on e-value distances and the tree has been constructed with the single-linkage clustering method. Compare to Fig. S21.

**Figure S23:**
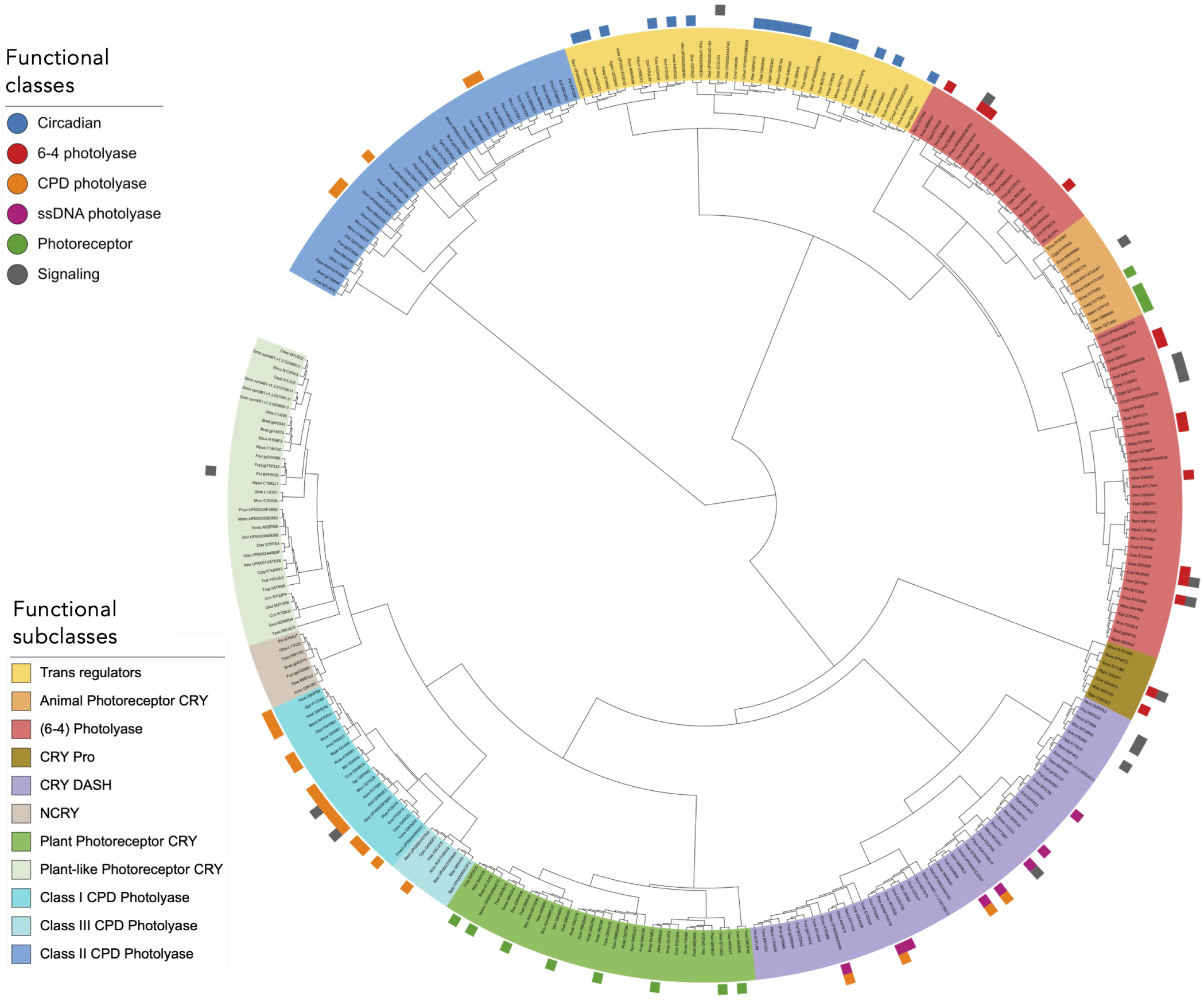
PLMView tree of CPF sequences. Subtrees are colored by considering functionally characterized sequences and by identifying the largest subtree containing exclusively these sequences, following the analysis in ^28^.

**Figure S24:**
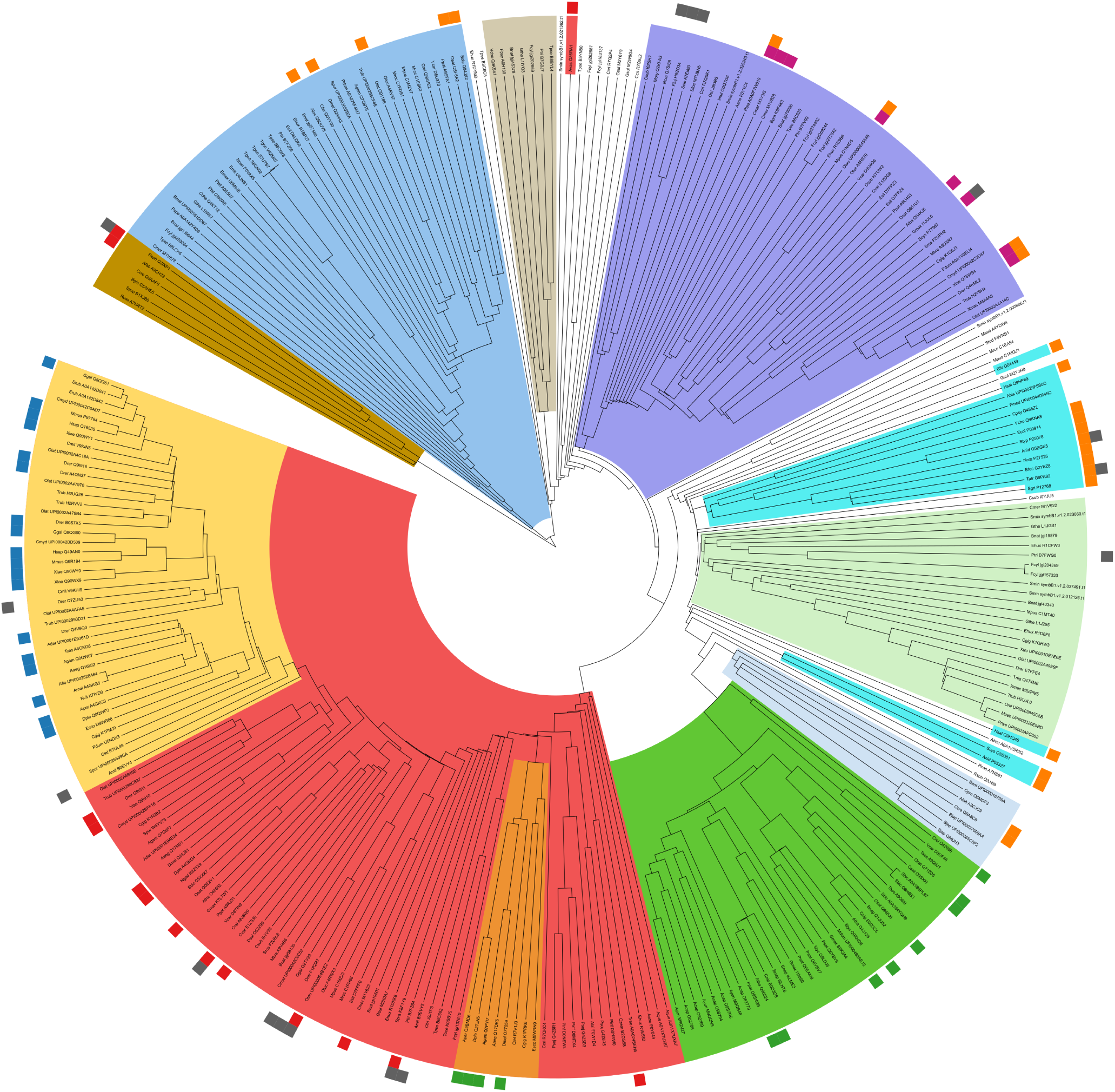
Sequence Similarity Network of CPF sequences based on NBS distances. Sequences are the same as in the PLMView reconstruction shown in Fig. S23. Functionally characterized sequences are highlighted in the outer circle. Subtrees are color-coded according to the legend in Fig. S23, using the clade coloring feature in iTOL. The tree was constructed using the NBS distance and the average-linkage clustering method (see Methods).

**Figure S25:**
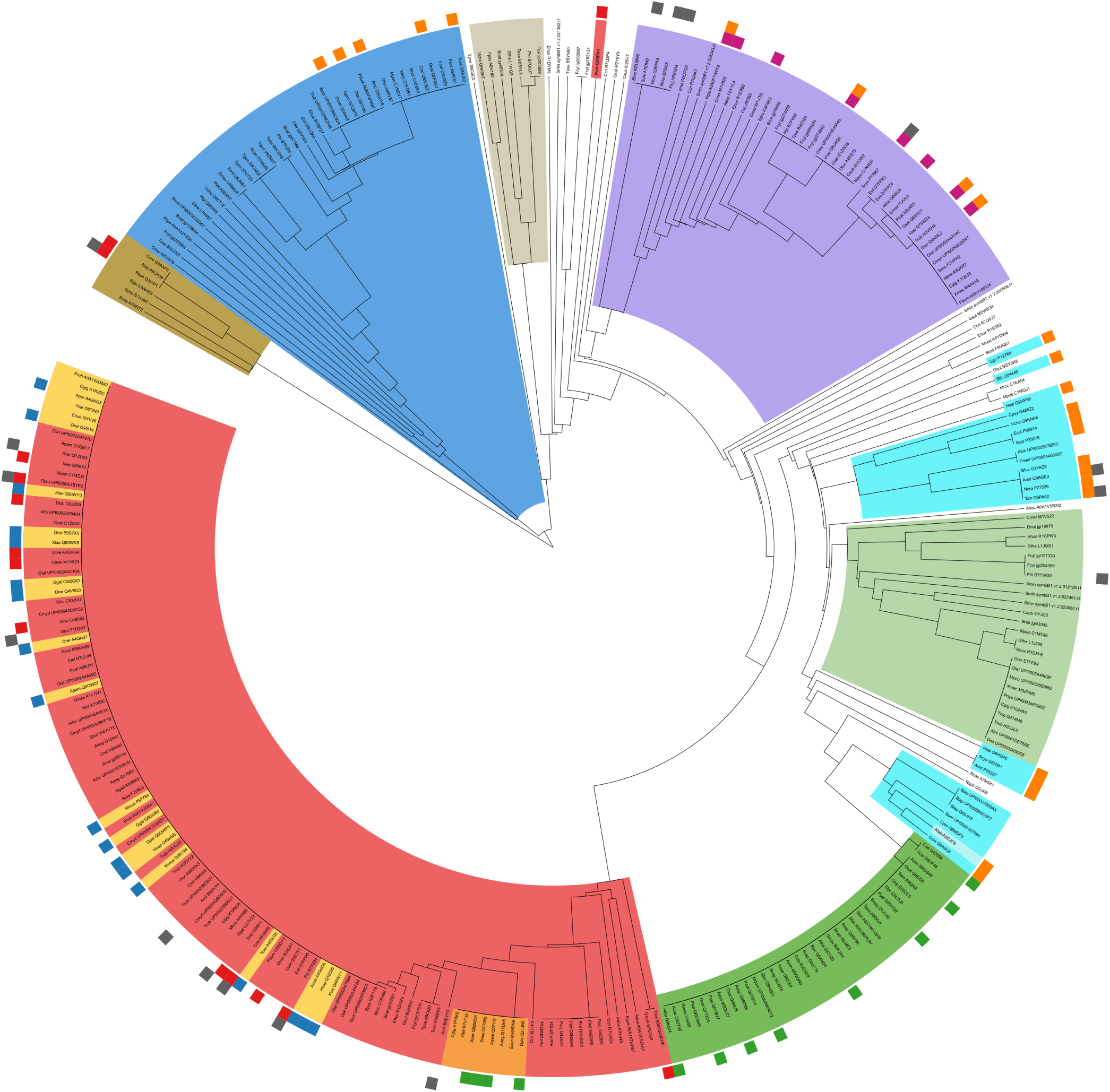
Sequence Similarity Network of CPF sequences based on e-value distances. Sequences are the same as in the PLMView reconstruction shown in Fig. S23. Functionally characterized sequences are highlighted in the outer circle. Subtrees are color-coded according to the legend in Fig. S23, using the clade coloring feature in iTOL. The tree was constructed using the e-value distance and the single-linkage clustering method (see Methods).

**Figure S26:**
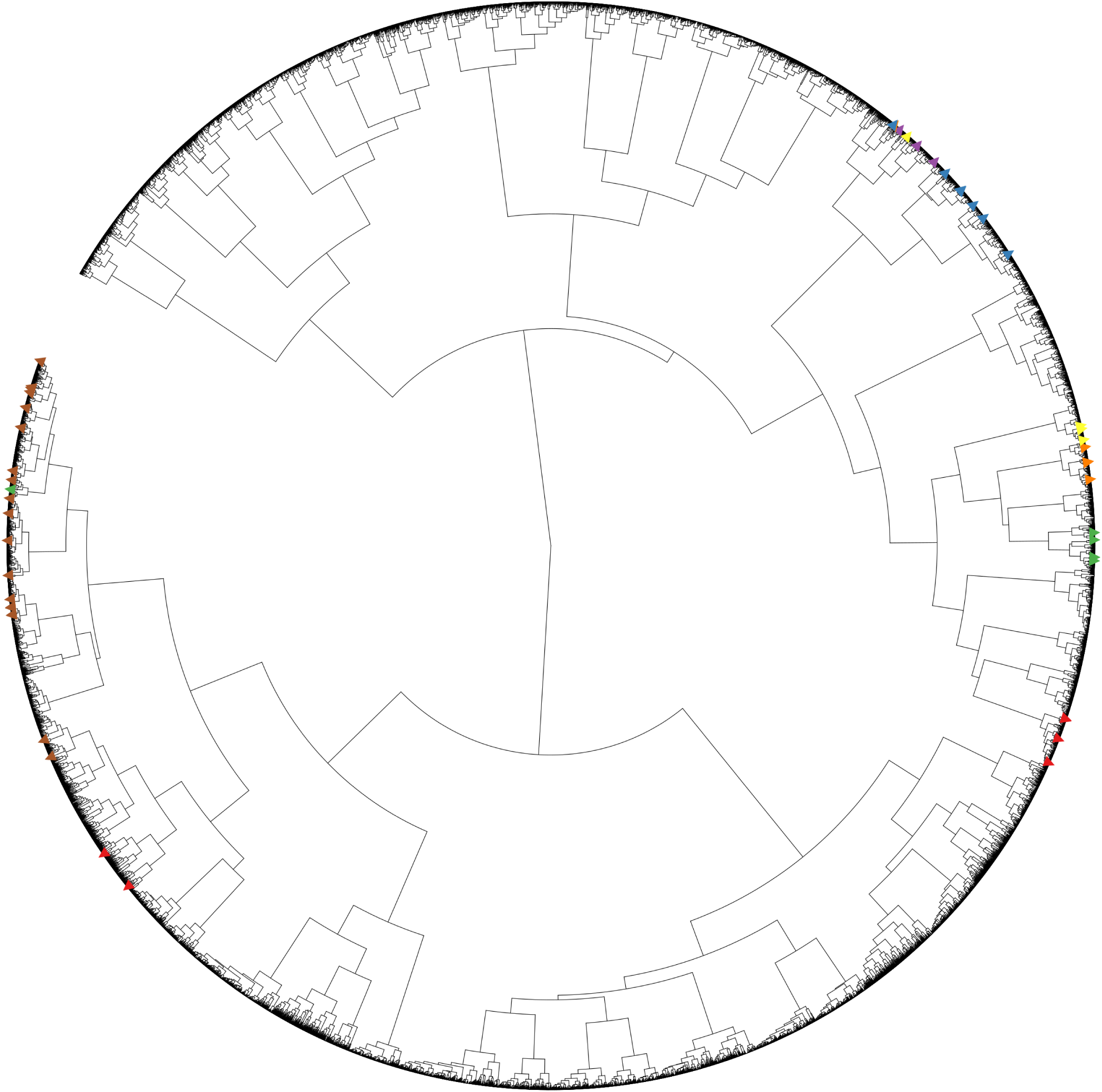
Classification of TRX sequences based on pooled ESM embeddings. TRX sequences have been trimmed to eliminate the transit peptide region. Colors are as in Fig. S3. The tree was constructed using Ward linkage.

**Figure S27:**
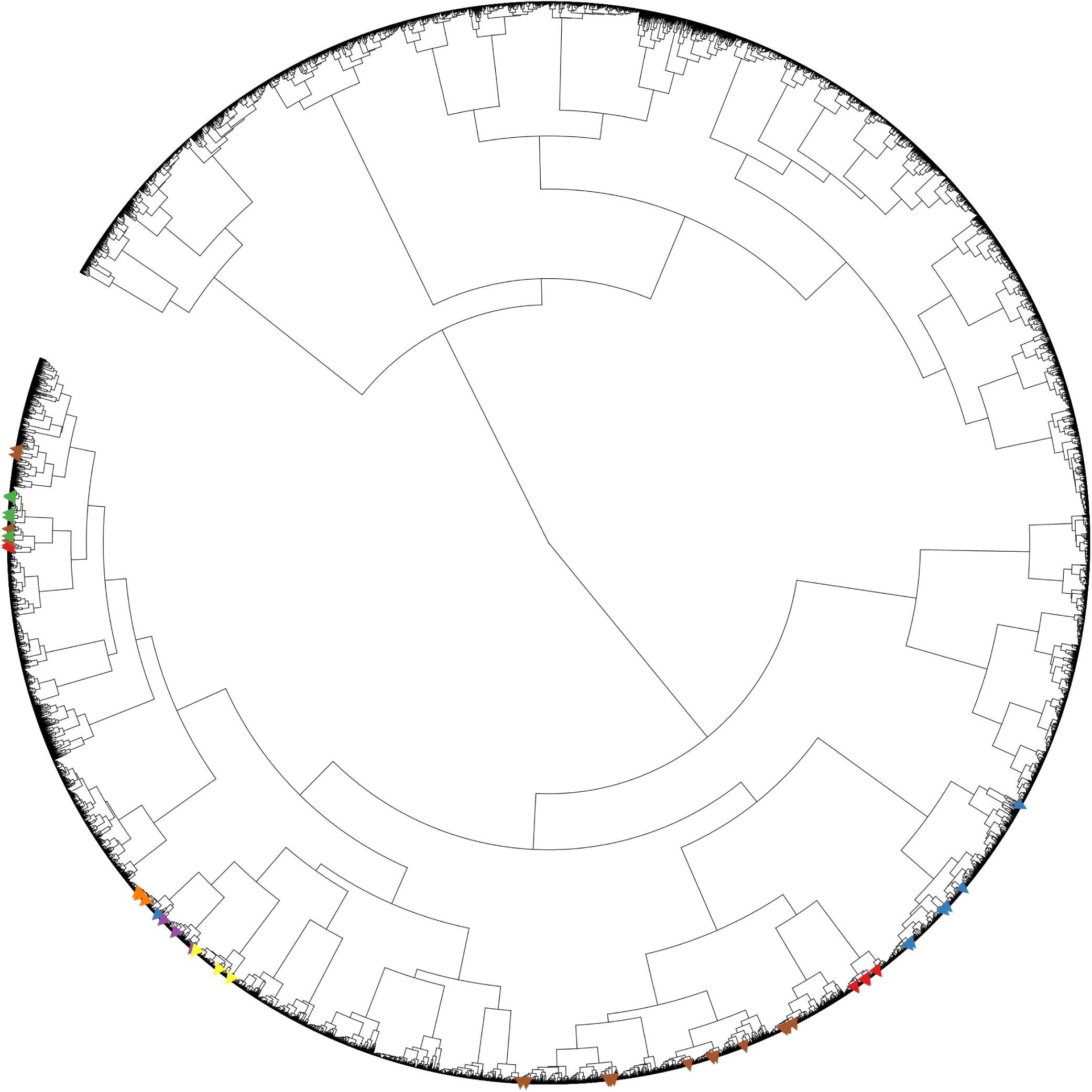
Classification of TRX sequences based on pooled Ankh embeddings. TRX sequences have been trimmed to eliminate the transit peptide region. Colors are as in Fig. S3. The tree was constructed using Ward linkage.

**Figure S28:**
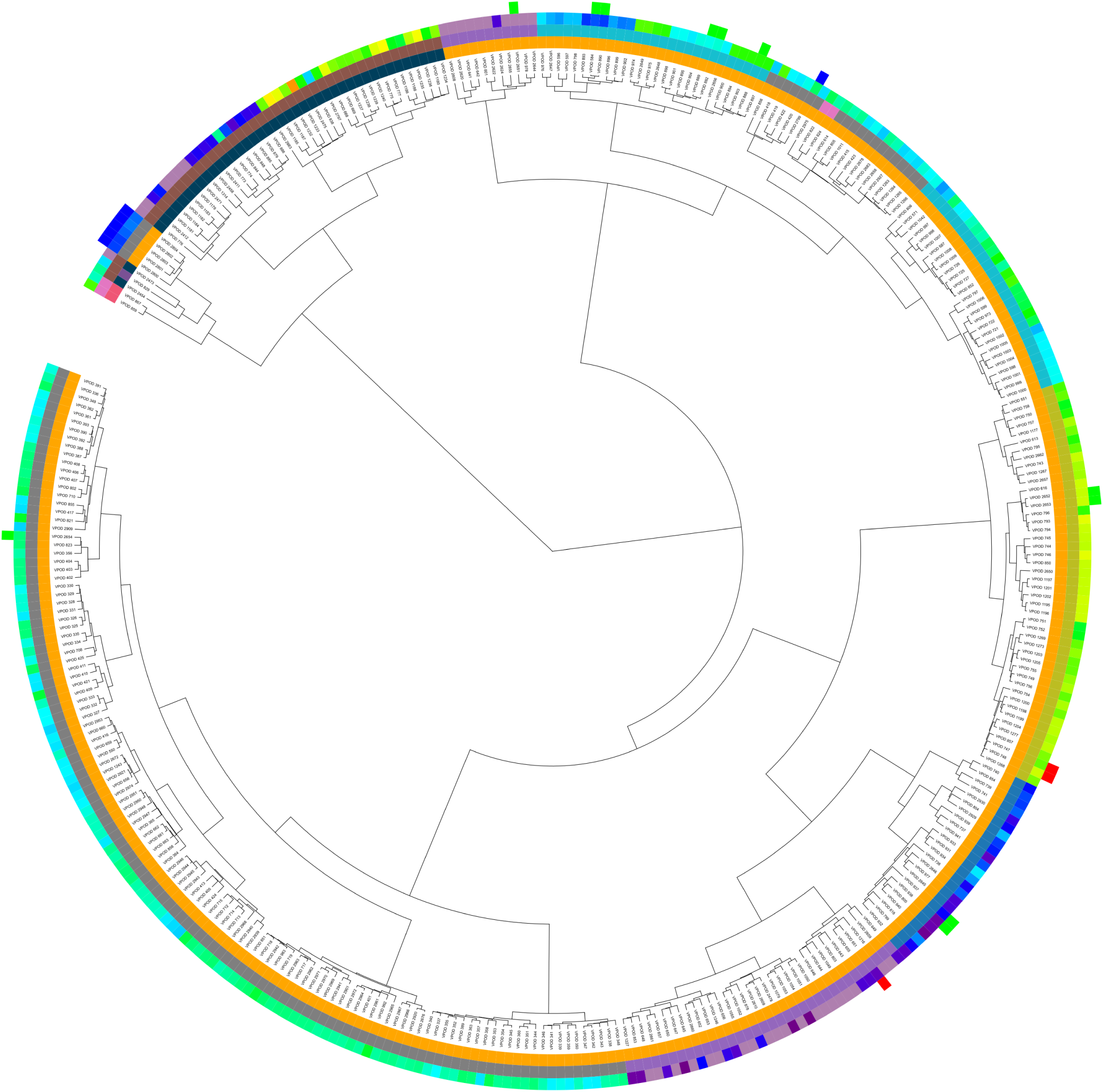
Classification of opsin sequences based on pooled ESM embeddings. Sequences are the same as in PLMView reconstruction of Fig. 5A. See legend of Fig. 5A for color coding and description of the rings. The tree was constructed using Ward linkage.

**Figure S29:**
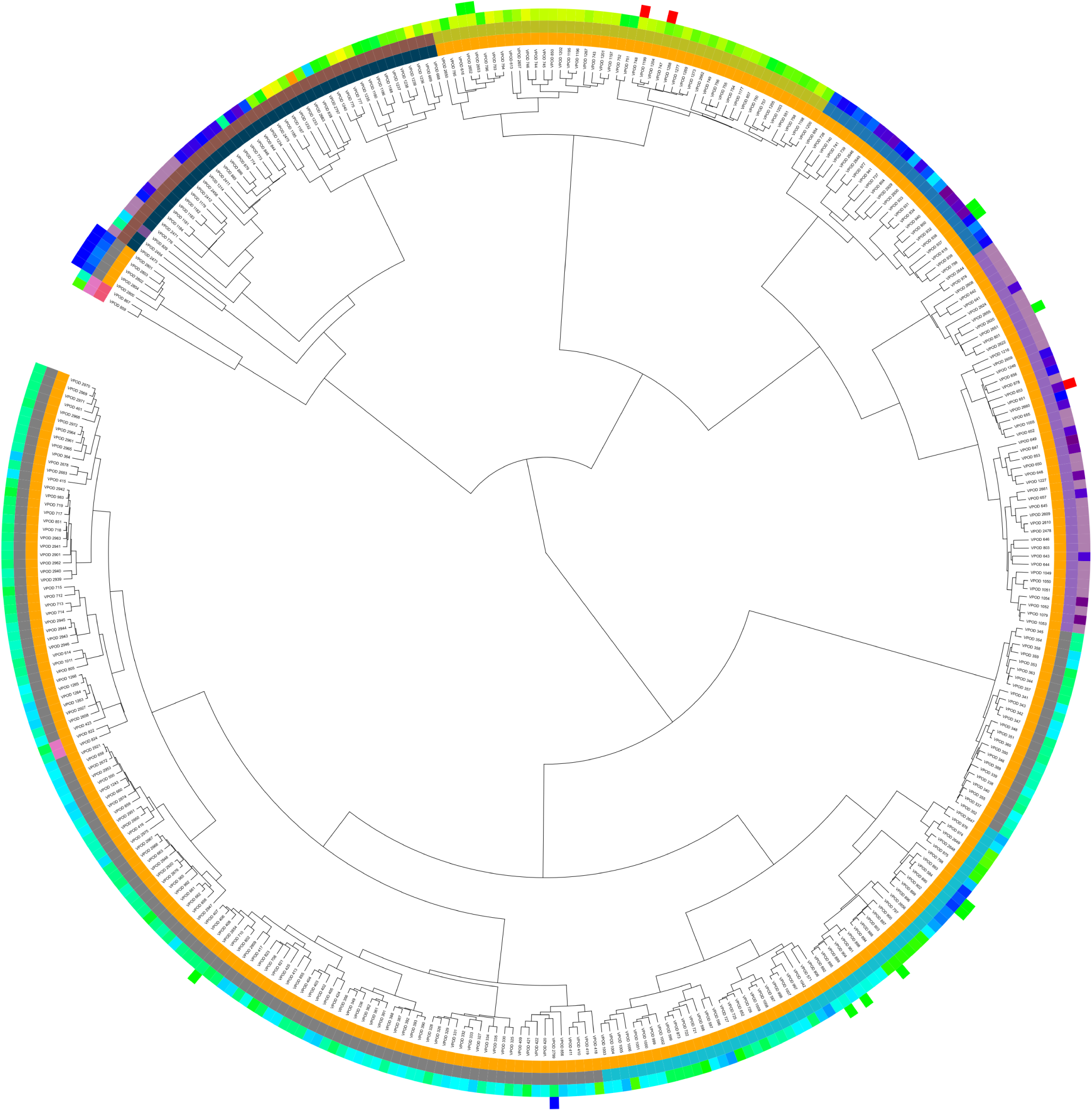
Classification of opsin sequences based on pooled Ankh embeddings. Sequences are the same as in PLMView reconstruction of Fig. 5A. See legend of Fig. 5A for color coding and description of the rings. The tree was constructed using Ward linkage.

**Figure S30:**
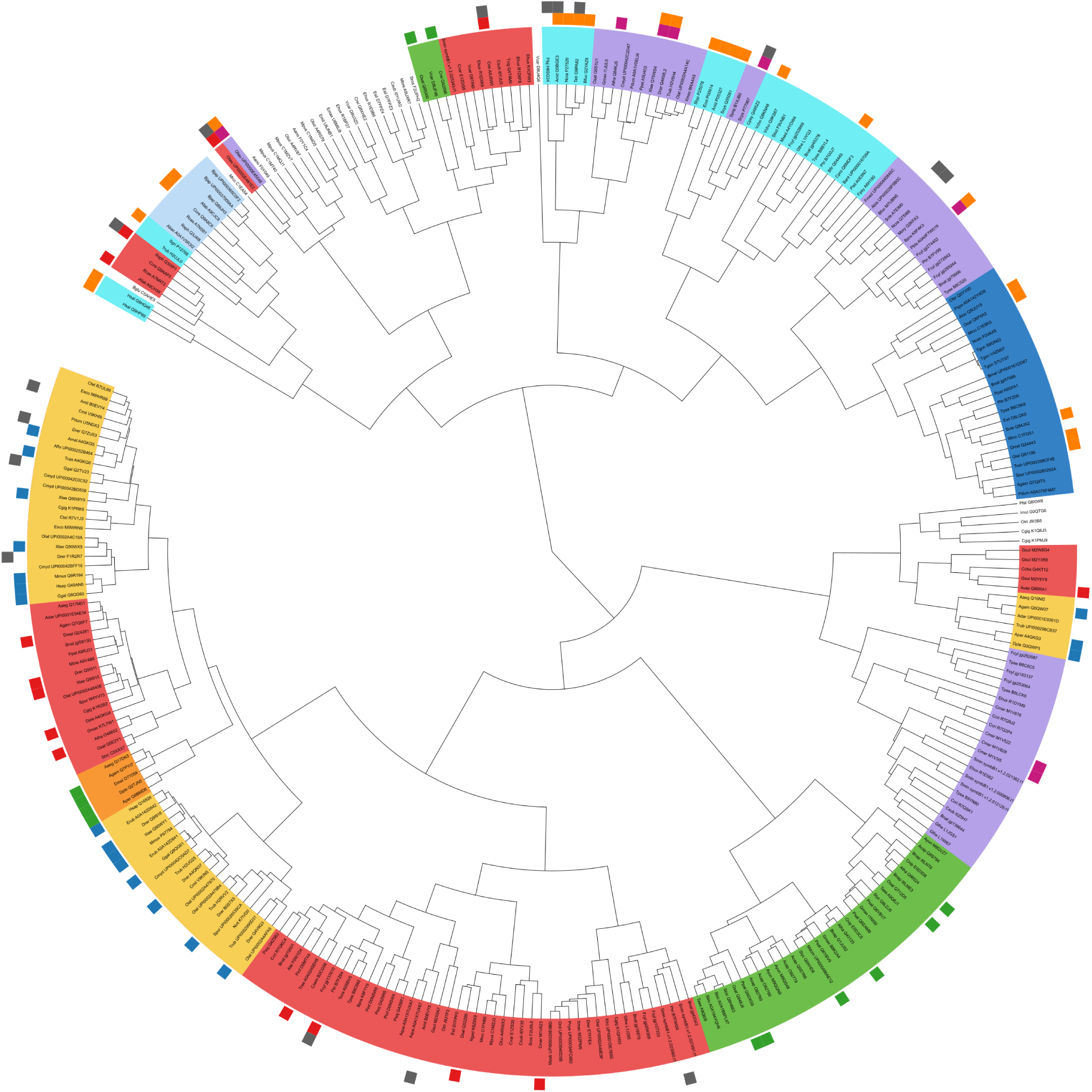
Classification of CPF sequences based on pooled ESM embeddings. Sequences are the same as in the PLMView reconstruction shown in Fig. S23. Functionally characterized sequences are highlighted in the outer circle. Subtrees are color-coded according to the legend in Fig. S23, using the label coloring feature in iTOL. The tree was constructed using Ward linkage.

**Figure S31:**
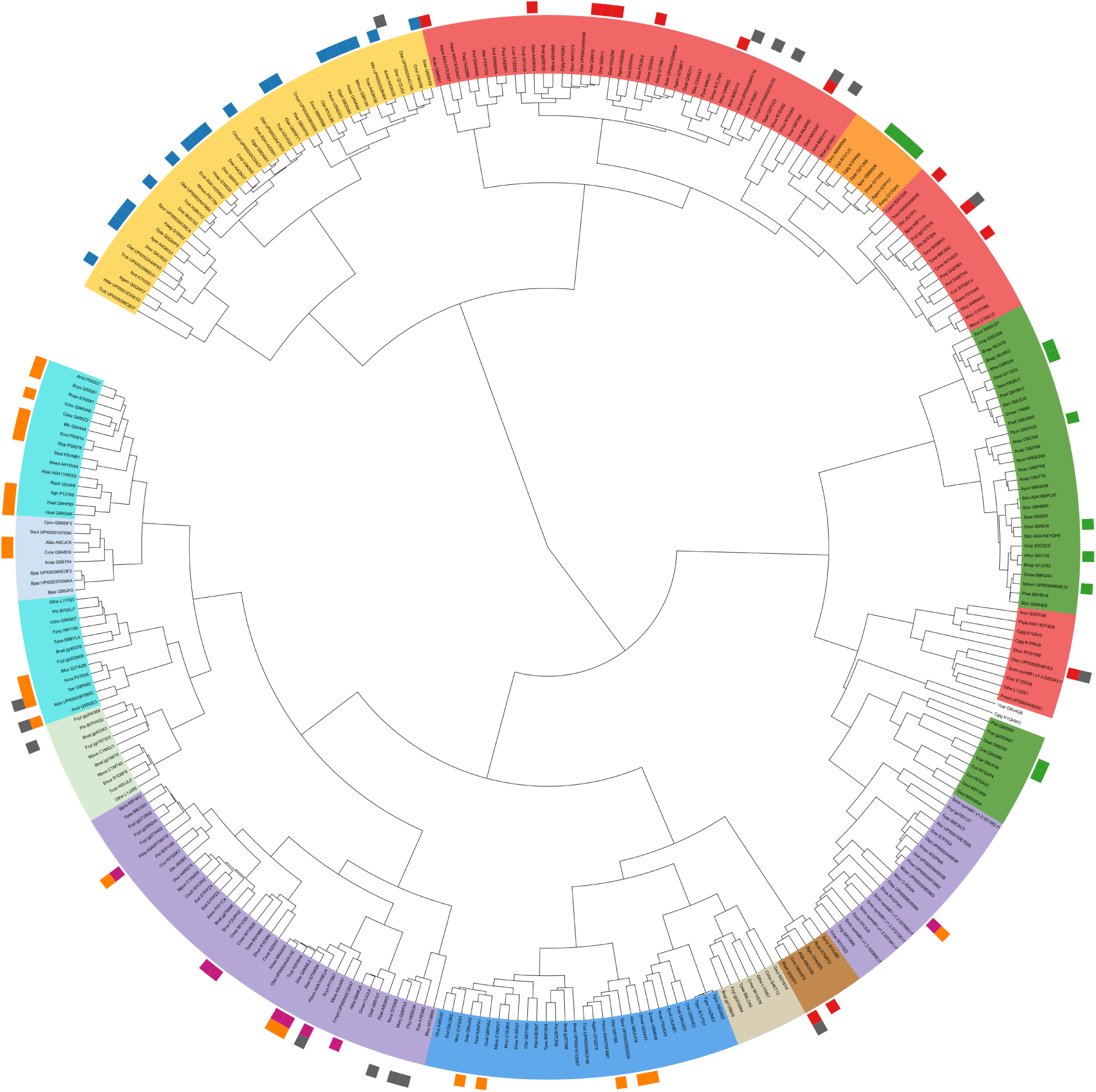
Classification of CPF sequences based on pooled Ankh embeddings. Sequences are the same as in the PLMView reconstruction shown in Fig. S23. Functionally characterized sequences are highlighted in the outer circle. Subtrees are color-coded according to the legend in Fig. S23, using the label coloring feature in iTOL. The tree was constructed using Ward linkage.

**Figure S32:**
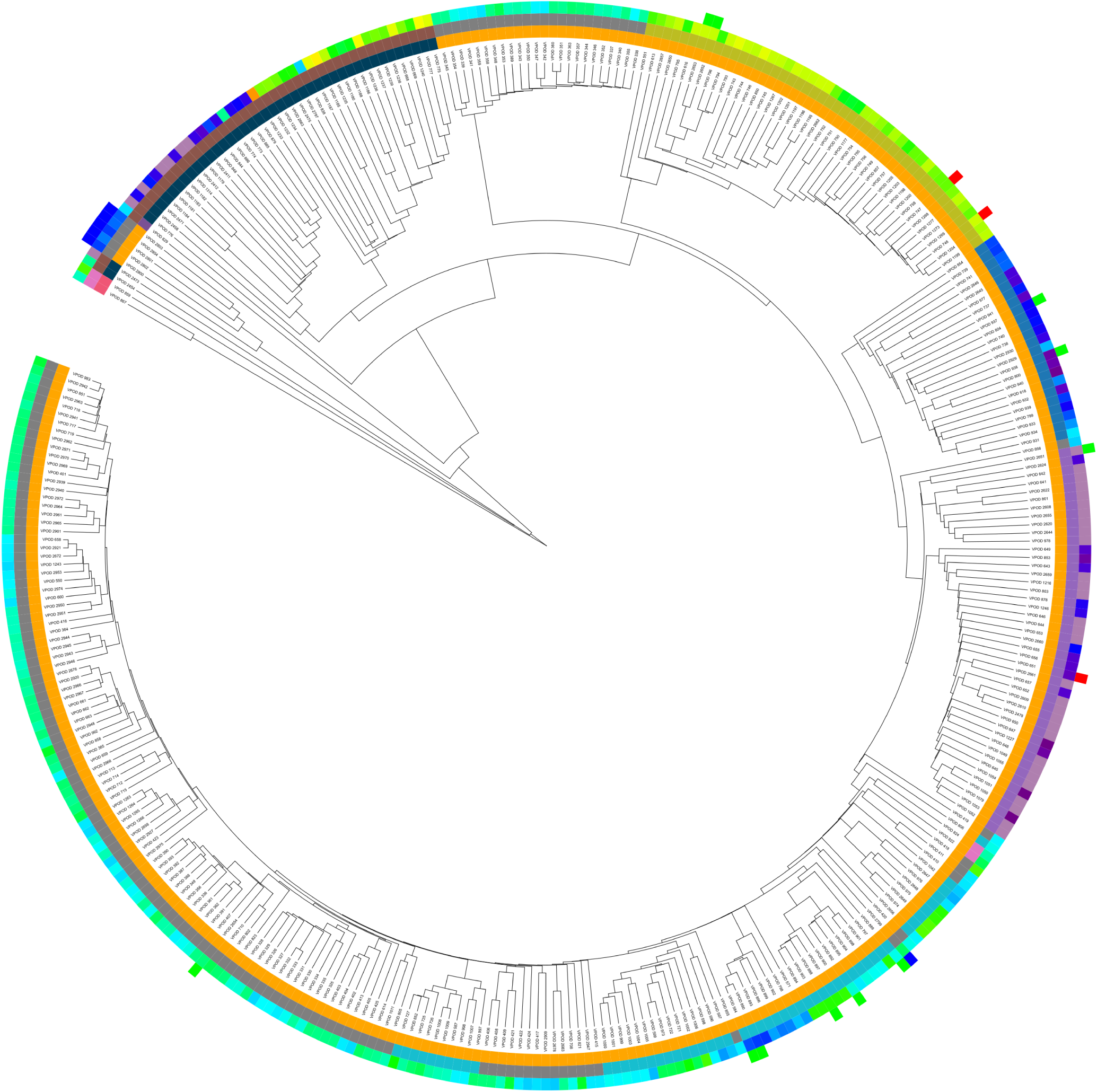
Classification of opsin sequences based on FANTASIA V1/ProtT5 metrics. Sequences are the same as in PLMView reconstruction of Fig. 5A. See legend of Fig. 5A for color coding and description of the rings. The tree was constructed using the euclidean distance and the single-linkage clustering method (see Methods).

**Figure S33:**
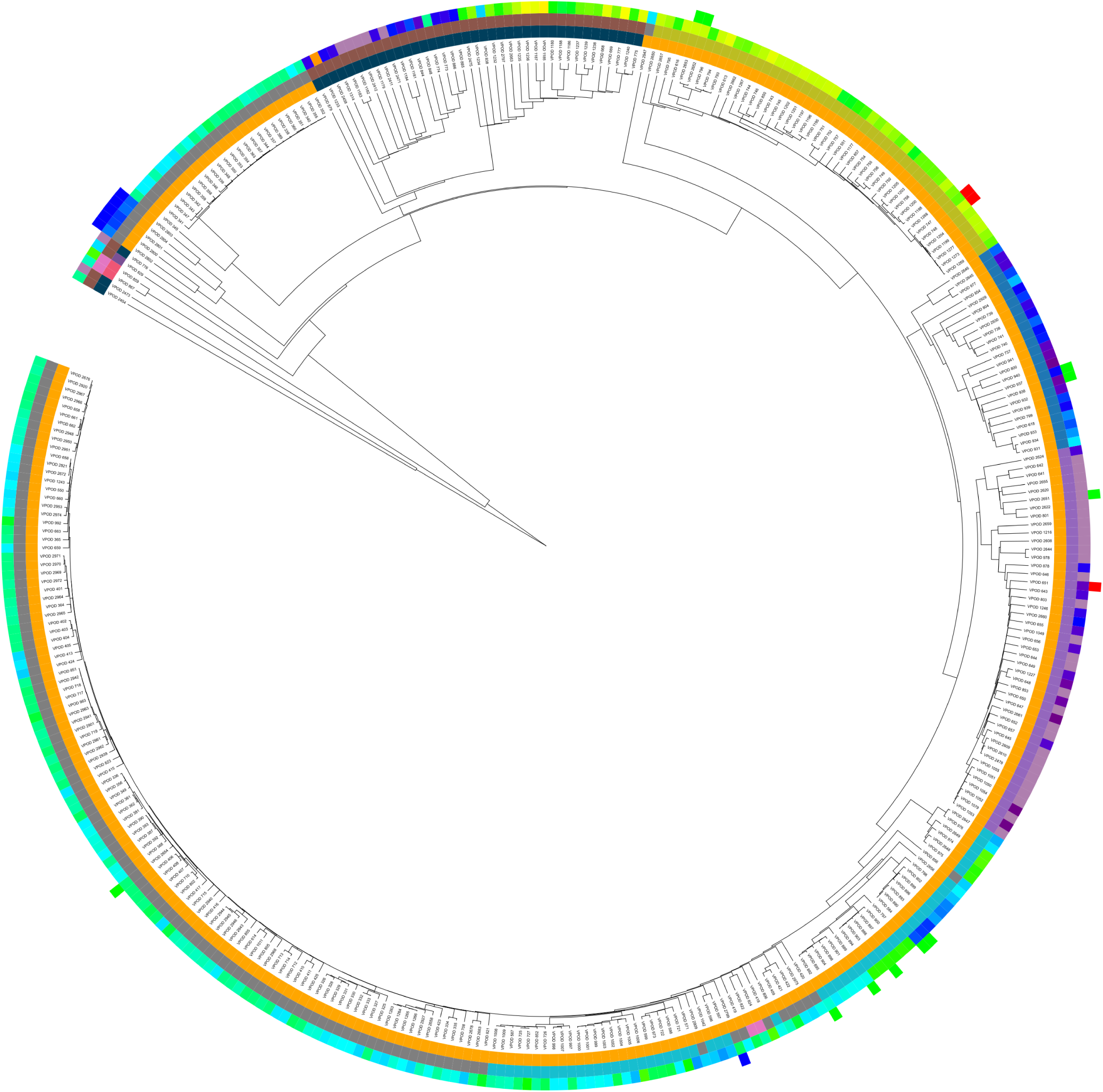
Classification of opsin sequences based on FANTASIA V2/ProstT5 metrics. Sequences are the same as in PLMView reconstruction of Fig. 5A. See legend of Fig. 5A for color coding and description of the rings. The tree was constructed using cosine distance and the single-linkage clustering method (see Methods).

## Notes

https://zenodo.org/records/20555097

https://gitlab.lcqb.upmc.fr/Vinh-Son/PLMView

### Competing Interest Statement

The authors have declared no competing interest.

## References

1. Lewin, H.A., Robinson, G.E., Kress, W.J., Baker, W.J., Coddington, J., Crandall, K.A., Durbin, R., Edwards, S.V., Forest, F., Gilbert, M.T.P. et al. (2018). Earth biogenome project: sequencing life for the future of life. Proceedings of the National Academy of Sciences 115, 4325–4333.

2. Carroll, D., Daszak, P., Wolfe, N.D., Gao, G.F., Morel, C.M., Morzaria, S., Pablos-Méndez, A., Tomori, O., and Mazet, J.A. (2018). The global virome project. Science 359, 872–874.

3. The Integrative HMP (iHMP) Research Network Consortium (2019). The integrative human microbiome project. Nature 569, 641–648.

4. Sunagawa, S., Acinas, S.G., Bork, P., Bowler, C., Eveillard, D., Gorsky, G., Guidi, L., Iudicone, D., Karsenti, E. et al. (2020). *Tara* Oceans: towards global ocean ecosystems biology. Nature Reviews Microbiology 18, 428–445.

5. Pavlopoulos, G.A., Baltoumas, F.A., Liu, S., Selvitopi, O., Camargo, A.P., Nayfach, S., Azad, A., Roux, S., Call, L., Ivanova, N.N. et al. (2023). Unraveling the functional dark matter through global metagenomics. Nature 622, 594–602.

6. Pazos, F., Rausell, A., and Valencia, A. (2006). Phylogeny-independent detection of functional residues. Bioinformatics 22, 1440–1448.

7. Rauer, C., Sen, N., Waman, V.P., Abbasian, M., and Orengo, C.A. (2021). Computational approaches to predict protein functional families and functional sites. Current Opinion in Structural Biology 70, 108–122.

8. McDonald, A.G., and Tipton, K.F. (2023). Enzyme nomenclature and classification: the state of the art. The FEBS journal 290, 2214–2231.

9. Kanehisa, M., Sato, Y., Kawashima, M., Furumichi, M., and Tanabe, M. (2016). KEGG as a reference resource for gene and protein annotation. Nucleic Acids Research 44, D457– D462.

10. Gaudet, P., and Dessimoz, C. (2016). Gene Ontology: pitfalls, biases, and remedies. In The Gene Ontology Handbook pp. 189–205.. Springer pp. 189–205.

11. Carbon, S., Douglass, E., Good, B.M., Unni, D.R., Harris, N.L., Mungall, C.J., Basu, S., Chisholm, R.L., Dodson, R.J., Hartline, E. et al. (2020). The Gene Ontology resource: enriching a GOld mine. Nucleic Acids Research 49.

12. Pitarch, B., Chagoyen, M., Ranea, J.A., and Pazos, F. (2025). A review on Gene Ontology evaluations. Database 2025, baaf058.

13. Shaw, R., Love, S.D., and McWhite, C.D. (2025). Evaluating pretrained protein language model embeddings as proxies for functional similarity. Journal of Molecular Evolution 93, 765–776.

14. Martınez-Redondo, G.I., Perez-Canales, F.M., Carbonetto, B., Fernandez, J.M., Barrios-Núñez, I., Vázquez-Valls, M., Cases, I., Rojas, A.M., and Fernández, R. (2025). FANTASIA leverages language models to decode the functional dark proteome across the animal tree of life. Communications Biology 8, 1227.

15. Senoner, T., Koludarov, I., Guenther, J., Shehu, A., Rost, B., and Bromberg, Y. (2025). Which pLM to choose? bioRxiv. doi: 10.1101/2025.10.30.685515.

16. Littmann, M., Bordin, N., Heinzinger, M., Schutze, K., Dallago, C., Orengo, C., and Rost, B. (2021). Clustering FunFams using sequence embeddings improves EC purity. Bioinformatics 37, 3449–3455.

17. Gligorijevic, V., Renfrew, P.D., Kosciolek, T., Leman, J.K., Berenberg, D., Vatanen, T., Chandler, C., Taylor, B.C., Fisk, I.M., Vlamakis, H., et al. (2021). Structure-based protein function prediction using graph convolutional networks. Nature Communications 12, 3168.

18. Yu, T., Cui, H., Li, J.C., Luo, Y., Jiang, G., and Zhao, H. (2023). Enzyme function prediction using contrastive learning. Science 379, 1358–1363.

19. Rong, D., Zhong, B., Zheng, W., Hong, L., and Liu, N. (2025). Autoregressive enzyme function prediction with multi-scale multi-modality fusion. Briefings in Bioinformatics 26, bbaf476.

20. Adams, E., Bai, L., Lee, M., Yu, Y., and AlQuraishi, M. (2025). From mechanistic interpretability to mechanistic biology: training, evaluating, and interpreting sparse autoencoders on protein language models. bioRxiv. doi: 10.1101/2025.02.06.636901.

21. Simon, E., and Zou, J. (2025). InterPLM: discovering interpretable features in protein language models via sparse autoencoders. Nature Methods 22, 2107–2117.

22. Pantolini, L., Studer, G., Pereira, J., Durairaj, J., Tauriello, G., and Schwede, T. (2024). Embedding-based alignment: combining protein language models with dynamic programming alignment to detect structural similarities in the twilight-zone. Bioinformatics 40, btad786.

23. Iovino, B.G., and Ye, Y. (2024). Protein embedding based alignment. BMC Bioinformatics 25, 85.

24. Malec, J., Golding, G.B., and Ilie, L. (2026). Protein embeddings and local alignments. Computational and Structural Biotechnology Journal 31, 24–37.

25. Chen, J., Wu, H., and Wang, N. (2024). KEGG orthology prediction of bacterial proteins using natural language processing. BMC Bioinformatics 25, 146.

26. Barrios-Núñez, I., Martínez-Redondo, G.I., Medina-Burgos, P., Cases, I., Fernández, R., and Rojas, A.M. (2024). Decoding functional proteome information in model organisms using protein language models. NAR Genomics and Bioinformatics 6, lqae078.

27. Schnoes, A.M., Ream, D.C., Thorman, A.W., Babbitt, P.C., and Friedberg, I. (2013). Biases in the experimental annotations of protein function and their effect on our understanding of protein function space. PLoS Computational Biology 9, e1003063.

28. Vicedomini, R., Bouly, J., Laine, E., Falciatore, A., and Carbone, A. (2022). Multiple profile models extract features from protein sequence data and resolve functional diversity of very different protein families. Molecular Biology and Evolution 39, msac070.

29. Pierella Karlusich, J.J., Cosnier, K., Zinger, L., Henry, N., Nef, C., Bernard, G., Scalco, E., Dvorak, E., Rocha Jimenez Vieira, F., et al. (2025). Patterns and drivers of diatom diversity and abundance in the global ocean. Nature Communications 16, 3452.

30. Lombardi, G., and Carbone, A. (2024). MuLAN: Mutation-driven Light Attention Networks for investigating protein-protein interactions from sequences. bioRxiv pp. 2024–08. doi: 10.1101/2024.08.24.609515.

31. Maeda, K., Hagglund, P., Finnie, C., Svensson, B., and Henriksen, A. (2006). Structural basis for target protein recognition by the protein disulfide reductase thioredoxin. Structure 14, 1701–1710.

32. Lemaire, S.D., Lombardi, G., Mancini, A., Carbone, A., and Henri, J. (2025). Functional recoding of chlamydomonas reinhardtii thioredoxin type-h into photosynthetic type-f by switching selectivity determinants. Frontiers in Plant Science 16, 1554272.

33. Yokoyama, S., and Radlwimmer, F.B. (1998). The” five-sites” rule and the evolution of red and green color vision in mammals. Molecular Biology and Evolution 15, 560–567.

34. Matsumoto, Y., Hiramatsu, C., Matsushita, Y., Ozawa, N., Ashino, R., Nakata, M., Kasagi, S., Di Fiore, A., Schaffner, C.M., Aureli, F., et al. (2014). Evolutionary renovation of L/M opsin polymorphism confers a fruit discrimination advantage to ateline New World monkeys. Molecular Ecology 23, 1799–1812.

35. Chi, H., Cui, Y., Rossiter, S.J., and Liu, Y. (2020). Convergent spectral shifts to blue-green vision in mammals extends the known sensitivity of vertebrate M/LWS pigments. Proceedings of the National Academy of Sciences 117, 8303–8305.

36. Asenjo, A.B., Rim, J., and Oprian, D.D. (1994). Molecular determinants of human red/green color discrimination. Neuron 12, 1131–1138.

37. Abramson, J., Adler, J., Dunger, J., Evans, R., Green, T., Pritzel, A., Ronneberger, O., Willmore, L., Ballard, A.J., Bambrick, J. et al. (2024). Accurate structure prediction of biomolecular interactions with AlphaFold3. Nature 630, 493–500.

38. Heinemann, U., and Roske, Y. (2021). Cold-shock domains—abundance, structure, properties, and nucleic-acid binding. Cancers 13, 190.

39. Zeeb, M., and Balbach, J. (2003). Single-stranded dna binding of the cold-shock protein cspb from bacillus subtilis: Nmr mapping and mutational characterization. Protein Science 12, 112–123.

40. Wang, N., Yamanaka, K., and Inouye, M. (2000). Acquisition of double-stranded dna-binding ability in a hybrid protein between escherichia coli cspa and the cold shock domain of human yb-1. Molecular microbiology 38, 526–534.

41. Ugarte, A., Vicedomini, R., Bernardes, J., and Carbone, A. (2018). A multi-source domain annotation pipeline for quantitative metagenomic and metatranscriptomic functional profiling. Microbiome 6, 149.

42. Mancini, A., Pho, V.S., Bianchi, A., Lombardi, G., Lyu, C., and Carbone, A. (2026). Scaling the profile of life by function with spin. Bioinformatics Advances 6, vbag064.

43. Vaswani, A., Shazeer, N., Parmar, N., Uszkoreit, J., Jones, L., Gomez, A.N., Kaiser, L., and Polosukhin, I. (2017). Attention is all you need. In I. Guyon, U.V. Luxburg, S. Bengio, H. Wallach, R. Fergus, S. Vishwanathan, and R. Garnett, eds. Advances in Neural Information Processing Systems vol. 30. Curran Associates, Inc.

44. Lin, Z., Akin, H., Rao, R., Hie, B., Zhu, Z., Lu, W., Smetanin, N., Verkuil, R., Kabeli, O., Shmueli, Y., dos Santos Costa, A., Fazel-Zarandi, M., Sercu, T., Candido, S., and Rives, A. (2023). Evolutionary-scale prediction of atomic-level protein structure with a language model. Science 379, 1123–1130.

45. Meier, J., Rao, R., Verkuil, R., Liu, J., Sercu, T., and Rives, A. (2021). Language models enable zero-shot prediction of the effects of mutations on protein function. Advances in Neural Information Processing Systems 34, 29287–29303.

46. Marquet, C., Heinzinger, M., Olenyi, T., Dallago, C., Erckert, K., Bernhofer, M., Nechaev, D., and Rost, B. (2022). Embeddings from protein language models predict conservation and variant effects. Human Genetics 141, 1629–1647.

47. Yeung, W., Zhou, Z., Li, S., and Kannan, N. (2023). Alignment-free estimation of sequence conservation for identifying functional sites using protein sequence embeddings. Briefings in Bioinformatics 24, bbac599.

48. Malec, J., Rusen, K., Golding, G.B., and Ilie, L. (2025). Ankh-score produces better sequence alignments than AlphaFold3. Proteins: Structure, Function, and Bioinformatics. doi: 10.1002/prot.70143.

49. Anonymous Authors (2025). SCUT: Spectral Clustering for Unsupervised Classification Trees. OpenReview. URL: https://openreview.net/forum?id=wjTysDxSg5 submission to Transactions on Machine Learning Research, Paper 6704.

50. Katoh, K., and Standley, D.M. (2013). MAFFT multiple sequence alignment software version 7: improvements in performance and usability. Molecular Biology and Evolution 30, 772– 780.

51. Katoh, K., and Standley, D.M. (2016). A simple method to control over-alignment in the MAFFT multiple sequence alignment program. Bioinformatics 32, 1933–1942.

52. Minh, B.Q., Schmidt, H.A., Chernomor, O., Schrempf, D., Woodhams, M.D., Von Haeseler, A., and Lanfear, R. (2020). IQ-TREE 2: new models and efficient methods for phylogenetic inference in the genomic era. Molecular Biology and Evolution 37, 1530–1534.

53. Kalyaanamoorthy, S., Minh, B.Q., Wong, T.K., Von Haeseler, A., and Jermiin, L.S. (2017). ModelFinder: fast model selection for accurate phylogenetic estimates. Nature Methods 14, 587–589.

54. Frazer, S.A., Baghbanzadeh, M., Rahnavard, A., Crandall, K.A., and Oakley, T.H. (2024). Discovering genotype–phenotype relationships with machine learning and the visual physiology opsin database (vpod). GigaScience 13, giae073.

55. Delmont, T.O., Gaia, M., Hinsinger, D.D., Frémont, P., Vanni, C., Fernandez-Guerra, A., Eren, A.M., Kourlaiev, A., d’Agata, L., Clayssen, Q., et al. (2022). Functional repertoire convergence of distantly related eukaryotic plankton lineages abundant in the sunlit ocean. Cell Genomics 2.

56. Camacho, C., Coulouris, G., Avagyan, V., Ma, N., Papadopoulos, J., Bealer, K., and Madden, T.L. (2009). BLAST+: architecture and applications. BMC Bioinformatics 10, 421.

57. Slater, G.S.C., and Birney, E. (2005). Automated generation of heuristics for biological sequence comparison. BMC Bioinformatics 6, 31.

58. Elnaggar, A., Essam, H., Salah-Eldin, W., Moustafa, W., Elkerdawy, M., Rochereau, C., and Rost, B. (2023). Ankh: optimized protein language model unlocks general-purpose modelling. bioRxiv. doi: 10.1101/2023.01.16.524265.

59. Paszke, A., Gross, S., Massa, F., Lerer, A., Bradbury, J., Chanan, G., Killeen, T., Lin, Z., Gimelshein, N., Antiga, L. et al. (2019). Pytorch: An imperative style, high-performance deep learning library. Advances in Neural Information Processing Systems 32.

60. Wolf, T., Debut, L., Sanh, V., Chaumond, J., Delangue, C., Moi, A., Cistac, P., Rault, T., Louf, R., Funtowicz, M. et al. (2020). Transformers: state-of-the-art natural language processing. In Proceedings of the 2020 conference on Empirical Methods in Natural Language Processing: System Demonstrations. pp. 38–45.

61. Behnel, S., Bradshaw, R., Citro, C., Dalcin, L., Seljebotn, D.S., and Smith, K. (2010). Cython: the best of both worlds. Computing in Science & Engineering 13, 31–39.

62. Nickolls, J., Buck, I., Garland, M., and Skadron, K. (2008). Scalable parallel programming with CUDA: is CUDA the parallel programming model that application developers have been waiting for? ACM Queue 6, 40–53.

63. Manavski, S.A., and Valle, G. (2008). CUDA compatible GPU cards as efficient hardware accelerators for Smith-Waterman sequence alignment. BMC Bioinformatics 9, S10.

64. Li, D., and Becchi, M. (2012). Multiple pairwise sequence alignments with the Needleman-Wunsch algorithm on GPU. In 2012 SC companion: high performance computing, networking storage and analysis. IEEE pp. 1471–1472.

65. Steinegger, M., and Söding, J. (2017). MMseqs2 enables sensitive protein sequence searching for the analysis of massive data sets. Nature Biotechnology 35, 1026–1028. URL: 10.1038/nbt.3988.

66. Roy, O., and Vetterli, M. (2007). The effective rank: a measure of effective dimensionality. In 2007 15th European signal processing conference. IEEE pp. 606–610.

67. Hornung, B.V.H., and Terrapon, N. (2023). An objective criterion to evaluate sequence-similarity networks helps in dividing the protein family sequence space. PLoS Computational Biology 19, e1010881.

